# Chromatin accessibility dynamics of neurogenic niche cells reveal defects in neural stem cell adhesion and migration during aging

**DOI:** 10.1101/2021.03.29.437585

**Authors:** Robin W. Yeo, Olivia Y. Zhou, Brian L. Zhong, Eric D. Sun, Paloma Navarro Negredo, Surag Nair, Mahfuza Sharmin, Tyson J. Ruetz, Mikaela Wilson, Anshul Kundaje, Alexander R. Dunn, Anne Brunet

## Abstract

Aging is accompanied by a deterioration in the regenerative and repair potential of stem cell niches in the brain^1–5^. However, the mechanisms underlying this decline are largely unknown. Here we characterize genome-wide chromatin accessibility in young and old neurogenic niche cells *in vivo*, revealing defects in neural stem cell (NSC) adhesion and migration during aging. Interestingly, chromatin accessibility at cell adhesion and migration genes decreases with age in quiescent NSCs but increases with age in activated (proliferative) NSCs, and this is accompanied by corresponding expression changes in these genes. We experimentally validate that quiescent and activated NSCs exhibit opposing adhesion and migration behaviors with age: quiescent NSCs become less adhesive (and more migratory) whereas activated NSCs and progeny become more adhesive (and less migratory) during aging. We also show that the ability of activated NSCs and progeny to mobilize out of the niche during *in vivo* neurogenesis diminishes during aging. Using tension sensors with single molecule resolution, we find that one of the cellular mechanisms by which aging impairs the migration of old activated NSCs and progeny involves increased force-producing adhesions. We identify inhibition of the cytoskeletal-regulating kinase ROCK^6, 7^ as a way to reduce force-producing adhesions and restore migration in old activated NSCs *in vitro*. Interestingly, inhibition of ROCK in the neurogenic niche of old mice boosts neurogenesis to the olfactory bulb *in vivo*. These results have important implications for restoring the migratory potential of NSCs and progeny and for improving neurogenesis in the aged brain.

## Main text

The adult brain contains regenerative neural stem cell (NSC) niches, with progenitors that can migrate to distal brain regions to generate new neurons and glial cells^1–5^. The regenerative potential of stem cell niches in the brain declines with age, and this is accompanied by a corresponding deterioration in aspects of sensory and cognitive function as well as repair ability^2, 8–10^. The subventricular zone (SVZ) neurogenic niche provides an excellent model system for studying NSC function and migration during aging. In this niche, quiescent NSCs line the ventricular wall^11–13^. Quiescent NSCs can activate and in turn generate progenitors and neuroblasts that migrate through the rostral migratory stream towards the olfactory bulb to produce new neurons^14^. NSC progeny also migrate to sites of injury to mitigate damage by generating new neurons^15^ and astrocytes^16^. Both the regenerative potential and repair abilities of the SVZ neurogenic region decline with age^17–21^. Previous transcriptomic studies have started to uncover defects in inflammation, signaling pathways, and cell cycle in SVZ neurogenic niches during aging^22–25^. However, the mechanisms underlying regenerative decline during aging – and how defects in migratory potential might be involved – remain largely unknown.

Epigenomic changes that affect chromatin states play an important role in the regulation of cell fate^26^ and aging^27^. So far, however, epigenomic studies of NSCs have been limited to whole tissues *in vivo*^28, 29^, developmental studies^30^, or culture systems^29, 31–33^. Importantly, age-dependent epigenomic changes in different cell types of the neurogenic niche *in vivo* remain unknown. Such changes to the chromatin landscape of NSCs could have a longer-lasting impact on progeny and reveal features of aging that were previously undetected. Identifying chromatin changes in different cell types from the neurogenic niche during aging could also help identify novel ways to reverse age-dependent defects in regeneration and repair and counter brain aging.

### Chromatin profiling of five cell types freshly isolated from the SVZ neurogenic region reveals opposing changes with age in quiescent and activated NSCs involving cell adhesion and migration pathways

To determine the impact of aging on the chromatin landscape of cells in the neurogenic niche *in vivo*, we generated chromatin accessibility profiles from 5 distinct cell types freshly isolated from the subventricular zone (SVZ) neurogenic niche of young and old mice. We aged cohorts of transgenic mice expressing green fluorescent protein driven by the human promoter for glial fibrillary acidic protein (GFAP-GFP)^34^, which, in combination with other markers (see Methods), enables the isolation of 5 different cell types by fluorescence-activated cell sorting (FACS)^35^ (Fig. 1a, Extended Data Fig. 1a). The SVZ neurogenic niches of young (3-5 months old) and old (20-24 months old) GFAP-GFP mice were micro-dissected, and 5 cell populations from this region were freshly isolated by this FACS method^35, 36^– endothelial cells, astrocytes, quiescent NSCs (qNSCs), activated NSCs (aNSCs), and neural progenitor cells (NPCs) (Fig. 1a, Extended Data Fig. 1a). We verified that FACS markers mostly did not change with age in isolated quiescent and activated NSCs (though there was a small increase in EGFR in old activated NSCs) (Extended Data Fig. 1a-d). This observation suggests that quiescent and activated NSCs isolated from young and old animals are largely similar in terms of cell identity, consistent with our previous findings^36^ (see also below for independent single cell RNA-seq validation and examination of cell identity, Extended Data Fig. 5, 6).

**Figure 1:**
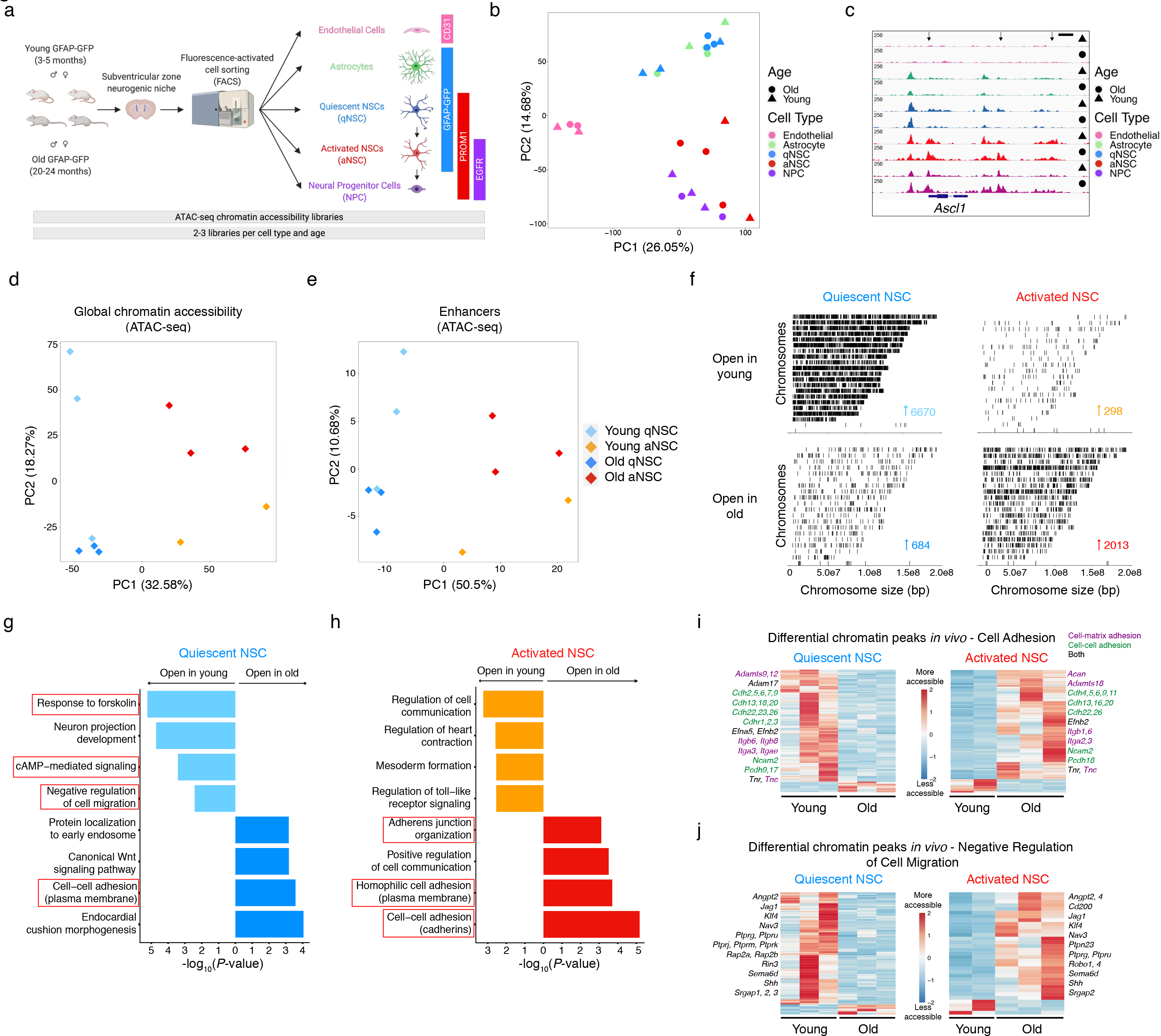
Chromatin profiling of five cell types freshly isolated from the SVZ neurogenic region reveals opposing changes with age in quiescent and activated NSCs involving cell adhesion and migration pathways. **a,** Experimental design for freshly isolating cells from the subventricular zone (SVZ) neurogenic niche in young (3-5 months) and old (20-24 months) GFAP-GFP mice. For each library, 800-2000 cells were pooled from male and female GFAP-GFP mice using fluorescence-activated cell sorted (FACS) with different markers (see Methods). qNSC, quiescent neural stem cell. aNSC, activated neural stem cell. NPC, neural progenitor cell. GFAP, glial fibrillary acidic protein. PROM1, Prominin-1. EGFR, epidermal growth factor receptor. CD31, endothelial cell marker. **b,** Principal Component Analysis (PCA) on all chromatin peaks from endothelial cells, astrocytes, quiescent NSCs (qNSC), activated NSCs (aNSC), and neural progenitor cells (NPC) freshly isolated from the SVZ of young (triangle) and old (round) GFAP-GFP mice where each dot represents a single ATAC-seq library. PCA was generated using the variance stabilizing transformation (VST)-normalized global consensus count matrix. PC, principal component. **c,** Genome browser (IGV) view of chromatin accessibility signal tracks from representative RPKM-normalized libraries of all SVZ niche cell types around the *Ascl1* loci. Black arrows represent sites of differentially accessible peaks that open in young aNSCs when compared to young qNSCs. *Ascl1*, Achaete-Scute Family bHLH Transcription Factor 1. Scale bar, 1kb. **d,** PCA on all genome-wide chromatin peaks from qNSCs and aNSCs freshly isolated from the young and old SVZ where each dot represents a single sequencing library (PC1 vs. PC2). PCA was generated using variance stabilizing transformation (VST)-normalized read counts. PC, principal component. **e,** PCA on chromatin peaks that overlap with marks of enhancers (H3K27ac and p300 binding) from qNSCs and aNSCs freshly isolated from the young and old SVZ where each dot represents a single ATAC-seq library (PC1 vs. PC2). PCA was generated using the variance stabilizing transformation (VST)-normalized NSC consensus count matrix. PC, principal component. **f,** Chromosome-level visualization of differentially accessible ATAC-seq peaks (FDR<0.05) that change with age in quiescent NSC (left) and activated NSC (right) respectively. Each vertical bar represents a dynamic ATAC-seq peak aligned to mouse chromosomes (*mm10* version of the mouse genome) that is differentially open in young (top) or old (bottom) NSCs. **g,h,** Selected GO terms for genes associated with differentially accessible ATAC-seq peaks (FDR<0.05) that change with age in quiescent **(g)** and activated **(h)** NSCs generated by EnrichR and ranked by *P*-value. ATAC-seq peaks were annotated with their nearest gene using ChIPSeeker (v1.18.0). Red boxes indicate GO terms associated with adhesion and migration. **i,** Heatmap showing accessibility levels of differential ATAC-seq peaks associated with the “Cell Adhesion” GO category (GO:0007155) that change with age in quiescent NSC (left) and activated NSC (right) respectively. Selected gene names with associated differentially accessible peaks are displayed. TMM-normalized read counts (by EdgeR), scaled row-wise. Genes names colored according to their classification on AmiGO as cell-matrix adhesion genes (purple), cell- cell adhesion genes (green), or both (black). **j,** Heatmap showing accessibility levels of differential ATAC-seq peaks associated with the “Negative Regulation of Cell Migration” GO category (GO:0030336) that change with age in quiescent NSC (left) and activated NSC (right) respectively. Selected gene names with associated differentially accessible peaks are displayed. TMM-normalized read counts (by EdgeR), scaled row-wise.

To assess chromatin accessibility genome-wide on these rare cell populations (∼100-1000 cells per individual), we used ATAC-seq^37–41^ (Fig. 1a) (see Methods). ATAC-seq libraries across all conditions exhibited stereotypical 147-bp nucleosome periodicity (Extended Data Fig. 1e) and strong enrichment of accessibility around transcription start sites (TSSs) (Extended Data Fig. 1f). Principal component analysis (PCA) (Fig. 1b) and hierarchical clustering (Extended Data Fig. 2a) on accessible chromatin peaks separated endothelial cells from other brain cells (PC1), and quiescent cells (qNSCs and astrocytes) from activated ones (aNSCs and NPCs) (PC2). The locus for *Ascl1*, a neural lineage gene involved in NSC activation^42^, shows accessible chromatin peaks in all neural cell types (but not in endothelial cells) and more accessible peaks in aNSCs and NPCs compared to qNSCs and astrocytes (Fig. 1c), consistent with *Ascl1* bulk mRNA expression^36^ (Extended Data Fig. 2b). Additionally, chromatin accessibility at genome-wide promoters positively correlated with gene expression from single cell RNA-seq data^24^ (Extended Data Fig. 2c). Thus, these genome-wide chromatin accessibility datasets represent a useful resource for studying age-related chromatin changes in 5 different cell types freshly isolated from the regenerative SVZ neurogenic niche during aging.

**Figure 2:**
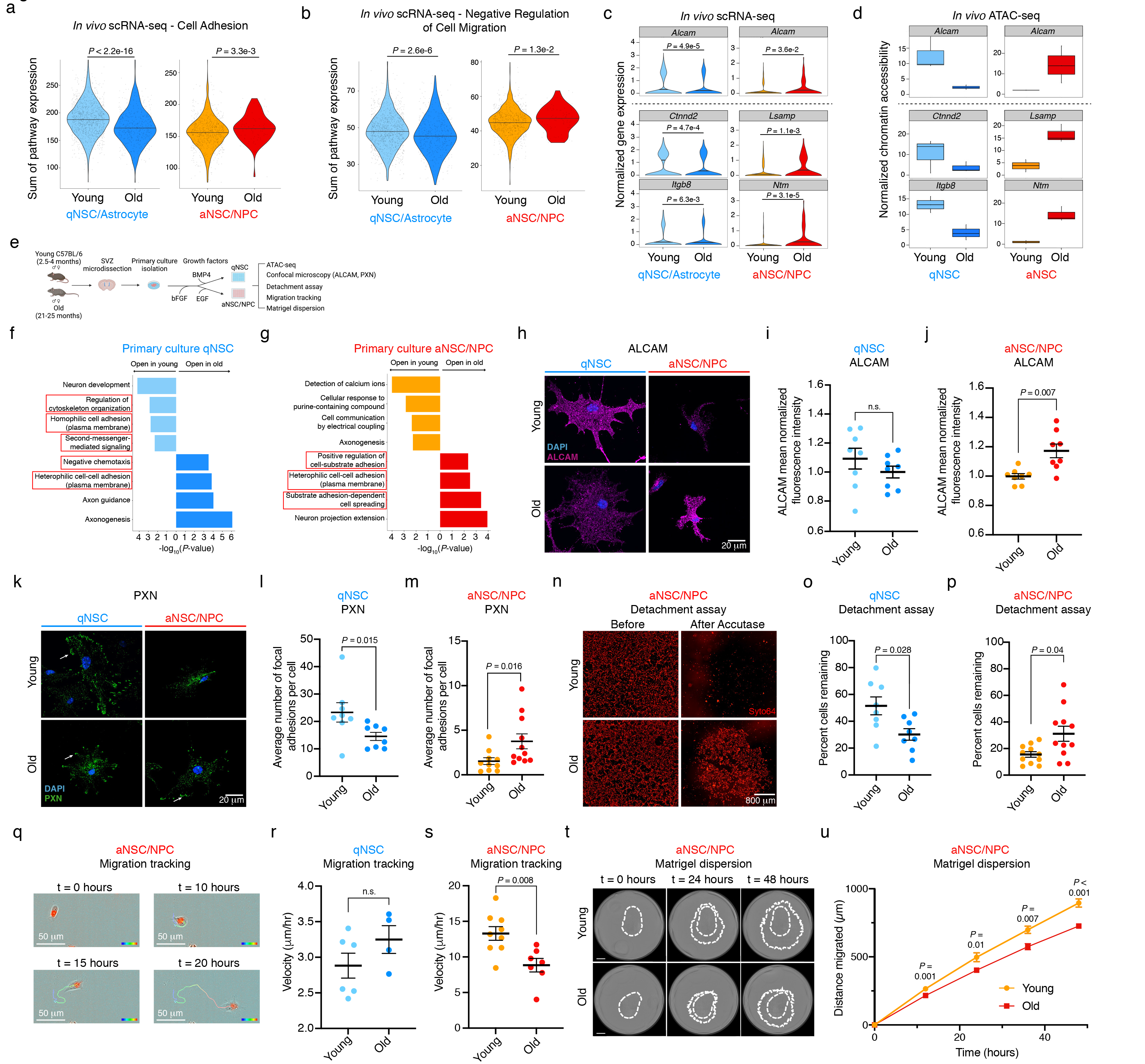
Opposing chromatin changes in quiescent and activated NSCs during aging are associated with gene expression changes and functional defects in cell adhesion and migration. **a,b,** Violin plots of the distribution of single cell cumulative expression profiles of genes within the “Cell Adhesion” GO category (GO:0007155) **(a)** and “Negative Regulation of Cell Migration” GO category (GO:0030336) **(b)** for young and old qNSCs/Astrocytes (left) and aNSCs/NPCs (right) from single cell RNA-seq data (see Methods). Each dot represents the cumulative expression of genes within the GO category in a single cell. *P*-values were calculated using a two-tailed Mann-Whitney test. **c,** Violin plots of select cell adhesion gene expression profiles that change with age for young and old qNSCs/Astrocytes (left) and aNSCs/NPCs (right) from single cell RNA-seq data (see Methods). Gene above dotted line is example of gene that is shared between qNSCs/Astrocytes and aNSCs/NPCs and genes below dotted line are examples of genes that are not shared between qNSCs/Astrocytes and aNSCs/NPCs. *P*-values were calculated using a two-tailed Mann-Whitney test. **d,** Boxplots of TMM- normalized chromatin accessibility changes in differentially accessible peaks near cell adhesion genes from (c) for young and old qNSCs/Astrocytes (left) and aNSCs/NPCs (right) freshly isolated from the SVZ. Gene above dotted line is example of gene that is shared between qNSCs/Astrocytes and aNSCs/NPCs and genes below dotted line are examples of genes that are not shared between qNSCs/Astrocytes and aNSCs/NPCs. Boxplot displays median and lower and upper quartile values. **e,** Schematic for experiments in primary cultures of qNSCs and aNSCs/NPCs. Young (2.5-4 months) and old (21-25 months) C57BL/6 SVZs were microdissected and processed for primary NSC culture. Cultures were maintained and expanded as aNSCs/NPCs with bFGF and EGF growth factors (low passage) and quiescence (for qNSCs) was induced with bFGF and BMP4 growth factors for 5-10 days for all *in vitro* experiments (ATAC-seq, confocal microscopy, detachment assay, and live-cell migration tracking). Mixed male and female mice were used for ATAC-seq experiments on primary NSC culture (similar to the animals used for the *in vivo* design, Fig. 1a). Male mice were used for other primary NSC culture experiments. **f,g,** Selected GO terms for genes associated with differentially accessible ATAC-seq peaks (FDR<0.05) that change with age in primary cultures of qNSCs **(f)** and aNSCs/NPCs **(g)** generated by EnrichR and ranked by *P*-value. Red boxes indicate GO terms associated with cell adhesion and cell migration. **h,** Representative images of immunofluorescence staining for ALCAM of young (2.5-3 months) and old (21-22.1 months) qNSCs (left) and aNSCs/NPCs (right) cultured on Matrigel. Purple, ALCAM (transmembrane glycoprotein involved in cell-cell adhesion); Blue, DAPI (nuclei). Scale bar, 20 mm. **i,j,** Quantification of ALCAM mean normalized fluorescence intensity of young (2.5-3 months) and old (21-22.1 months) qNSCs **(i)** and aNSCs/NPCs **(j)** on Matrigel. Each dot represents mean normalized fluorescence intensity of 30 fields (each containing 1-3 cells) in a primary culture derived from an individual mouse, normalized by experiment and cell size. *n* = 8 young male mice, and *n* = 8 old male mice, combined over 2 (qNSCs) or 3 (aNSCs/NPCs) independent experiments. Data are mean ± SEM. *P*-values were calculated using a two-tailed Mann-Whitney test. Data from independent experiments are in Source Data. **k,** Representative images of immunofluorescence staining for paxillin (PXN) of young (2.5-3 months) and old (21-22.1 months) qNSCs (left) and aNSCs/NPCs (right) cultured on Matrigel. Green, paxillin (PXN, focal adhesions); Blue, DAPI (nuclei). White arrow indicates localization of PXN to peripheral focal adhesions. Scale bar, 20 mm. **l,m,** Quantification of PXN immunostaining of young (2.5-3 months) and old (22.1-22.1 months) qNSCs **(l)** and aNSCs/NPCs **(m)** cultured on Matrigel. Each dot represents average number of focal adhesions per cell (28-32 cells per dot) in a primary culture derived from an individual mouse. *n* = 8 young male mice, and *n* = 8 old male mice (qNSCs), *n* = 10 young male mice, and *n* = 11 old male mice (aNSCs/NPCs), combined over 2 (qNSC) or 3 (aNSCs/NPCs) independent experiments. Data are mean ± SEM. *P*-values were calculated using a two-tailed Mann-Whitney test. Data from independent experiments are in Source Data. **n,** Representative images of young (3 months) and old (21 months) aNSCs/NPCs in the presence of cell-permeant red fluorescent nucleic acid stain Syto64 cultured on Poly-D-Lysine taken before (left) and after (right) a 5 minute incubation with Accutase and wash with 1x PBS. The Incucyte S3 system was used for 4x whole-well imaging. Incucyte S3 built-in analysis software was used for cell counts for percent cells remaining after Accutase (or Trypsin for qNSCs) treatment. Scale bar, 800 mm. **o,** Quantification of percent cells remaining of young (3-4 months) and old (21-25 months) qNSCs cultured on Poly-D-Lysine after a 15 minute incubation with trypsin and wash with 1x PBS. Each dot represents average percent cells remaining after trypsin treatment of 2-4 technical replicates (each plated at a density of 10,000 cells) per primary culture derived from an individual mouse. *n* = 8 young male mice and *n =* 8 old male mice combined over 2 independent experiments. Data are mean ± SEM. *P*-values were calculated using a two-tailed Mann-Whitney test. Data from independent experiments are in Source Data. **p,** Percent cells remaining of young (2.5-4 months) and old (21-22.1 months) aNSCs/NPCs cultured on Poly-D-Lysine after 5 minute incubation with Accutase and wash with 1x PBS. Each dot represents average percent cells remaining after Accutase treatment of 2-4 technical replicates (each plated at a density of 30,000 cells) per primary culture derived from an individual mouse. *n* = 11 young male mice, and *n* = 11 old male mice, combined over 6 experiments. Data are mean ± SEM. *P*-values were calculated using a two-tailed Mann-Whitney test. Data from independent experiments are in Source Data. **q,** Representative images of a young (3-4 months) aNSC/NPC taken at t = 0, 10, 15, and 20 hours after plating onto Poly-D-Lysine. The Incucyte system was used for 20x imaging and migration tracks were determined using Imaris (v9.3.0). Color gradient bar represents the passage of time from 0 (blue) to 20 (red) hours. Scale bar, 50 mm. **r,** Migration speed of young (3-4 months) and old (23-24 months) qNSCs on Poly-D-Lysine. Each dot represents the average velocity over a 20-hour period of 5-42 cells in a primary culture derived from an individual mouse. *n* = 6 young male mice and *n* = 4 old male mice, combined over 2 independent experiments. Data are mean ± SEM. *P*-values were calculated using a two-tailed Mann-Whitney test. Data from independent experiments are in Source Data. **s,** Migration speed of young (3-4 months) and old (21-24 months) aNSCs/NPCs on Poly-D-Lysine. Each dot represents the average velocity per mouse over a 20-hour period of 6-28 cells. *n* = 9 young male mice and *n* = 7 old male mice, combined over 3 independent experiments. Data are mean ± SEM. *P*- values were calculated using a two-tailed Mann-Whitney test. Data from independent experiments are in Source Data. **t,** Representative images of young (top) and old (bottom) aNSC/NPC dispersion through Matrigel taken 0, 24, and 48 hours after plating aNSCs/NPCs. The dotted white outline represents the outermost extent of invasion and the inner dotted white line represents the initial extent of the cells after plating (t = 0 hours). Scale bar, 800 mm. **u,** Migration distance timecourse of young (3-4 months) and old (21-24 months) aNSC/NPC dispersion through Matrigel over 48 hours with 12- hour intervals. At each timepoint, dispersion distance of aNSC cultures was averaged over 1-4 technical replicates (each seeded with 50,000 cells) from a primary culture derived from an individual mouse. *n* = 7 young male mice, and *n* = 10 old male mice, combined over 2 independent experiments. Data are mean ± SEM. *P*-values were calculated at each timepoint using a two-tailed Mann-Whitney test. Data from independent experiments are in Source Data.

Chromatin accessibility allowed separation of quiescent and activated NSCs by age (Fig. 1d for PC2, Extended Data Fig. 3a for PC3). In line with transcriptional studies^24, 25, 36^, more chromatin peaks change with age in qNSCs (7,354) than in aNSCs (2,311) (FDR < 0.05) (Extended Data Fig. 3b, Supplementary Table 2). As expected, chromatin peaks were largely located either at promoters or at intronic and distal regions, which are known to contain non-coding regulatory elements such as enhancers (Extended Data Fig. 3c). Chromatin peaks that overlap with marks of enhancers (H3K27ac and p300 binding^31^, see Methods), as well as those at distal and intronic regions, were sufficient to separate NSCs based on age whereas chromatin peaks at promoters alone were not (Fig. 1e, Extended Data Fig. 3d-g). Furthermore, chromatin peaks that dynamically change with age in qNSCs and aNSCs were almost entirely distal or intronic (Extended Data Fig. 3h, Supplementary Table 2). Thus, non-coding regulatory elements may be particularly sensitive to changes during aging.

**Figure 3:**
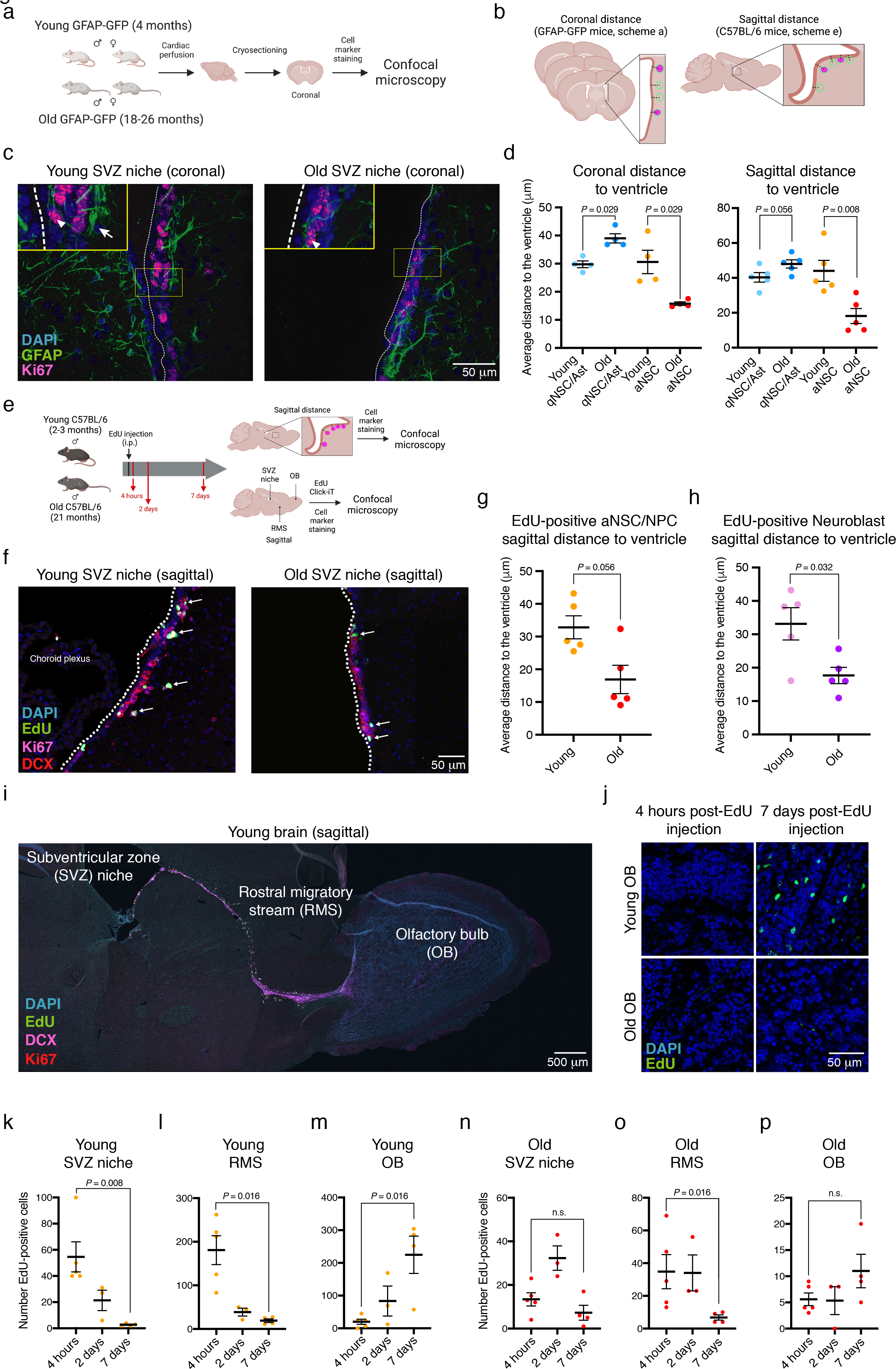
Age-dependent location defects of quiescent and activated NSCs and progeny *in vivo* in the niche, the rostral migratory stream, and the olfactory bulb. **a,** Design of immunofluorescence experiments for quantifying the location of qNSCs/astrocytes and aNSCs *in vivo* during aging in coronal brain sections. Young (4 months) and old (18-26 months) mixed-sex GFAP-GFP animals were perfused with 4% PFA, brains were cryosectioned at a thickness of 16 mm and stained for cell markers prior to quantification with confocal microscopy. **b,** Schematic depicting how coronal (a) and sagittal (e) distance of cells to the ventricle were quantified. For coronal sections, three serial coronal sections from GFAP-GFP mice were used per animal, and distance was measured from center of nuclei of cell type of interest to ventricle wall (demarcated by vinculin staining for coronal sections stained with GFAP). The corpus callosum was used as an anatomical landmark for coronal sections to ensure correspondence among sections. For sagittal sections, one sagittal section from C57BL/6 animal containing the rostral migratory stream (RMS) was used to quantify sagittal distance of cell types of interest. Only cells within 200 mm of the ventricle wall were quantified as to include only niche cells (see Methods). **c,** Representative images of immunofluorescence staining of coronal SVZ sections from young (4 months) (left) and old (25 months) (right) GFAP-GFP mice. Yellow box denotes inset containing qNSCs and niche astrocytes (GFAP+/Ki67-) and aNSC (GFAP+/Ki67+) indicated with white arrows and arrowheads, respectively. The ventricular lining is indicated by a dotted white line (see Extended Data Fig. 9a for demarcation of ventricle wall with vinculin). Green, GFAP (astrocyte/NSC marker); Pink, Ki67 (proliferation marker); Blue, DAPI (nuclei). Scale bar, 50 mm. **d,** Quantification of NSC distance to the ventricle for qNSCs and niche astrocytes (Ast) (GFAP+/Ki67-) and aNSCs (GFAP+/Ki67+) in coronal sections (left, a) of young (4 months) and old (18-26 months) SVZs from mixed sex GFAP-GFP mice and sagittal sections (right, e) of young (2-3 months) and old (21 months) SVZs from male C57BL/6 mice. Each dot represents the mean distance from the ventricle of 19 - 231 cells in 3-7 fields per section (3 sections for coronal sections or 1 section for sagittal section) per individual mouse. Coronal sections: *n* = 4 young, and *n* = 4 old mixed-sex GFAP-GFP mice, combined over 2 independent experiments. Sagittal sections: *n* = 5 young male mice, and *n* = 5 old male C57BL/6 mice combined over 2 independent experiments. Data are mean ± SEM. *P*-values were calculated using a two-tailed Mann-Whitney test. Data from independent experiments are in Source Data. **e,** Design of immunofluorescence experiments to assess both the location of quiescent and activated NSCs, NPCs, and neuroblasts in the SVZ neurogenic niche (d,f-h) and EdU-labelled NSC localization along migratory path *in vivo* (i-p). Young (2-3 months) and old (21 months) C57/BL6 male mice were intraperitoneally injected with EdU, and then sacrificed, and perfused with 4% PFA after a time interval of 4 hours, 2 days, or 7 days after EdU injection. Perfused brains were sagittally cryosectioned at a thickness of 16 mm and stained for EdU and cell markers prior to quantification with confocal microscopy. EdU, 5-Ethynyl-2’-deoxyuridine. i.p., intraperitoneal. SVZ, subventricular zone niche. RMS, rostral migratory stream. OB, olfactory bulb. **f,** Representative images of immunofluorescence staining of sagittal SVZ sections of young (2-3 months) (left) and old (21 months) (right) male C57BL/6 mice 4 hours after intraperitoneal EdU injection. Green, EdU (proliferation marker); Pink, Ki67 (aNSC/NPC/neuroblast marker); Red, DCX (neuroblast marker); Blue, DAPI (nuclei). Dotted white line indicates ventricle wall and white arrows indicate EdU-positive cells. Scale bar, 50 mm. EdU, 5-Ethynyl-2’-deoxyuridine. **g,** Distance to the ventricle was calculated for EdU+ aNSCs/NPCs (Ki67+/DCX-) in sagittal sections of young (2-3 months) and old (21 months) SVZs 4 hours after EdU injection. Each dot represents the mean aNSC/NPC distance from the ventricle of 2-35 cells from 1 section per individual mouse. *n* = 5 young male mice, and *n* = 5 old male mice, combined over 2 experiments. Data are mean ± SEM. *P*-values were calculated using a two-tailed Mann-Whitney test. Data from independent experiments are in Source Data. **h,** Distance to the ventricle was calculated for EdU+ neuroblasts (DCX+) in sagittal sections of young (2-3 months) and old (21 months) SVZs 4 hours after EdU injection. Each dot represents the mean neuroblast distance from the ventricle of 3-25 cells from 1 section per individual mouse. *n* = 5 young male mice, and *n* = 5 old male mice, combined over 2 experiments. Data are mean ± SEM. *P*-values were calculated using a two-tailed Mann-Whitney test. Data from independent experiments are in Source Data. **i,** Representative immunofluorescence staining of sagittal section of young brain encompassing the subventricular zone (SVZ) niche, rostral migratory stream (RMS), and olfactory bulb (OB) of a young (2-3 months) male C57BL/6 mouse 4 hours after intraperitoneal EdU injection. Green, EdU (proliferation marker); Red, Ki67 (aNSC/NPC/neuroblast marker); Pink, DCX (neuroblast marker); Blue, DAPI (nuclei). Scale bar, 500 mm. **j,** Representative images of immunofluorescence staining in sagittal sections of the olfactory bulb from a young (2-3 months) (top) or old (21 months) (bottom) male C57BL/6 mouse 4 hours (left) or 7 days (right) after intraperitoneal EdU injection. Green, EdU (proliferation marker); Blue, DAPI (nuclei). Scale bar, 50 mm. **k-m,** Quantification of EdU+ cells from young (2-3 months) sagittal sections 4 hours, 2 days, and 7 days post-injection of EdU. Each colored dot represents the total number of EdU+ cells counted in the SVZ (along the entire length of the ventricle **(k)**, along the entire length of the RMS **(l)**, and in the entire OB **(m)** of one sagittal section from an individual mouse. *n* = 5 young male mice 4 hours post-injection, *n* = 3 young male mice 2 days post-injection, and *n* = 4 young male mice 7 days post-injection, combined over 2 experiments. Data are mean ± SEM. *P*-values were calculated using a two-tailed Mann-Whitney test. Data from independent experiments are in Source Data. **n-p,** Quantification of EdU+ cells from old (21 months) sagittal sections 4 hours, 2 days, and 7 days post-injection of EdU. Each colored dot represents the total number of EdU+ cells counted in the SVZ (along the entire length of the ventricle **(n)**, along the entire length of the RMS **(o)**, and in the entire OB **(p)** in one sagittal section from an individual mouse. *n* = 5 old male mice 4 hours post-injection, *n* = 3 old male mice 2 days post-injection, and *n* = 4 old male mice 7 days post-injection, combined over 2 experiments. Data are mean ± SEM. *P*-values were calculated using a two-tailed Mann-Whitney test. Data from independent experiments are in Source Data.

Surprisingly, aging had opposing effects on chromatin accessibility dynamics in qNSCs and aNSCs. While most dynamic chromatin peaks in activated NSCs open with age, dynamic chromatin peaks in quiescent NSCs largely closed with age (Fig. 1f, Extended Data Fig. 3i).

Likewise, the genome-wide chromatin landscape of old aNSCs contained more accessible chromatin peaks and less nucleosomes than that of young aNSCs while the opposite was true for qNSCs (Extended Data Fig. 3j,k). These data suggest that chromatin of quiescent NSCs becomes more repressed with age while that of their activated counterparts become more permissive.

Cell adhesion and migration was a defining hallmark of genes with opposing chromatin aging changes in both quiescent and activated NSCs. Indeed, Gene Ontology (GO) biological pathway enrichment revealed that old qNSCs had decreased chromatin accessibility (open in young) at regulatory regions of genes involved in promoting cell adhesion and inhibiting cell migration (“response to forskolin”, “cAMP-mediated signaling”, “negative regulation of cell migration” (Fig. 1g, Supplementary Table 3). We also noted a “Cell-cell adhesion (plasma membrane)” signature enriched in genes with increased chromatin accessibility in old qNSCs (Fig. 1g, see Extended Data Fig. 5e for gene expression). Conversely, old aNSCs had increased chromatin accessibility (open in old) at regulatory regions of genes associated with increased cell adhesion, especially genes involved in cell-cell interactions (“cell-cell adhesion (cadherins)”, “homophilic cell adhesion (plasma membrane)”, “adherens junction organization”) and some cell-matrix interactions (“extracellular matrix assembly”) (Fig. 1h, Supplementary Table 3).

Consistently, young qNSCs and old aNSCs were grouped together along the principal component 2 (PC2) axis in PCA, based largely on cell adhesion pathways (Extended Data Fig. 4a, Supplementary Tables 4,5).

**Figure 4:**
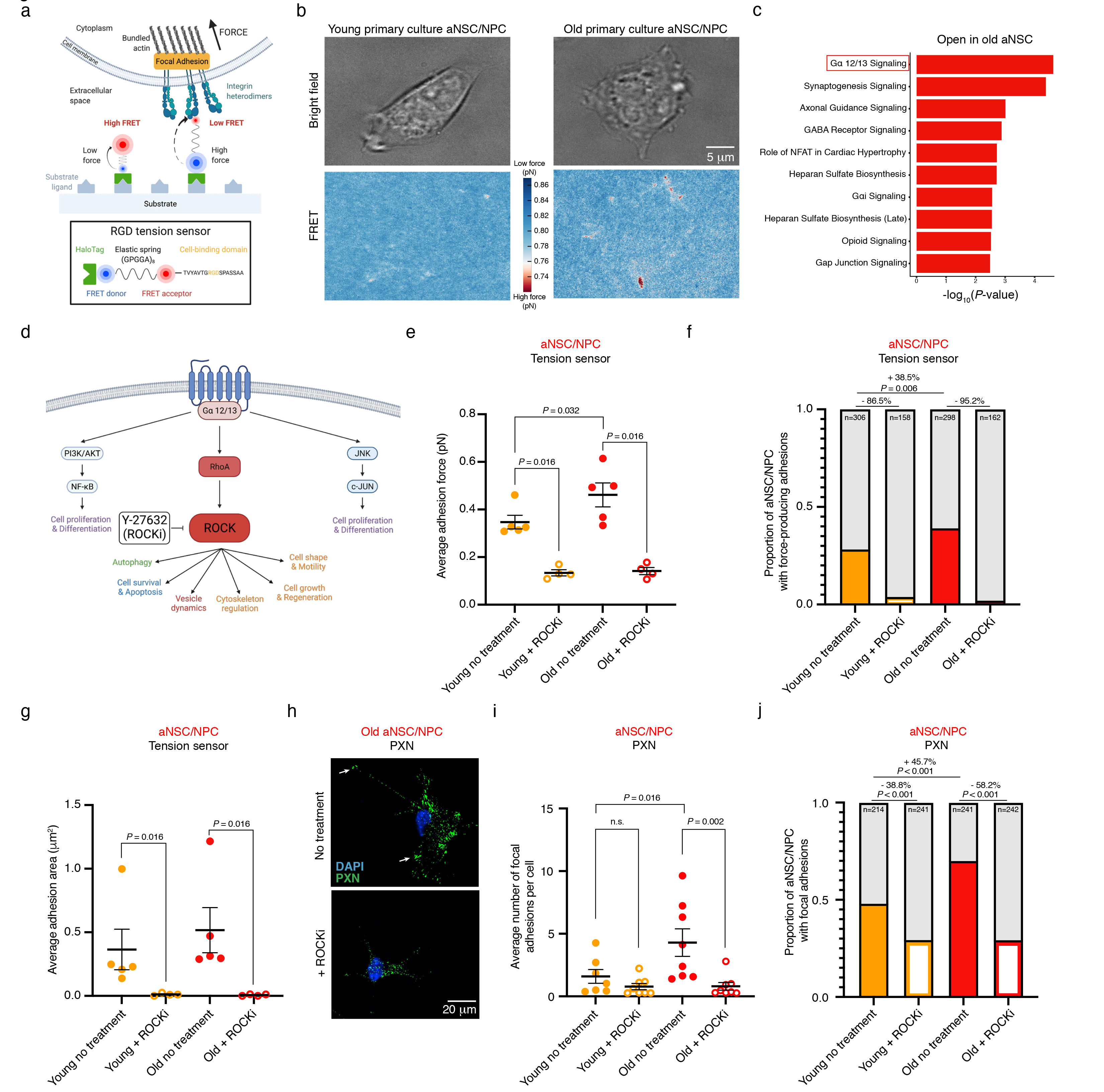
Molecular tension sensors reveal an increase in the occurrence of force-producing adhesions in old aNSCs/NPCs which can be eliminated by ROCK inhibition. **a,** Diagram of RGD molecular tension sensor (adapted from previous work, see Methods). As cells adhere to the tension sensor-coated substrate, integrin heterodimers bind to the RGD cell-binding domain and exert force across a molecular spring, resulting in decreased FRET signal. RGD, peptide containing tripeptide Arg-Gly-Asp motif. FRET, Förster resonance energy transfer. **b,** Representative image of a young (3 months) (left) and old (21 months) (right) cultured aNSCs/NPCs seeded on RGD molecular tension sensors taken with bright field illumination (top) and FRET donor/acceptor fluorescence wavelengths to generate a high-resolution traction map of FRET efficiency (bottom). The colored bar reflects the FRET efficiency, a measure of adhesion strength, where low FRET efficiency indicates high force (red color) and high FRET efficiency indicates low force (blue color). Scale bar, 5 mm. **c,** Top 10 canonical pathways enriched for genes associated with differentially accessible peaks that open with age in freshly isolated aNSCs (FDR<0.05) generated by Ingenuity Pathway Analysis (IPA) and ranked by *P*-value. ATAC-seq peaks were annotated with their nearest gene using ChIPSeeker (v1.18.0) and *P*-values were calculated using Fisher’s Exact Test by IPA. **d,** Diagram of the Ga 12/13 signaling pathway with selected targets including ROCK and downstream biological processes regulated by ROCK (adapted from the IPA canonical pathway diagram “Ga 12/13 Signaling” and previous work, see Methods). RhoA, Ras homolog family member A. ROCK, Rho- associated coiled-coiled kinase. PI3K, phosphoinositide 3-kinase. AKT, AKT serine/threonine kinase 1. NF-kB, Nuclear factor-kappa B. JNK, c-Jun N-terminal kinase. Y-27632, pharmacological inhibitor of ROCK. ROCKi, ROCK inhibitor. **e,** Quantification of average adhesion force (pN) exhibited by young (2.4-3 months) and old (20.5-21 months) cultured aNSCs/NPCs treated with H2O vehicle (no treatment, solid dots) or 10 mM ROCKi (ROCKi treatment, open circles) determined using RGD molecular tension sensors. Each dot represents the average force produced by one cell (15– 89 cells per dot) in a primary culture derived from an individual mouse. *n* = 5 young male mice and *n* = 5 old male mice (no treatment), *n =* 4 young male mice and *n* = 4 old male mice (ROCKi treatment), combined over 5 independent experiments. Data are mean ± SEM. *P*-values were calculated using a two-tailed Mann-Whitney test. Data from independent experiments are in Source Data. **f,** Proportion of young (2.4-3 months) and old (20.5-21 months) cultured aNSCs/NPCs exhibiting force-producing adhesion patterns as determined from RGD tension sensor experiments, when treated with H2O (no treatment, solid bars), or ROCKi (ROCKi treatment, open bars). Same experiment as in (e). Colored bars represent proportion of cells that have force-producing adhesions. *P*-values were calculated using a combined Fisher’s exact test. Data from independent experiments are in Source Data. **g,** Quantification of average adhesion area under force-producing adhesions (mm^2^) of young (2.4-3 months) and old (20.5-21 months) cultured aNSCs/NPCs treated with H2O vehicle (no treatment, solid dots) or 10 mM ROCKi (ROCKi treatment, open circles) determined using RGD molecular tension sensors. Each dot represents the average adhesion area of force-producing adhesions from a single cell (15 – 89 cells per dot) in a primary culture derived from an individual mouse. Same experiment as in (e). Data are mean ± SEM. *P*-values were calculated using a two-tailed Mann-Whitney test. Data from independent experiments are in Source Data. **h,** Representative immunofluorescence staining of old (22 months) cultured aNSCs/NPCs on Matrigel treated with H2O vehicle (top, no treatment) or 10 mM ROCKi (bottom, ROCKi treatment). Green, paxillin (PXN, focal adhesions); Blue, DAPI (nuclei). White arrows indicate PXN localization to focal adhesions. Scale bar, 20 mm. **i,** Quantification of average number of focal adhesions, determined using PXN staining, exhibited by young (2.5 months) and old (22.1 months) cultured aNSCs/NPCs treated with H2O vehicle (no treatment, solid dots) or 10 mM ROCKi (ROCKi treatment, open circles). Each dot represents the average number of focal adhesions per cell from a primary culture (30 cells per dot) derived from an individual mouse. *n* = 7 young male mice and *n* = 8 old male mice (no treatment) and *n* = 8 young male mice and *n =* 8 old male mice (ROCKi treatment), combined over 2 experiments. Data are mean ± SEM. *P*-values were calculated using a two-tailed Mann-Whitney test. Data from independent experiments are in Source Data. **j,** Proportion of young (2.5 months) and old (22.1 months) cultured aNSCs/NPCs exhibiting focal adhesions, determined using PXN staining, when treated with H2O vehicle (no treatment, solid bars) or 10 mM ROCKi (ROCKi treatment, open bars) exhibiting focal adhesions. Same experiment as in (i). Colored bars represent proportion of cells that have focal adhesions. *P*-values were calculated using a combined Fisher’s exact test. Data from independent experiments are in Source Data.

As for specific genes, old qNSCs freshly isolated from the brain showed reduced chromatin accessibility at regulatory regions of genes involved in cell adhesion to the extracellular matrix (e.g. *Itga3*, *Itgae, Itgb6*, *Itgb8*), in cell-cell adhesion (e.g. cadherins (*Cdh)* and protocadherins (*Pcdh*)) (Fig. 1i), and negative regulation of cell migration (e.g. *Jag1, Nav3, Ptpru*) (Fig. 1j). Conversely, old aNSCs exhibited increased chromatin accessibility at regulatory regions of genes implicated in cell adhesion to the extracellular matrix (e.g. *Itgb1*, *Itgb6, Itga2, Itga3*), in cell-cell adhesion (e.g. cadherins and *Pcdh18* (proto-cadherin 18)) (Fig. 1i), and negative regulation of cell migration (e.g. *Jag1, Nav3, Ptpru*) (Fig. 1j, Supplementary Tables 2,3). A number of cell adhesion genes enriched in young qNSCs were shared with those enriched in old aNSCs (Extended Data Fig. 4b). Thus, chromatin accessibility dynamics suggest that quiescent and activated NSCs exhibit opposing changes at cell migration and adhesion genes with age.

Other cell types in the niche also showed changes in chromatin accessibility with age, including at genes involved in adhesion pathways (Extended Data Fig. 4c-e, Supplementary Table 3). For example, endothelial cells had increased chromatin accessibility at several cell adhesion pathways with age (Extended Data Fig. 4d, Supplementary Table 3). Old neural progenitor cells (NPCs) exhibited increased accessibility at adhesion pathways with age (Extended Data Fig. 4e). Interestingly, a number of cell adhesion genes that showed age-related chromatin changes in old aNSCs and old NPCs were shared, suggesting that some of these changes may be preserved in downstream progeny (Extended Data Fig. 4f).

Collectively, these results indicate that aging has opposing effects on the global chromatin landscape of quiescent and activated NSCs (and NPC progeny), including opposing changes to chromatin accessibility in regulatory regions involved in cell adhesion and migration.

### Opposing chromatin changes in quiescent and activated NSCs during aging are associated with gene expression changes and functional defects in cell adhesion and migration

We next asked if the opposing changes to chromatin in quiescent and activated NSCs during aging are associated with expression changes in cell adhesion and migration genes and manifest in individual cells. Analysis of available single cell RNA-seq datasets from neurogenic niches of young and old mice^24, 43^ revealed that old quiescent and activated NSCs showed opposing expression changes in gene signatures involved in cell adhesion, negative regulation of migration, cell-cell adhesion, and cell-matrix adhesion (Fig. 2a,b, Extended Data Fig. 5), consistent with predictions from chromatin changes (with the exception of “cell-cell adhesion (plasma membrane)” which showed increased chromatin accessibility in old qNSCs but decreased gene expression in old qNSCs (Fig. 1g, Extended Data Fig. 5e)). Some of these gene signature changes were also preserved in neuroblasts, suggesting that they can persist in downstream progeny (Extended Data Fig. 5h, j). Specifically, old qNSCs/astrocytes showed downregulation of genes involved in cell adhesion, including adhesion to the extracellular matrix (ECM) (e.g. *Itgb8*) or to cells (e.g. *Alcam, Ctnnd2*) (Fig. 2c, Extended Data Fig. 5k). Conversely, old aNSCs/NPCs showed upregulation of genes involved in cell-cell adhesion (e.g. *Alcam, Lsamp, Ntm*) (Fig. 2c, Extended Data Fig. 5l), consistent with chromatin accessibility changes at these genetic loci (Fig. 2d). Thus, changes in gene expression of cell adhesion and migration pathways by single cell RNA-seq (which is not dependent on gating with FACS markers for cell type classification) corroborate opposing changes in chromatin states in quiescent and activated NSCs. We also noted a few genes for which there was not a one-to-one correlation between chromatin and expression (e.g. *Cadm2, Itgb1, Src* for quiescent NSCs and *Ccdc80, Epdr1, Flbn1, Megf10* for activated NSCs). In these cases, age-dependent chromatin changes may be poising these loci to regulate future gene expression in downstream progeny (e.g. neuroblasts).

**Figure 5:**
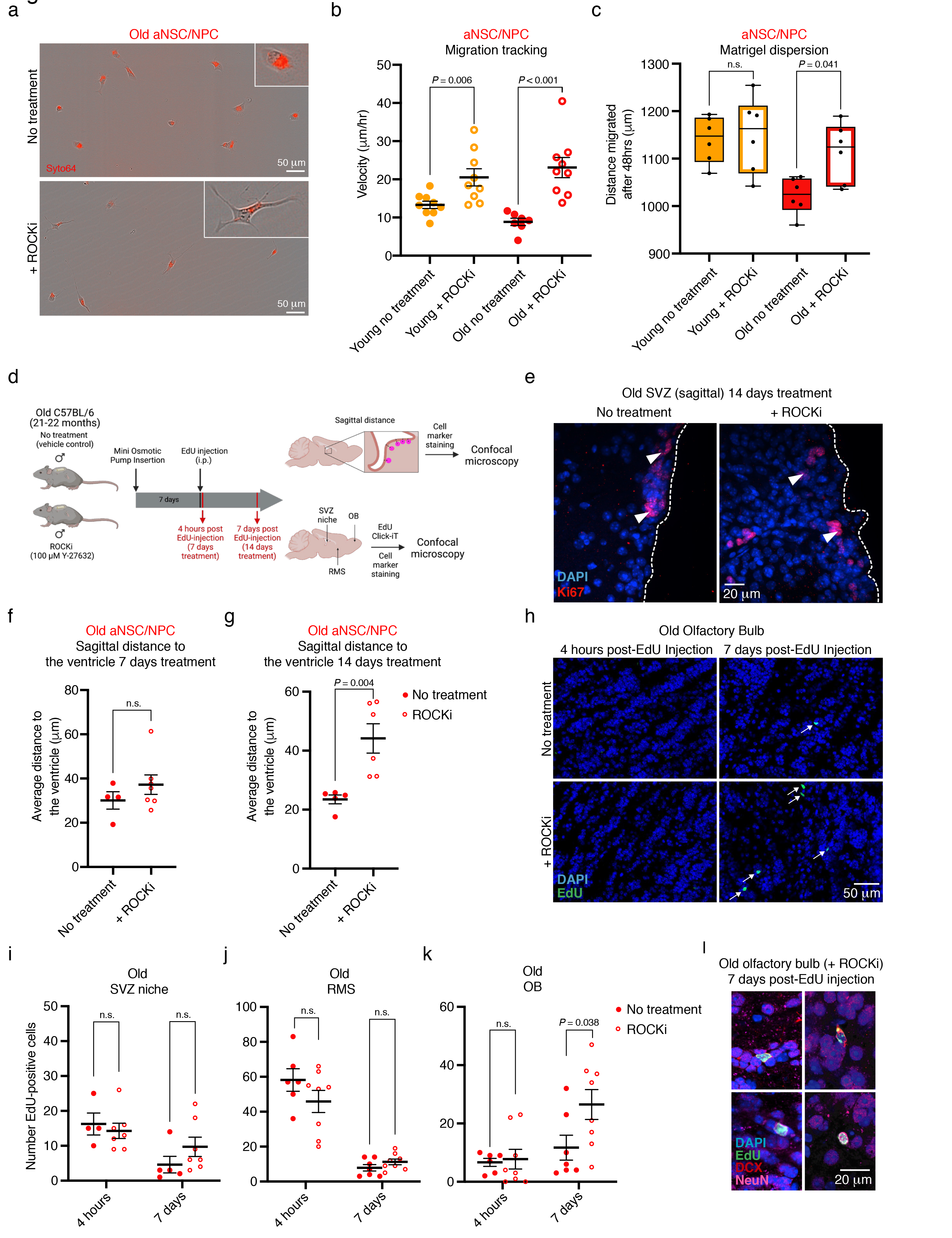
Inhibition of ROCK boosts migration speed in aNSCs/NPCs cultured from aged brains and improves *in vivo* neurogenesis in old mice. **a,** Representative images of old (21-24 months) cultured aNSCs/NPCs 12 hours after plating onto Poly-D-Lysine-coated migration plates treated with H2O vehicle (top, no treatment) or 10 mM ROCKi (bottom, ROCKi treatment). Images were taken with the Incucyte S3 system with phase contrast and red fluorescence imaging overlaid. Live cells were stained with Syto64, a cell-permeant red fluorescent nucleic acid stain, for migration tracking. Outlined inset (top-right) displays a representative magnified cell. Scale bar, 50 mm. **b,** Quantification of migration speed of young (3-4 months) and old (21-24 months) aNSCs/NPCs cultured on Poly-D-Lysine treated with H2O vehicle (no treatment, solid dots) or with 10 mM ROCKi (ROCKi treatment, open circles). Each dot represents the average velocity over a 20-hour period of cultured cells (2-28) derived from one individual mouse. *n* = 9 young male mice and *n* = 7 old male mice (no treatment) and *n* = 9 young male mice and *n* = 9 old male mice (ROCKi treatment), combined over 3 experiments. Data are mean ± SEM. *P*-values were calculated using a two-tailed Mann-Whitney test. Data from independent experiments are in Source Data. **c,** Quantification of cell dispersion through Matrigel after 48 hours by young (3-4 months) and old (21-23 months) cultured aNSCs/NPCs treated with H2O vehicle (no treatment, solid bars) or with 10 mM ROCKi (ROCKi treatment, open bars). Each dot represents the dispersion distance through Matrigel after 48 hours of cultured aNSCs/NPCs derived from an individual mouse. *n* = 6 young male mice and *n* = 6 old male mice for treated and untreated conditions, combined over 2 experiments. For each biological replicate, 1-4 technical replicates were evaluated, and dispersion distance was averaged. Boxplot displays median, and lower and upper quartile values. *P*-values were calculated using a two- tailed Mann-Whitney test. **d,** Design of *in vivo* ROCK inhibitor (ROCKi) immunofluorescence experiments to assess EdU-labelled NSC localization along migration path to the olfactory bulb. Mini- osmotic pumps with 100 mM of ROCKi or artificial cerebrospinal fluid vehicle control were stereotactically implanted into old (21-22 months) C57BL/6 male mice to continuously deliver ROCKi or vehicle control to the lateral ventricles for 7 or 14 days. After 7 days of treatment, animals were intraperitoneally injected with EdU, and then sacrificed and perfused with 4% PFA after a time interval of 4 hours (7 days treatment) or 7 days (14 days treatment). Perfused brains were sagittally cryosectioned at a thickness of 16 mm and stained for EdU and cell markers prior to quantification with confocal microscopy. EdU, 5-Ethynyl-2’-deoxyuridine. i.p., intraperitoneal. SVZ, subventricular zone niche. RMS, rostral migratory stream. OB, olfactory bulb. **e,** Representative images of immunofluorescence staining in sagittal sections of the SVZ of old (22 months) vehicle control (left, no treatment) and ROCKi-treated (right, ROCKi treatment) male mice after 14 days of treatment. Dotted line represents ventricle border. Arrowheads indicate aNSCs/NPCs (Ki67+/DCX-). Scale bar, 20 mm. **f,** Quantification of sagittal distance to the ventricle for aNSCs/NPCs (Ki67+/DCX-) in sagittal sections of old (21-22 months) SVZs treated with vehicle control (no treatment, solid dots) or ROCKi (ROCKi treatment, open circles) for 7 days (4 hours post-EdU injection). Each dot represents the mean distance from the ventricle of 18-47 cells in one sagittal section per mouse. *n* = 4 old male mice (no treatment) and *n* = 7 old male mice (ROCKi treatment), combined over 2 experiments. Data are mean ± SEM. *P*-values were calculated using a two-tailed Mann-Whitney. Data from independent experiments are in Source Data. **g,** Quantification of sagittal distance to the ventricle for aNSCs/NPCs (Ki67+/DCX-) in sagittal sections of old (21-22 months) SVZs treated with vehicle control (no treatment, solid dots) or ROCKi (ROCKi treatment, open circles) for 14 days (7 days post-EdU injection). Each dot represents the mean distance from the ventricle of 6-47 cells per individual mouse. *n* = 5 old male mice (no treatment) and *n* = 6 old male mice (ROCKi), combined over 2 experiments). Data are mean ± SEM. *P*-values were calculated using a two-tailed Mann-Whitney. Data from independent experiments are in Source Data. **h,** Representative images of immunofluorescence staining in sagittal sections of the olfactory bulb from old (21 months) male C57BL/6 mouse with treated with vehicle control (top) or ROCKi (bottom) 4 hours (left) or 7 days (right) after intraperitoneal EdU injection. White arrows indicate EdU-positive cell. Green, EdU (proliferation marker); Blue, DAPI (nuclei). Scale bar, 50 mm. EdU, 5-Ethynyl-2’-deoxyuridine. **i-k,** Quantification of EdU+ cells counted in the SVZ **(i)**, RMS **(j)**, and OB **(k)** from sagittal sections 4 hours and 7 days post-injection of EdU in old (21-22 months) mice treated with vehicle control (no treatment, solid dots) or ROCKi (ROCKi treatment, open circles). Each dot represents the number of EdU+ cells counted in one sagittal section from a single mouse. *n =* 4-7 old male mice (no treatment), and *n =* 7-8 old male mice (ROCKi treatment) for each time point (4 hours, 7 days), combined over 2 experiments. Data are mean ± SEM. *P*-values were calculated using a two-tailed Mann-Whitney test. Data from independent experiments are in Source Data. **l,** Representative immunofluorescence images of old male mouse olfactory bulb (22.4 months) 7 days after EdU injection. EdU-positive cells in the olfactory bulb are DCX+ (top) or NeuN+ (bottom). Green, EdU; Red, DCX; Pink, NeuN. Blue, DAPI (nuclei). Scale bar, 20 mm.

We used single cell RNA-seq data to examine whether changes in migration and adhesion genes occurred at the level of individual cells or were due to changes in NSC subpopulations. For most cell migration and adhesion pathways, single cell gene expression data were not bimodal (Fig. 2a,b, Extended Data Fig. 5), suggesting that changes occur in individual cells. In addition, while cell cycle analysis of single cell RNA-seq data indicated a small shift toward a subpopulation of more “quiescent-like” cells in old activated NSCs/NPCs (Extended Data Fig. 6a,b), not all age-dependent adhesion changes occurred in this subpopulation (Extended Data Fig. 6c). Thus, while subpopulation shifts in old NSCs may contribute to some of the observed changes in adhesion gene expression, age-dependent changes also occur in individual cells (see also below, immunostaining).

To experimentally test the prediction that adhesion and migration properties of NSCs change during aging, we used a culture system for quiescent NSCs and proliferative NSCs (a mix of aNSCs/NPCs)^31, 36, 44–46^ (Fig. 2e). In this culture system, cells are directly isolated from the SVZ neurogenic niches of young and old mice, and cell identity is maintained in culture with specific growth factors (Fig. 2e, see Methods). We verified that the chromatin landscape of cultured NSCs was similar to that of freshly isolated NSCs (Extended Data Fig. 7a-d, Supplementary Tables 2,3,4), notably in age-related changes to cell adhesion and migration pathways (Fig. 2f,g, Supplementary Table 3).

To examine specific adhesion proteins or pathways identified by changes in chromatin and gene expression, we performed immunostaining on young and old primary cultures of qNSCs and aNSCs/NPCs (Fig. 2e, Extended Data Fig. 7e). We first stained for ALCAM – a transmembrane glycoprotein directly involved in cell-cell adhesion^47, 48^ and indirectly implicated in cell-matrix adhesion^49^ – and whose gene exhibits opposing chromatin and gene expression changes in quiescent and activated NSCs with age (see Fig. 2c,d). Consistently, ALCAM immunostaining quantification revealed that while old qNSCs exhibited a modest (non- significant) decrease in ALCAM protein levels (Fig. 2h,i, Extended Data Fig. 7f), old aNSCs/NPCs showed a significant increase in ALCAM protein levels compared to their young counterparts (Fig. 2h,j, Extended Data Fig. 7g). We also stained for paxillin (PXN) – a marker of focal adhesions and cell-matrix adhesion^50, 51^ – given that integrin (*Itg*) genes, which regulate focal adhesions, exhibit opposing age-dependent chromatin changes in quiescent and activated NSCs with age and some of them also show gene expression changes (see Fig. 1i, Fig. 2c,d).

Paxillin immunostaining quantification revealed that consistently, old qNSCs had fewer focal adhesions than their young counterparts, whereas old aNSCs/NPCs had more focal adhesions compared to young counterparts (Fig. 2k-m, Extended Data Fig. 7h,i). These opposing changes in PXN staining were also observed within the subpopulation of cells containing at least one focal adhesion (Extended Data Fig. 7j,k), suggesting that aging induces changes at the individual cell level (though there are also subpopulation changes). These immunostaining data corroborate the opposing changes in adhesion in quiescent and activated NSCs with age predicted by chromatin and transcriptional data and indicate that at least some of the age-related changes in adhesion molecules occur at the level of individual cells.

We next functionally assessed if aging impacts cell adhesion in quiescent and activated NSCs. To probe age-related changes to cell adhesion, we adapted a detachment assay used in other cell types^52, 53^ for NSCs. In this assay, cultured NSCs are plated as a monolayer and imaged before and after enzymatic dissociation (see Methods). Old qNSCs became less adhesive (i.e. detached more easily) than their young counterparts, whereas old aNSCs/NPCs became more adhesive (i.e. detached less easily) (Fig. 2n-p). Using this detachment assay, we next tested the functional importance of a member of the NFI family of transcription factors, which regulates cell adhesion^31^ and is enriched in the accessible chromatin landscape of old aNSCs (Extended Data Fig. 8a,b). Interestingly, CRISPR/Cas9 knock-out of the NFIC transcription factor family member blunted the difference in cell adhesion between young and old aNSCs, with aNSCs/NPCs knocked out for NFIC no longer exhibiting increased adhesion with age (Extended Data Fig. 8c-g). These observations support predictions about age-related adhesion changes in NSCs and are consistent with the possibility that changes in NFIC regulation during aging are among the factors that contribute to the adhesive phenotype of old aNSCs.

To assess the migratory properties of NSCs, we performed continuous live-cell imaging of cultured NSCs over the course of 20 hours (Fig. 2q). Whereas old qNSCs were slightly more migratory than their young counterparts (non-significant) (Fig. 2r, Extended Data Fig. 7l), old aNSCs/NPCs were significantly less migratory than their young counterparts (Fig. 2s, Extended Data Fig. 7m). We also quantified the dispersion of cultured aNSCs/NPCs through Matrigel, an assay that integrates migration as well as other factors (e.g. proliferation). In this assay, aNSCs/NPCs cultured from old brains also exhibited impaired ability to disperse through extracellular matrix compared to their young counterparts (Fig. 2t,u). These dispersion differences are likely due to defects in migration (rather than proliferation) because they already manifest at early time points (12 hours, before NSCs would have time to significantly expand) and we did not detect significant proliferation differences in cultured young and old aNSCs/NPCs (Extended Data Fig. 7n). Collectively, these results indicate that aNSCs/NPCs become more adhesive and less migratory during aging.

Hence, aging has opposing effects on the adhesive and migratory properties of quiescent and activated NSCs in culture, with qNSCs becoming less adhesive/more migratory and aNSCs/NPCs becoming more adhesive/less migratory in old age.

### Age-dependent location defects of quiescent and activated NSCs and progeny *in vivo* in the niche, the rostral migratory stream, and the olfactory bulb

In the subventricular zone neurogenic niche, quiescent NSCs line the ventricles and can become activated (aNSCs) to give rise to neural progenitors (NPCs) and neuroblasts^1–4^.

Neuroblasts then migrate long distances along the rostral migratory stream (RMS) to the olfactory bulb (OB) to give rise to new neurons (neurogenesis)^3, 14, 54^. While NSC/NPC and neuroblast migration has begun to be examined in older animals^55–59^, the adhesive and migratory properties of quiescent and activated NSCs (and their progenitors) has not been systematically studied *in vivo* during aging.

Location within the subventricular zone niche is important for NSC function^60, 61^, and adhesion and migration defects in quiescent and activated NSCs during aging could manifest as changes to their niche location. We therefore assessed the location of quiescent and activated NSCs with respect to the ventricle in the SVZ neurogenic niches during aging. To this end, we performed immunostaining of brain sections – either serial coronal sections or sagittal sections that contain the rostral migratory stream – from young and old individuals. To identify specific NSC populations, we used GFAP (which marks NSCs and astrocytes), S100a6 (which marks NSCs in the adult SVZ neurogenic niche^62^), Ki67 (which marks proliferating cells), and in some cases DCX (which marks neuroblasts).

To determine the location of quiescent and activated NSCs within the SVZ neurogenic niche (and to avoid including striatal astrocytes), we quantified cells that line the ventricle within 200 μm of the ventricle border (the vast majority of cells were located within 50μm of the ventricle border) (Fig. 3a,b). For coronal sections, the ventricle border was demarcated by vinculin staining (for GFAP staining, Extended Data Fig. 9a) or ARL13b staining (for S100a6 staining, see Methods). We found that in coronal sections from old brains, GFAP+/Ki67- cells (which include qNSCs and niche astrocytes) and S100a6+/Ki67- cells (which include mostly qNSCs) were located farther away from the ventricle than in young counterparts, consistent with the possibility that quiescent NSCs (and perhaps some astrocytes) move away from their location with age (Fig. 3c,d, Extended Data Fig. 9b-d). In contrast, GFAP+/Ki67+ (which include aNSCs and proliferative niche astrocytes) and S100a6+/Ki67+ (which include mostly aNSCs) were located closer to the ventricle than in young brains (Fig. 3c,d, Extended Data Fig. 9b-d), consistent with the possibility that activated NSCs may not move as far with age. Although GFAP+/Ki67+ or S100a6+/Ki67 cells could also include repairing SVZ astrocytes that intercalate in the ependymal layer in old mice^19, 63^ or reactive astrocytes^64–68^, we observed only 4 ependymal-repairing SVZ astrocytes and no reactive astrocytes in our single cell RNA-seq dataset of 21,458 cells in both young and old animals^43^ (Extended Data Fig. 9e, markers^19, 63, 69, 70^, Supplementary Table 8, similar results with our other single cell RNA-seq dataset^24^). These data suggest that aging disrupts the location of quiescent and activated NSCs in the SVZ neurogenic niche (though we cannot exclude that some repairing or reactive astrocytes may also contribute).

We also determined the location of quiescent and activated NSCs (and progeny) in sagittal sections. We quantified distance to the ventricle for GFAP+/Ki67- (qNSCs/astrocytes), GFAP+/Ki67+ (aNSCs), DCX-/Ki67+ (aNSCs/NPCs), and DCX+/Ki67+ or DCX+/Ki67-(neuroblasts) from sagittal sections that contained the rostral migratory stream as an anatomical landmark (Fig. 3b,e). Consistent with results from coronal sections, we also observed opposing changes in location of qNSCs/astrocytes and aNSCs/NPCs with age in sagittal sections (Fig. 3f,d,g, Extended Data Fig. 9d,f). Similar to aNSCs/NPCs, neuroblasts were also located closer to the ventricles in old brain sections (Fig. 3h, Extended Data Fig. 9g). While location changes in the neurogenic niche could be due to factors other than adhesion/migration, the opposing location changes of qNSCs and aNSCs/NPCs (and neuroblasts) are consistent with *in vitro* results. This opposing directionality also suggests that location changes are unlikely to solely be due to the age-dependent thinning of the ventricle (or repairing or reactive astrocytes)^17, 19, 63, 67^.

To follow NSCs along their migratory path *in vivo*, we quantified the location of newborn NSCs and their progeny (NPCs, neuroblasts) in the SVZ niche, along the rostral migratory stream (RMS), and in the olfactory bulb (OB) during aging. To this end, we injected young and old mice with the thymidine analog 5-Ethynyl-2’-deoxyuridine (EdU), which incorporates into the DNA of replicating cells, to label and trace activated NSCs and their progeny (Fig. 3e). We verified that EdU labeling efficiency was similar in young and old individuals (Extended Data Fig. 9h,i). We then assessed the number of EdU-positive (EdU+) cells in the SVZ neurogenic niche, the migration route (RMS), and distal destination (OB), at a short (4 hours), mid- (2 days), or longer (7 days) time point after EdU injection (Fig. 3e,i-p, Extended Data Fig. 9j). In young individuals, EdU+ cells were numerous in the niche and along the RMS 4 hours after EdU injection (Fig. 3k, l, Extended Data Fig. 9j). After 7 days, young animals showed a dramatic reduction of EdU+ cells in the SVZ neurogenic niche and RMS, and a corresponding increase in EdU+ cells in the OB, consistent with mobilization of labelled cells out of the neurogenic niche and clearance from the RMS towards the OB (Fig. 3j-m, Extended Data Fig. 9j). In old individuals, EdU+ cells were less numerous overall, as expected^19–21, 71^ (Fig. 3n-p, Extended Data Fig. 9h). Interestingly, after 7 days, old animals showed no reduction of EdU+ cells in the SVZ neurogenic niche and no corresponding increase in EdU+ cells in the OB (Fig. 3n,j,p), consistent with reduced mobilization of NSCs (and progeny) out of the niche. There was some clearance of EdU+ cells through the old RMS after 7 days in old animals (Fig. 3o), suggesting that old neuroblasts retain motility in the RMS^55^ but migrate more slowly than young neuroblasts. While this experimental design integrates cell migration as well as other factors (e.g. EdU dilution, differences in cell cycle, survival, etc.), these results suggest that aging decreases the mobilization of aNSCs and progeny (NPCs, neuroblasts) out of the neurogenic niche and diminishes the migratory ability of neuroblasts to reach their distal destination *in vivo*.

### Molecular tension sensors reveal an increase in the occurrence of force-producing adhesions in old aNSCs/NPCs which can be eliminated by ROCK inhibition

We next used molecular tension sensors to assess the cellular properties underlying the increased adhesion observed in aNSCs and progenitors during aging, focusing on cell-matrix adhesion. To directly visualize the mechanical forces exerted by aNSCs/NPCs interacting with their ECM substrate, we leveraged Förster resonance energy transfer (FRET)-based molecular tension sensors (Fig. 4a)^72, 73^. Molecular tension sensors can reveal cellular forces on surfaces coated with synthetic arginine-glycine-aspartate (RGD) peptides known to bind integrins and mediate adhesion^73^ at single molecule resolution, with higher forces corresponding to a reduction in FRET efficiency (Fig. 4a,b). We verified that cells in this assay were generally healthy and did not express DCX, a marker of differentiation into neuroblasts and immature neurons (Extended Data Fig. 10a,b).

FRET measurements revealed that aNSCs/NPCs primarily exert force through discrete adhesion complexes at their periphery (Fig. 4b), consistent with PXN staining for focal adhesions (see Fig. 2k). To investigate how aging affects NSC adhesion, we quantified the adhesive force patterns for young and old cultured aNSCs/NPCs. Old aNSCs/NPCs exhibit higher average adhesion force (*P*=0.032) (Fig. 4e, Extended Data Fig. 10c), without changes in cell size (Extended Data Fig. 10d), compared to young aNSCs/NPCs. Furthermore, old aNSCs/NPCs were more likely to exhibit force-producing adhesions than young aNSCs/NPCs (Fig. 4f) (*P*=0.006, combined Fisher’s exact test). In the subset of cells that had force-producing adhesions, old aNSCs/NPCs showed a trending (non-significant) increase in average adhesion force (Extended Data Fig. 10f), consistent with increased immunostaining with paxillin (PXN), a marker of focal adhesions (see Fig. 2m, Extended Data Fig. 7k). Together, these data indicate that aging not only increases the proportion of aNSCs/NPCs that exhibit focal adhesions but also increases the average adhesion strength of individual old aNSCs/NPCs. These cellular changes with age could contribute to the adhesion/migration defects of old activated aNSCs (and progeny) in the old brain.

We next sought to identify a molecular target to counter increased adhesion strength observed in old aNSCs, as a way to restore age-related mobilization of old activated NSCs and progeny out of the niche and subsequently improve neurogenesis in the old brain. Focal adhesions are known to be regulated by several pathways, including via the Rho and Rho-associated protein kinase (ROCK) pathway^6^. Indeed, Ingenuity Pathway Analysis (IPA)^74^ showed that the most enriched signaling pathway associated with the chromatin accessibility changes in old aNSCs/NPCs was Gα 12/13 signaling (Fig. 4c, Supplementary Table 6), a pathway that regulates cell adhesion in part via ROCK^6, 75^ (Fig. 4d).

To determine if the age-related increase in cell adhesion exhibited by old aNSCs could be ameliorated via modulation of the ROCK pathway, we targeted ROCK with the small molecule inhibitor Y-27632^7, 59, 76–82^ in cultured aNSCs/NPCs. Quantification of force patterns at single-molecule resolution using RGD molecular tension sensors revealed that treating old aNSCs/NPCs with the ROCK inhibitor Y-27632 (ROCKi) eliminated force-producing adhesions and decreased adhesion area (Fig. 4e,g, Extended Data Fig.10c,e) in both young and old aNSCs/NPCs. ROCKi also decreased the proportion of cells with force-producing adhesions as assessed by tension sensors (Fig. 4f) as well as focal adhesions (marked by PXN) in young and old aNSCs/NPCs but had a greater effect on old aNSCs/NPCs (Fig. 4h-j, Extended Data Fig. 10i). In contrast, ROCKi did not affect the expression of ALCAM protein (Extended Data Fig. 10j), suggesting ROCKi acts downstream of some of the age-related adhesion gene expression changes. Together, these results indicate that ROCK inhibition eliminates force-producing adhesions and focal adhesions in aNSCs/NPCs.

### Inhibition of ROCK boosts migration speed in aNSCs/NPCs cultured from aged brains and improves *in vivo* neurogenesis in old mice

We asked if ROCK inhibition could improve the migration properties of old aNSCs *in vitro*. In cultured aNSCs/NPCs, ROCKi treatment resulted in improved migration speed in both young and old aNSCs/NPCs (Fig. 5a,b, Extended Data Fig. 10k, Supplementary Videos 1,2).

Furthermore, ROCKi rescued the age-related decline in old aNSC/NPC dispersion through Matrigel but had no effect on young aNSC/NPC dispersion (Fig. 5c), consistent with the observation that ROCKi had a more profound effect on decreasing focal adhesions in old aNSCs/NPCs than in young counterparts (Fig. 4i,j). ROCKi induced cell morphological changes^80^ (Extended Data Fig. 10a) – consistent with its ability to improve cell migration – but it did not overtly affect aNSC/NPC differentiation, proliferation, or survival under these culture conditions (Extended Data Fig. 10b,l,m). Together, these results indicate that ROCK inhibition boosts the migratory ability of aNSCs/NPCs cultured from old mice.

To determine how ROCKi treatment in the old neurogenic niche impacts the location of cells in the niche and long-distance neurogenesis *in vivo*, we delivered ROCKi (or vehicle control) via a mini-osmotic pump into the lateral ventricles of old mice, in close proximity to the SVZ neurogenic niche (Fig. 5d). Seven days after ROCKi delivery, we injected old mice with EdU and quantified EdU+ cells in the SVZ neurogenic niche, rostral migratory stream (RMS), and olfactory bulb (OB), 4 hours or 7 days after EdU-injection (corresponding, respectively, to 7 and 14 days of ROCKi treatment) (Fig. 5d). We quantified both the location of aNSCs/NPCs in the neurogenic niche in sagittal sections as well as the number of EdU+ cells in the niche and at different locations along the neurogenic migration route (Fig. 5d), as we had previously done (see Fig. 3). Interestingly, ROCKi delivery in the ventricles significantly increased aNSC/NPC distance from the ventricle 14 days after treatment (7 days post-EdU injection) in old mice (Fig. 5e-g) – a feature associated with young neurogenic niches (see Fig. 3). These results suggest that ROCK inhibition could restore at least in part the location of aNSCs/NPCs in the neurogenic niche of old animals. While ROCKi did not affect the number of EdU+ cells in the SVZ neurogenic niche or RMS (Fig. 5i,j), ROCKi delivery in the ventricles led to a significant increase in EdU+ cells in the OB of old mice 7 days post-EdU injection (Fig. 5h,k). These EdU+ cells in the OB exhibited markers of neuroblasts and neurons (DCX, NeuN) (Fig. 5l).

Collectively, these results indicate that ROCK inhibition in the neurogenic niche restores the age-related defects in location of old aNSCs/NPCs in the niche and boosts neurogenesis at a distance in old brains.

Together with our observation that ROCK inhibition improves migration of old aNSCs/NPCs in culture, these *in vivo* results are consistent with the possibility that defects in adhesion and migration of activated NSCs (and progeny) during aging could underlie age-related neurogenic decline (though other factors could also contribute). Thus, inhibition of the ROCK pathway could be a strategy to improve the migratory property of old activated NSCs (and their progeny) and to boost neurogenesis in old brains.

## Discussion

Our study identifies genome-wide changes to the global chromatin landscape of five freshly isolated cell types from the neurogenic niche of young and old animals. This analysis uncovers previously uncharacterized opposing changes in chromatin dynamics in quiescent and activated NSCs during aging *in vivo* – with some changes observed in activated NSCs preserved in downstream progeny. Interestingly, many of these opposing chromatin changes in quiescent and activated NSCs affect genes in adhesion and migration pathways and are accompanied by corresponding expression changes in these genes. Functionally, quiescent and activated NSCs exhibit opposing changes in adhesion and migration with age, with old quiescent NSCs becoming less adherent (and more migratory) and activated NSCs – and downstream progenitors – becoming more adherent (and less migratory) compared to young counterparts, and we further identify some of the adhesion molecules involved in these age-dependent changes. By assessing the force-producing ability of NSCs at the single molecule level using molecular tension sensors, we observe increased force production in old activated NSCs – a phenotype that could explain at least in part the migration defects of activated NSC with age. We also identify the ROCK pathway as a potential target to revert the defects in adhesion and migration of old activated NSCs. Indeed, a small molecule ROCK inhibitor could decrease force-production in old aNSCs and restore part of their migration ability *in vitro*. *In vivo*, ROCK inhibitor injected in the ventricles – in the vicinity of the neurogenic niche – reverses at least part of the age-dependent changes in the location of NSCs (and progeny) within the niche and boosts neurogenesis in old individuals. Our results have important implications for the role of NSC adhesion and migration during aging and point to ROCK as a potential therapeutic target to restore age-dependent defects that occur during NSC aging.

The chromatin landscape of quiescent NSCs becomes more closed with age whereas that of activated NSCs becomes more open with age. The closing of chromatin regions with age in quiescent NSCs is consistent with observations in cultured NSCs^33^, hair follicle stem cells^83^, and with findings of increased repressive chromatin marks such as H3K27me3 in quiescent stem cells from other niches^84^, including muscle satellite cells^85, 86^ and hematopoietic stem cells^87^ during aging. In contrast, the chromatin landscape of activated NSCs generally becomes more permissive with age, consistent with the observation that reducing the repressive 5-hydroxymethylcytosine (5hmC) mark in hippocampal NPCs mimics age-dependent defects^28^. Interestingly, many of these age-related chromatin changes occur in regulatory regions of cell adhesion and migration genes. The migratory properties of NSCs and neuroblasts has started to be examined with age^55–59^ and in the context of innervating distal tumors^88^. Recent live-cell imaging of SVZ wholemounts showed that aNSC and NPC migration distance and speed decreased with age^59^. However, the changes in migration and adhesion in quiescent NSCs during aging have remained largely unknown. Adhesion molecules have been shown to play important roles in both quiescent and activated stem cell function^31, 60, 61, 89–92^, and targeting some of the adhesion pathways could be beneficial to counter age-dependent functional decline. Our study reveals a dichotomy between changes to the adhesive and migratory properties of quiescent and activated NSCs with age that could be a result of both intrinsic and extrinsic changes. While our findings in cultured NSCs support the importance of intrinsic changes to cellular adhesion and migration pathways with age, extrinsic changes likely also play a key role. Indeed, age-related differences in the biomechanical properties of regenerative niches, such as stiffness, have been shown to extrinsically induce age-related phenotypes^93^. It is also possible that the mechanical memory of the *in vivo* niche could manifest in cell autonomous changes in cultured NSCs, as has been observed with cultured mesenchymal stem cells^94, 95^ and epithelial cells^96^ which retain information from past physical environments. Given that mechanical cell deformations and niche stiffness can affect chromatin states and promoter accessibility^83, 97–99^, epigenomic profiling might be especially sensitive to identifying changes in the regulation of adhesion and migration in stem cells and their progeny. Understanding the contribution of intrinsic and extrinsic responses, and how they influence each other during aging, will be critical to identify strategies to boost regeneration during aging.

Inhibiting Rho-associated kinase (ROCK) – a regulator of cytoskeletal dynamics^6^ – with the small molecule Y-27632 strikingly decreases force-producing adhesions in old activated NSCs/NPCs and improves their motility *in vitro*. Inhibiting ROCK also restores some of the age-dependent change in NSC/NPC location in the neurogenic niche and improves neurogenesis to the olfactory bulb in old mice *in vivo*. ROCK inhibition is known to impact migration in a variety of cells, including myoblasts, glioma cells, and microglia with different effects^100–107^. However, the impact of ROCK inhibitor on aNSCs/NPCs and neurogenesis remains unclear and have not been thoroughly tested during aging. Some studies indicate that ROCK inhibition causes decreased migration speed in young NSCs^59^, neuroblasts^108^, and human embryonic NSCs^109^. On the other hand, other studies show that ROCK inhibition increases NSC migration^80, 110^ and hippocampal neurogenesis^76^ in young animals. The effects of ROCK inhibition on NSC migration may depend on context and age. In the future, leveraging lineage tracing approaches to label and track stem cells as a function of space and time, coupled with *in vivo* imaging of the niche^111^, will be critical to better understand the specific effect of ROCK inhibition on neurogenesis. While ROCK has pleiotropic roles in aspects other than migration (e.g. autophagy, apoptosis, vesicle dynamics, etc.), this protein kinase may be a promising target in the aging brain to restore aspects of migration notably in cases of injury and neurodegenerative disease^112^. ROCK inhibitors are well tolerated in humans and have begun to be studied in the context neurodegenerative disease^112^ and stroke^113, 114^, and could potentially be used to ameliorate other age-dependent defects in the aging brain.

## Acknowledgements

We thank D. Wagh from the Stanford Functional Genomics Facility for Illumina NextSeq sequencing; the Stanford Shared FACS Facility for FACS use and support; K. Tsui and M. Bassik for assistance and use of the Incucyte Live Cell Imaging system; M. T. Buckley, G. A. Reeves, X. Zhao, P. P. Singh, C.-K. Hu, and K. Papsdorf for their feedback and reading of the manuscript; M. T. Buckley, G. A. Reeves, P. P. Singh, and A. McKay for independently checking scripts used in this study; J. Butterfield for help with mouse husbandry and genotyping; and M. Bassik, J. Sage, and T. Wyss-Coray for guidance. This work was supported by P01AG036695 (A.B.), R35GM130332 (A.R.D.), the Stanford Genome Training Program (R.W.Y.), a Stanford Graduate Fellowship (R.W.Y.), a Genentech Foundation Pre-doctoral Fellowship (R.W.Y.), and Stanford Medical Scientist Training Program grant T32GM007365 (O.Y.Z.). Figure schematics created using biorender.com.

## Authors contributions

R.W.Y., O.Y.Z., and A.B. planned the study. R.W.Y. performed experiments for ATAC-seq, *in vitro* migration assays, and EdU migration assays, and bioinformatically analyzed ATAC-seq and single cell-RNA seq data. O.Y.Z. performed and analyzed experiments for *in vitro* detachment assay, CRISPR-Cas9 perturbations, all immunofluorescent staining, and EdU migration assays. B.L.Z. performed the RGD tension sensor experiments and analyzed the FRET data supervised by A.R.D.. E.D.S. helped with single cell RNA-seq analyses. P.N.N. performed the mini-osmotic pump implantations for *in vivo* ROCK inhibition and helped with single cell RNA-seq. S.N. and M.S. generated the NSC deep learning model and performed motif enrichment supervised by A.K.. T.R. assisted in the *in vivo* NSC migration study. M.W. helped O.Y.Z. with staining experiments and quantification. A.K. and A.R.D. also provided intellectual contribution. R.W.Y., O.Y.Z. and A.B. wrote the manuscript, and all authors provided comments.

## Data availability

All raw sequencing reads and processed bam files for ATAC-seq libraries can be found under BioProject PRJNA715736.

## Code availability

The code used to analyze genomic data in the current study is available in the Github repository for this paper (https://github.com/sunericd/Yeo_RW_NSC_ATACseq/). Code used to process ATAC-seq data is available at https://github.com/kundajelab/atac_dnase_pipelines. Code used for deep learning model training for transcription factor binding site identification is available at https://github.com/kundajelab/retina-models.

## Methods

### Laboratory animals

For *in vivo* ATAC-seq libraries, an equal number of male and female GFAP-GFP (FVB/N background) mice^1^ were pooled and used. GFAP-GFP transgenic mice express green fluorescent protein (GFP) driven by the human promoter for glial fibrillary acidic protein (GFAP). For ATAC-seq libraries generated from cultured NSCs, the SVZs from one male and one female C57BL/6 mouse obtained from the NIA Aged Rodent colony were pooled and used. For gene knockout experiments, male and female Rosa26-Cas9 knock-in mice^2^ (C57BL/6 background) were used (https://www.jax.org/strain/024858). The Rosa26-Cas9 mice constitutively express the

Cas9 endonuclease and an EGFP reporter under the control of a CAG promoter knocked into the *Rosa26* locus. For immunohistochemistry of coronal brain sections, male and female GFAP-GFP (FVB/N background) mice^1^ were used. For all other experiments, male C57BL/6 mice obtained from the NIA Aged Rodent colony were used. In all cases, mice were habituated for more than one week at Stanford before use. At Stanford, all mice were housed in either the Comparative Medicine Pavilion or the Neuro Vivarium, and their care was monitored by the Veterinary Service Center at Stanford University under IACUC protocol 8661.

### ATAC-seq library generation from freshly isolated cells

We used fluorescence-activated cell sorting (FACS) to freshly isolate populations of endothelial cells, astrocytes, quiescent NSCs (qNSCs), activated NSCs (aNSCs), and neural progenitor cells (NPCs) from GFAP-GFP (FVB/N background) animals^1^, as previously described^3^. This GFAP- GFP strain expresses GFP under the control of the human GFAP promoter and has been used to isolate NSCs by FACS^3–6^. See also below *“Analysis of potential changes in FACS markers with age”*.

Briefly, we micro-dissected and processed the subventricular zones from young (3-5 months old) and old (20-24 months old) GFAP-GFP mice following a previously described protocol^4^ with the addition of negative gating for CD45 (hematopoietic lineage) and sorting of endothelial cells (CD31+) as described^3^ (see Extended Data Fig. 1a). All FACS sorting was performed at the Stanford FACS facility on a BD Aria II sorter, using a 100-μm nozzle at 13.1 pounds/square inch (psi), and Flowjo (v8) software was used for data analysis. Due to the rarity of NSC lineage cells, we pooled sorted cells from 2 young male and 2 young female GFAP-GFP mice for the young conditions (3-5 months old), and from 3 old male and 2-3 old female GFAP-GFP mice for the old conditions (20-24 months old). For each respective library, we sorted either 2000 astrocytes (CD45-/CD31-/GFAP-GFP+/PROM1-/EGFR-), 2000 qNSCs (CD45-/CD31-/GFAP-

GFP+/PROM1+/EGFR-) (with the exception of a single library which only had 1670 cells), 800- 1000 aNSCs (CD45-/CD31-/GFAP-GFP+/PROM1+/EGFR+), 2000 NPCs (CD45-/CD31-

/GFAP-GFP-/EGFR+), or 2000 endothelial cells (CD45-/CD31+) from GFAP-GFP animals for ATAC-seq (see Supplementary Table 1). Young and old cells of the five cell types were sorted into 150 μL of NeuroBasal-A medium (Gibco, 10888-022) with penicillin-streptomycin-glutamine (Gibco, 10378-016) and 2% B27 minus vitamin A (Gibco, 12587-010) in a 96-well V-bottomed plate (Costar, 3894) and spun down at 300g for 5 min at 4°C. Sorted cells were washed with 100 μL ice-cold 1x PBS (Corning, 21-040-CV), male and female cells were pooled by age and cell type, and then spun down at 300g for 5 min at 4°C. Next, 50 μL of lysis buffer (10 mM Tris HCl pH 7.4 (Sigma, T2194), 10 mM NaCl, 3 mM MgCl_2_ (Ambion, AM9530G), 0.10% NP-40 (Thermo, 85124)) was added to each well (alternating between young and old wells) and was immediately spun down at 500g for 10 min at 4°C. Lysis buffer was carefully aspirated and 5 μL of transposition mix (2.5 μL 2x Tagment DNA (TD) buffer, 2.25 μL nuclease-free H_2_O, 0.25 μL Tn5 transposase (Illumina, FC-121-1030)) was added to each well and pipetted 6x to resuspend nuclei. Cells were incubated for 30 min at 37°C in a sealed 96-well plate and then briefly spun down at 500g for 1 min to account for evaporation. Transposed DNA was then purified using the Zymo DNA Clean & Concentrator kit (Zymo, D4014) and eluted in 20 μL of nuclease-free H_2_O. PCR amplification and subsequent qPCR monitoring was performed as previously described in the original ATAC-seq protocol^7^. ATAC-seq libraries from young and old cells were amplified with 11-14 PCR cycles and then purified using the Zymo DNA Clean & Concentrator kit (Zymo, D4014) and eluted in 15 μL of nuclease-free H_2_O. See below “*Library sequencing and ATAC-seq quality control of in vivo and cultured NSCs*” for details on sequencing of ATAC-seq libraries.

### Analysis of potential changes in FACS markers with age

To assess whether the protein levels of the FACS markers used to freshly isolate NSCs from the SVZ neurogenic niche changed with age, we used the same sorted cell populations from young and old GFAP-GFP animals described above in “*ATAC-seq library generation from freshly isolated cells*” and used Flowjo (v10.7.1) to export the compensated scaled fluorescence values for NSC markers (GFAP-GFP, PROM1, EGFR) from all propidium iodide-negative (live) cells, quiescent NSCs (CD45-/CD31-/GFAP-GFP+/PROM1+/EGFR-), and activated NSCs (CD45-/CD31-/GFAP-GFP+/PROM1+/EGFR+) freshly isolated from the SVZ. Fluorescence values for these three channels were normalized to the young mean for an experiment and the means for each animal were then visualized using box plots with R (v3.5.2). Statistical comparisons between sample means was performed in R using a two-tailed Mann-Whitney test.

For all live cells, there was a significant increase in EGFR and PROM1 expression with age and a significant decrease in GFAP expression with age (Extended Data Fig. 1b), likely reflecting changes in cell type composition with age in the SVZ neurogenic niche^8–11^. Despite the significant decrease in GFAP expression in all live cells, we are still able to gate and FACS-purify cells with high GFAP expression (Extended Data Fig. 1a). For FACS-purified quiescent NSCs, there was no age-related change in EGFR, GFAP, or PROM1 expression (Extended Data Fig. 1c). As quiescent NSCs do not express EGFR, raw fluorescence values for this marker were very low compared to fluorescence values for other markers. For FACS-purified activated NSCs, there was no change in GFAP or PROM1 expression, but a slight increase in EGFR expression with age (Extended Data Fig. 1d).

In previous studies, we verified that young and old FACS-isolated SVZ astrocytes, qNSCs, aNSCs, and NPCs expressed well-established cell type markers (validated by RT-qPCR) and showed expected cell cycle characteristics^3^. In addition, we found that NSCs isolated using this GFAP-GFP FACS scheme in young animals were similar to NSCs isolated with a different FACS scheme using the marker GLAST^5^. We also confirmed cell adhesion and migration molecular signatures observed by ATAC-seq by single cell RNA-seq datasets (Fig. 2a-d; Extended Data Fig. 5). While we cannot completely rule out that other cells could be present in the populations isolated with this FACS scheme, especially from old mice, it is unlikely that these other cells would strongly impact the results.

### Primary NSC cell culture

To obtain primary cultures of quiescent and activated NSCs from young and old mice for ATAC-seq, we used a previously described primary culture protocol^12^, which does not depend on FACS to isolate NSCs. We micro-dissected and pooled SVZs from pairs of male and female C57BL/6 animals at a young age (3 months old) or an old age (23 months old) obtained from the NIA Aged Rodent colony. To obtain primary cultures of quiescent and activated NSCs from young and old mice for immunofluorescence staining, detachment assays, migration assays, and tension sensor assays, we used the same protocol, but cells were isolated from SVZs from a single young C57BL/6 male (2.5-4 months old) or a single old C57BL/6 male (20-25 months old) animal obtained from the NIA Aged Rodent colony (see Source Data). To obtain primary cultures for CRISPR-Cas9 experiments, SVZs from a single young male or female (3.3-5.2 months old) or old male or female (21.8 – 25.3 months old) Rosa26-Cas9^2^ (C57BL/6 background) mice were used (see Source Data). In all cases, micro-dissected SVZs were finely minced, suspended in 5 mL of 1x PBS +0.1% Gentamicin (Thermo Fisher, 15710064) and spun down at 300g for 5 min at room temperature. We then dissociated SVZs by enzymatic digestion using 5 mL of HBSS (Corning, 21-021-CVR) with 1% penicillin-streptomycin-glutamine (Gibco, 10378-016), 1 U/mL Dispase II (STEMCELL Technologies, 07913), 2.5 U/mL Papain (Worthington Biochemical, LS003126), and 250 U/mL DNAse I (D4527, Sigma-Aldrich), vortexed briefly, and left at 37°C for 40 min on a rotator. Following digestion, the samples were spun down at 300g for 5 min at room temperature and resuspended in 5 mL of NeuroBasal-A medium (Gibco, 10888-022) with 1% penicillin-streptomycin-glutamine (Gibco, 10378-016) and 2% B27 minus vitamin A (Gibco, 12587-010) and triturated repeatedly (20x) with 2-3 washes.

Single-cell suspensions were then resuspended in “complete activated media”: Neurobasal-A (Gibco, 10888-022) supplemented with 2% B27 minus vitamin A (Gibco, 12587-010), 1% penicillin–streptomycin–glutamine (Gibco, 10378-016), 20 ng/mL of EGF (Peprotech, AF-100- 15), and 20 ng/mL of bFGF (Peprotech, 100-18B). For passaging, cells were dissociated with 1 mL Accutase (STEMCELL Technologies, 07920) for 5 min at 37°C, washed once with 5 mL 1x PBS, and resuspended in “complete activated media” for expansion of activated NSCs/NPCs (aNSCs/NPCs). For quiescent NSCs (qNSCs), quiescence was induced over 5-10 days by replacing “complete activated media” with “complete quiescent media”: Neurobasal-A (Gibco, 10888-022) supplemented with 2% B27 minus vitamin A (Gibco, 12587-010), 1% penicillin–streptomycin–glutamine (Gibco, 10378-016), 50 ng/mL of BMP4 (Biolegend 94073), and 20 ng/mL of bFGF (Peprotech, 100-18B). For adherent cultures of both qNSCs and aNSCs/NPCs, we coated plates with Poly-D-Lysine (PDL) (Sigma-Aldrich, P6407, diluted 1:20 in 1x PBS) for 30-120 min at 37°C, and washed plates 4x with 1x PBS prior to plating cells at the appropriate density. All cell counting was performed using the Countess II FL Automated Cell Counter (Life Technologies, AMQAF1000).

### ATAC-seq library generation from primary NSC cultures

To establish individual primary NSC cultures for ATAC-seq, we dissected and pooled the SVZs from one male and one female C57BL/6 NIA mouse from either a young cohort (3 months old) or an old cohort (23 months old). We dissociated and cultured NSCs as described above (in the section “*Primary NSC cell culture*”) to generate 4 young and 4 old biological replicates. At passage 5, NSCs were plated at a density of 1.2 million cells per 6cm PDL-coated plate in complete quiescent media for 8 days prior to sorting. At passage 7, NSCs from the same culture were plated at a density of 1.5 million cells per 6cm plate onto PDL-coated plates in complete activated media for 24 hours prior to sorting to synchronize quiescent and activated sorting experiments. Plates were washed 3x with 1x PBS. Adherent qNSCs were lifted from the plate using 1 mL of Accutase (STEMCELL Technologies, 07920) incubated for 15 min at 37°C and adherent aNSCs/NPCs were lifted from the plate using 1 mL of Accutase (STEMCELL Technologies, 07920) incubated for 5 min at 37°C. The Accutase (STEMCELL Technologies, 07920) cell suspension was diluted with 10 mL of 1x PBS, cells were spun down at 300g for 5 min, then resuspended in 200 μL of Neurobasal-A (Gibco, 10888-022) supplemented with 2% B27 minus vitamin A (Gibco, 12587-010), 1% penicillin–streptomycin–glutamine (Gibco, 10378-016) with propidium iodide (BioLegend, 421301, 1:5000) for live/dead staining. Cells were kept on ice during all subsequent steps.

Due to concern about differing levels of dead cells in the young vs. old cultures as well as the contaminating influence of dead cells on ATAC-seq libraries, all samples were sorted using fluorescence-activated cell sorting (FACS) based on the live gate (propidium iodide).

Specifically, 10,000-15,000 live cultured qNSCs and aNSCs/NPCs (see Supplementary Table 1) were respectively sorted into 100 μL of NeuroBasal-A medium (Gibco, 10888-022) with penicillin-streptomycin-glutamine (Gibco, 10378-016) and 2% B27 minus vitamin A (Gibco, 12587-010) in a 96-well V-bottomed plate (Costar, 3894) and spun down at 300g for 5 min at 4°C. Sorted cells were washed with 100 μL ice-cold 1x PBS (Corning, 21-040-CV), and spun down at 300g for 5 min at 4°C. 50 μL of lysis buffer (10mM Tris HCl pH 7.4 (Sigma, T2194), 10mM NaCl, 3mM MgCl_2_ (Ambion, AM9530G), 0.10% NP-40 (Thermo, 85124)) was added to each well (alternating between young and old wells) which were immediately spun down at 500g for 10 min at 4°C. Lysis buffer was carefully aspirated and 50 μL of transposition mix (12.5 μL 4x Tagment DNA (TD) buffer (gift from the Chang Lab), 35 μL nuclease-free H_2_O, 2.5 μL Tn5 (gift from the Chang Lab)) was added to each well and pipetted 6x to resuspend nuclei. Cells were incubated for 30 min at 37°C in a sealed 96-well plate and then briefly spun down at 500g for 1 min to account for evaporation. Transposed DNA was then purified using the Zymo DNA Clean & Concentrator kit (Zymo, D4014) and eluted in 20 μL of nuclease-free H_2_O. PCR amplification and subsequent qPCR monitoring was performed as previously described in the original ATAC-seq protocol^7^. All libraries were amplified for 5 PCR cycles, and then an additional 4 PCR cycles (based off of qPCR amplification curves), and then purified using the Zymo DNA Clean & Concentrator kit (Zymo, D4014) and eluted in 15 μL of nuclease-free H_2_O.

### Library sequencing and ATAC-seq quality control of *in vivo* and cultured NSCs

We quantified individual library concentrations using a Bioanalyzer (High Sensitivity) and pooled at a concentration of 5 nM for sequencing. Multiplexed libraries were sequenced using NextSeq (400M) by the Stanford Functional Genomics Facility. To assess individual library quality, individual library paired-end FASTQ files were processed using the Kundaje Lab’s ATAC-seq Pipeline (https://github.com/kundajelab/atac_dnase_pipelines) with default parameters (using “-species mm10” and including “-auto_detect_adapter”).

For *in vivo* ATAC-seq libraries generated from freshly isolated SVZ cells, libraries were excluded based on insufficient read coverage (<10 million unique reads) or low peak calling (<=20,000 peaks). In general, endothelial cell libraries were of worse quality than the other four sorted cell types and we additionally censored one endothelial library with low bowtie alignment (∼92%) since all other libraries had a bowtie alignment of >= 95%. The high-quality libraries were sequenced to a mean read depth of 29,187,427 unique reads (ranging from ∼10-69 million reads per library) (see Supplementary Table 1).

In general, ATAC-seq library quality was better for cultured NSCs than freshly isolated NSCs, so we used a different set of metrics for quality control. For cultured NSCs, 1 library (out of 4) from each condition was excluded to due to poor quality, defined as the library with the lowest transcription start site (TSS) enrichment (<20 in all cases). Additionally, both young and old quiescent cultures had one library that appeared highly anomalous (∼2-fold greater fraction of reads in peaks (FRiP) and TSS enrichment compared to every other library) so they were additionally excluded to not confound results. The remaining 2-3 high-quality libraries per condition were sequenced to a mean read depth of 26,271,688 unique reads (ranging from ∼18- 42 million reads per library) (see Supplementary Table 1).

### ATAC-seq pipeline and processing

Libraries that passed quality control were re-processed using the Kundaje Lab’s ATAC-seq Pipeline (https://github.com/kundajelab/atac_dnase_pipelines) starting from de-duplicated BAM files to call peaks per multi-replicate condition. De-duplicated, Tn5-shifted tagAlign files for each replicate were converted to BAM files (using bedToBam (v2.29.2)) and sorted (using samtools sort (v1.10)) for downstream analysis. Peaks per multi-replicate condition were selected using “overlap > optimal set” resulting in approximately 20,000-90,000 peaks per *in vivo* condition (with a mean peakset size of 65,243 peaks) and approximately 90,000-150,000 peaks per cultured NSC condition (with a mean peakset size of 118,987 peaks). To generate pooled read libraries, the 2-3 high-quality filtered, de-duplicated BAM files for each condition were merged (using samtools merge (v1.10)), sorted (using samtools sort (v1.10)), Tn5-shifted (using deepTools alignmentSieve (v3.4.3)), and indexed (using samtools index (v1.10)). All analysis was performed using the *mm10* mouse genome (“TxDb.Mmusculus.UCSC.mm10.knownGene”).

### Transcription start site (TSS) enrichment

Transcription start site (TSS) enrichment heatmaps were generated with ngsplot.R (v2.6.1)^13^ using pooled, Tn5-shifted, sorted BAM files as inputs for each of the 10 conditions. All analysis was performed using the *mm10* mouse genome (“TxDb.Mmusculus.UCSC.mm10.knownGene”).

### Generating consensus peaksets and count matrices

To generate consensus peaksets for downstream analysis, BAM files and multi-replicate peak files were loaded into a Large DBA object using Diffbind (v2.10.0)^14, 15^ “dba.count” with parameters “minOverlap=0” and “score=DBA_SCORE_READS”. We annotated peaks in the consensus count matrices using the “annotatePeak” function of the package ChIPSeeker (v1.18.0)^16^ with parameters “tssREgion=c(-3000,3000)”.

### Functional enrichment of genetic elements within global peaksets

The “annotatePeak” function of ChIPSeeker (v1.18.0)^16^ was used to identify the genetic element identity of each chromatin peak within the multi-replicate peakset for each condition with parameters (tssRegion=c(-3000, 3000), annoDb=“org.Mm.eg.db”), and the annotation statistics were extracted using “@annoStat”. The different promoter terms “Promoter (<=1kb)”,

“Promoter (1-2kb)”, and “Promoter (2-3kb)” were manually grouped together under “Promoter”, and the two intron terms were manually grouped under “Intron”.

### Principal Component Analysis (PCA)

DESeq2 (v1.22.2)^17^ was used to calculate dispersion estimates from raw consensus count matrices and then variance stabilizing transformations were applied prior to visualization by principal component analysis.

For *in vivo* ATAC-seq libraries generated from freshly isolated SVZ cells, PCA on all chromatin peaks was generated using the global consensus peakset of 141,970 peaks. PCA consisting of all young and old qNSC and aNSC libraries was generated using the count matrix consisting of these 87,796 peaks. Based on peak annotations, the NSC consensus peakset was sub-divided into a distal+intronic peakset (60,231 peaks), a distal peakset (31,660 peaks), an intronic peakset (28,571 peaks), and a promoter peakset (20,633 peaks)). For ATAC-seq libraries generated from cultured NSCs, PCA was generated from the consensus peakset with 121,497 peaks. To compare how cultured NSCs compared to NSCs freshly isolated from the SVZ, we performed PCA on the count matrix consisting of the 11 freshly isolated NSC libraries and the 10 cultured NSC libraries (156,963 peaks).

### PCA with ATAC-seq peaks with enhancer marks

To identify ATAC-seq peaks that have enhancer marks, we downloaded FASTQ files for chromatin immunoprecipitation followed by sequencing (ChIP-seq) datasets for histone H3 acetylated lysine 27 (H3K27ac) and p300 – two marks of active enhancers – obtained from

cultured quiescent NSCs and proliferative (activated) NSCs^12^. We then processed these datasets using the standard ENCODE ChIP-seq pipelines. ATAC-seq peaksets that overlap with both H3K27ac and p300 marks were generated for qNSCs and aNSCs using “bedtools intersect -wa - u” resulting in subsetted peaksets of size 5401 and 3875 respectively. The resulting peak files were used to generate an accessibility count matrix (6644 peaks) which was used for PCA as described above.

### Clustering ATAC-seq libraries from freshly isolated cells for heatmap visualization

To cluster and visualize all ATAC-seq libraries from freshly isolated SVZ cell populations together, we generated a heatmap from the global consensus peakset (141,970 peaks) with “cor()” using the default Pearson’s correlation with the R library “pheatmap” (v1.0.12).

### Correlating ATAC-seq promoter accessibility and single cell RNA-seq expression

For the five cell populations freshly isolated from the SVZ of young and old mice for ATAC-seq, the average (VST-normalized) chromatin accessibility value of each gene’s promoter peak was associated with the average single cell RNA-seq^11^ log-normalized expression value for that gene. Promoters were binned in deciles based on promoter accessibility in ATAC-seq, and the association between promoter chromatin accessibility and associated gene expression was plotted as boxplots of deciles using R (v3.5.2).

### Chromatin signal track visualization for freshly isolated NSCs

Alignment tracks were visualized using IGV (v2.4.19). For each condition, the BAM file for a single representative library (see Supplementary Table 1) was normalized by Reads per Kilobase

per Million mapped reads (RPKM-normalization) and converted to a bigwig file using deepTools (v3.4.3) with the following parameters: “--extendReads 100 --normalizeUsing RPKM --binSize 10”.

### Differential peak calling with Diffbind v2

To identify differentially accessible peaks that change with age for each cell type, count matrices consisting of young and old replicates within a single cell type were generated using Diffbind (v2.10.0)^14, 15^ as described above (see “*Generating consensus peaksets and count matrices*”).

Differential peak calling was accomplished using EdgeR (v3.24.3)^18, 19^ with the following parameters for “dba.analyze”: bCorPlot=FALSE, bParallel=TRUE, bTagwise=FALSE, bFullLibrarySize=TRUE, bReduceObjects=FALSE,method=DBA_EDGER”. For all comparisons, differential peaks were obtained using a false discovery rate (FDR) threshold of 0.05 (see Supplementary Table 2). Differential peaks were annotated and associated with their closest gene using the “annotatePeak” function of the package ChIPSeeker (v1.18.0)^16^ with parameters “tssREgion=c(-3000,3000)”. For the differential peaks that change with age in the freshly isolated qNSC and aNSC libraries respectively, differential peaks were aligned to the *mm10* chromosomes using the “covplot()” function in ChIPSeeker (v1.18.0)^16^ for ease of visualization.

During the course of this study, the Diffbind package was significantly updated leading to changes in differential peak calling (Diffbind v3). Using EdgeR with Diffbind v3, similar differential peaksets to those called by Diffbind v2 can be obtained with the settings “DBA$config$design <- FALSE”. We also tested another differential peak caller using Diffbind v3 (DESeq2), which could not call differential peaks in our samples at FDR<0.05, possibly due to the low cell number used as input and the resulting relatively shallow sequencing depth. Due to this discrepancy, we verified that the original differential peaksets had clean signal pileups (Extended Data Fig. 3i, see “*Global signal pileup analysis of chromatin accessibility in NSCs*” below) and that the FDR values of the original differential peaksets (called by EdgeR using Diffbind v2) correlated well with the *P*-values of DESeq2 peaks (using Diffbind v3) (Extended Data Fig. 3l,m).

### Functional enrichment of genetic elements within differential NSC peaksets

The differentially accessible peaks that change with age in the freshly isolated qNSC and aNSC conditions, respectively, were separated into sets that close with age or open with age and were annotated with the “annotatePeak” function of ChIPSeeker (v1.18.0)^16^ with parameters (tssRegion=c(-3000, 3000), annoDb=“org.Mm.eg.db”). The annotation statistics from “@annoStat” were manually grouped into 4 categories: “Distal Intergenic”, “Intron”, “Promoter”, and “Other”.

### Nucleosome peak calling

To identify whether the chromatin landscapes of young and old freshly isolated qNSCs and aNSCs had different levels of heterochromatin-associated nucleosomes, we used the package NucleoATAC (v0.2.1)^20^. We called nucleosome peaks from our ATAC-seq data using “nucleoatac run” with parameters: “--bed” the multi-replicate peak files for each condition, “-- bam” pooled, down-sampled (to 30 million unique reads), Tn5-shifted BAM files, and “--fasta”

the mm10 Mus musculus UCSC genome. The number of nucleosome peaks for each condition were taken from the “*.nucmap_combined.bed” files and plotted with R (v3.5.2).

### Global signal pileup analysis of chromatin accessibility in NSCs

We plotted the smoothed signal using the “fc.signal.bigwig” tracks outputted by the ENCODE ATAC-seq pipeline for each of the tracks within differential peaks output by EdgeR for young and old aNSCs and qNSCs and subsampled common peaks for aNSC and qNSC separately.

### Gene Ontology (GO) Biological Pathway Enrichment of differentially accessible chromatin peaks

Differentially accessible peaks were associated with nearby genes using the ChIPSeeker (v1.18.0)^16^ function “annotatePeak”. Gene lists were uploaded to the online tool EnrichR^21, 22^ and top ranked GO terms from “GO Biological Process 2018” were extracted for pathways that change with age (see Supplementary Table 3). Selected GO terms were ranked by *P*-value and plotted for visualization in R (v3.5.2).

In most cases, there was a direct correlation between chromatin changes and transcriptional (transcriptional upregulation of genes in a GO term found to enriched in differentially accessible chromatin peaks with age). Exceptions to this include the GO term “Cell-cell adhesion (plasma membrane)” (Fig. 1g) which opened in old qNSCs but is transcriptionally downregulated with age (Extended Data Fig. 5e), “Adherens Junction Organization” which opened in old aNSCs but showed no significant transcriptional change with age (Extended Data Fig. 5b), and “Response to Forskolin” which opened in young qNSCs but showed no significant transcriptional change with age (Extended Data Fig. 5c). For GO enrichment of differentially accessible peaks derived from cultured NSCs (Fig. 2f), the GO term “Heterophilic cell-cell adhesion (plasma membrane)” whose chromatin opened in old cultured qNSCs is transcriptionally downregulated with age and the GO term “Negative Chemotaxis” which opened in old qNSCs does not show a significant transcriptional change with age.

### Gene Ontology (GO) Biological Pathway Enrichment of genes driving PC axes

To identify what biological processes were associated with peaks driving the principal components (PCs) in the PCA of young and old freshly isolated qNSCs and aNSCs, the top 1000 peaks driving the principal components (either negatively or positively) were extracted and associated with genes (see Supplementary Table 4). The genes (under header “Symbol”) associated with these peaks were uploaded to EnrichR to identify top ranked GO terms from “GO Biological Process 2018” (see Supplementary Table 5). The top 6 GO terms for PC2 negative peaks (grouping old qNSCs and young aNSCs) and PC2 positive peaks (grouping young qNSCs and old aNSCs) were ranked by *P*-value and plotted in R (v3.5.2) for visualization.

### Chromatin peak heatmaps within cell adhesion pathways

To visualize how aging affects chromatin peak accessibility associated with cell adhesion and migration, genes associated with differential peaks that change with age in either freshly isolated qNSCs or aNSCs respectively were intersected with the “Cell Adhesion” GO gene list (GO:0007155) or the “Negative Regulation of Cell Migration” GO gene list (GO:0030336) (http://www.informatics.jax.org). Differential peak accessibility levels were plotted as heatmaps using “pheatmap” (v1.0.12) in R (v3.5.2) (TMM-normalized read counts, scaled row-wise).

### Venn diagrams of cell adhesion genes with dynamically accessible chromatin peaks

To visualize whether the opposing chromatin changes during aging in qNSCs and aNSCs involved shared adhesion genes, we used Venn diagrams to visualize the number of cell adhesion genes (from the “Cell Adhesion” GO gene list (GO:0007155)) with nearby chromatin peaks (annotated by ChIPSeeker (v1.18.0)^16^) that were differentially open in young qNSCs (compared to old qNSCs) and open in old aNSCs (compared young aNSCs). We also used a Venn diagram to visualize the overlap of cell adhesion genes with nearby chromatin peaks that open with age in both aNSCs and NPCs.

### *De novo* transcription factor binding patterns using a deep learning model and TF- MoDISco

To identify transcription factor binding patterns, ATAC-seq peaks for *in vivo* quiescent or activated NSCs isolated from young and old mice were separately converted to BigWig tracks of base-resolution Tn5 insertion sites with an +4/-4 shift to account for Tn5 shift. For each cell type, in addition to the peak regions, we selected an equal number of non-peak regions that were matched for GC content in their peaks. We then trained cell type-specific BPNet models to predict the log counts and base-resolution Tn5 insertion profiles as previously reported^23, 24^.

Briefly, the BPNet model takes as input a 2,114 bp one-hot encoded input sequence and predicts the ATAC-seq profile and log counts in a 1,000 bp window centered at the input sequence.

Following BPNet formulation, we used a multinomial negative log likelihood (MNLL) for the profile output of the model and a mean square error (MSE) loss for the log counts output of the model. The relative loss weight used for the counts loss was 0.1 times the mean total counts per region. During each epoch, training examples were jittered by up to 500 bp on either side and a random half of the sequences were reverse complemented. Each batch contained a 10:1 ratio of peaks to non-peak regions. Model training was performed using Keras/Tensorflow 2. Code used for model training is available at https://github.com/kundajelab/retina-models.

We computed importance scores for the counts output using the DeepSHAP^25^ implementation of DeepLIFT algorithm^26^. We next ran the TF-MoDISco algorithm^27^ to perform *de novo* motif discovery in each cell type. We mapped the discovered motifs to position weight matrices (PWMs) from HOCOMOCO^28^ using TomTom^29^ (Extended Data Fig. 8a). For each cell type, we computed the relative fraction of TF-MoDISco seqlets attributed to each motif (Extended Data Fig. 8a).

### Single cell RNA-seq expression values for cell adhesion and migration pathways

Single cell RNA-seq datasets from young and old SVZ neurogenic niches^11^, consisting of 14,685 cells (8,884 cells from young and 5,801 cells from old), were used to identify how gene expression of specific molecular signatures and genes involved in cell adhesion and migration differs by cell type and age at single-cell resolution.

Cell types were clustered and annotated as described previously^11^ with Seurat^30^. In brief, we performed t-SNE clustering using Seurat^30^ with the first 15 principal components and performed principal component analysis (PCA) on the 4,125 most variable genes. Significant clusters and

marker genes for each significant cluster were found using the Seurat functions FindClusters() and FindAllMarkers(). This identified 11 different cell types (qNSCs/astrocytes – which cluster together, aNSCs/NPCs – which cluster together, neuroblasts, neurons, oligodendrocytes, progenitor cells, oligodendrocytes, endothelial cells, mural cells, microglia, macrophages, and T-cells) which were annotated using a combination of marker genes identified from literature and gene ontology for cell types using EnrichR (http://amp.pharm.mssm.edu/Enrichr/). Single cell gene expression values for three cell types – qNSCs/astrocytes (which cluster together), aNSCs/NPCs (which cluster together), and neuroblasts – were extracted and subset based on genes from Gene Ontology lists of cell adhesion and migration pathways. For each single cell within a cell population, the expression levels of cell adhesion/migration genes were summed. The cumulative expression level of adhesion genes comparing young qNSCs/astrocytes, aNSCs/NPCs, and neuroblasts was visualized using violin plots with R (v3.5.2). Statistical comparisons between conditions was performed in R using a two-tailed Mann-Whitney test. We also performed a Welch-corrected t-test and results were similar. A two-tailed Mann-Whitney test was chosen for reporting because it does not require data to be normally distributed and is commonly used to make comparisons for single gene expression values^31, 32^.

### Differentially accessible chromatin peaks of specific cell adhesion genes that change with age

For specific “Cell Adhesion” genes (GO:007155) (*Alcam, Ctnnd2, Itgb8, Lsamp, Ntm*), TMM-normalized accessibility values for differential peaks that change with age were obtained from the output of EdgeR (Diffbind v2) (Supplementary Table 2) and visualized using boxplots with R (v4.1.0). When there were multiple chromatin peaks associated with the same gene that change with age, the peak upstream of, and closest to the TSS was chosen.

### Single cell RNA-seq expression values for specific cell adhesion genes

Using the same single cell RNA-seq dataset as above^11^, we used the FindMarkers function from Seurat (v4.0.5) and tested if specific “Cell Adhesion” genes (GO:007155) (*Alcam, Ctnnd2, Itgb8, Lsamp, Ntm*) whose chromatin changed with age in qNSCs and aNSCs (see “*Differentially accessible chromatin peaks of specific cell adhesion genes that change with age*”) also showed transcriptional changes. Statistical comparisons between conditions was performed in R using a two-tailed Mann-Whitney test. We validated these differentially expressed genes in two other single cell RNA-seq datasets of SVZ neurogenic niches from mice at different ages^33, 34^. We picked three representative genes that were significantly downregulated with age in qNSCs/astrocytes in at least two of the three single cell RNA-seq datasets and three representative genes that were significantly upregulated with age in aNSCs/NPCs in at least two of the three single cell RNA-seq datasets. Expression values were log-normalized counts per 10,000 transcripts and were visualized using violin plots with R (v3.5.2). Data shown is from same dataset^11^ described above.

### Gene expression trajectory of specific cell adhesion genes during aging time course

Gene expression trajectories as a function of age were derived from single cell gene expression data of the SVZ of 28 mice, tiling ages from young to old (3.3 months to 29 months)^33^.

Expression values were log-normalized counts per 10,000 transcripts. Each dot represents gene expression value per mouse. Shaded region corresponds to 95% confidence interval. Line is a smoothed fit using geom_smooth with method=”loess” (ggplot2 v3.3.5).

### Assessing cell cycle heterogeneity and cell adhesion pathways using single cell RNA-seq

Using single cell RNA-seq dataset from young and old SVZ neurogenic niches^11^, we investigated the heterogeneity in cell cycle and cell adhesion pathways (Extended Data Fig. 6a). Using Seurat package (v3.1.5)^30^, single cells were assigned to different cell cycle stages (G0/G1, G2/M, and S phase) and assigned a continuous “G2/M score” and “S phase score” based on the expression levels of G2/M phase marker genes and S phase marker genes, respectively, by the Seurat CellCycleScoring() function. Low “S phase score” indicate that cells express low levels of S phase marker genes. The proportion of single cells from young and old SVZ neurogenic niches that belong to these three different cell cycle stages was then plotted as barplots (Extended Data Fig. 6b).

We plotted the relationship between Seurat’s “S phase score” and the cumulative expression of genes from the “Cell Adhesion” GO gene list (GO:0007155) in young and old qNSCs/astrocytes, aNSCs/NPCs, and neuroblasts (Extended Data Fig. 6c). For this analysis, each young cell population was downsampled to the number of old cells (480 qNSCs/astrocytes, 82 aNSCs/NPCs, and 146 neuroblasts). We found that some activated NSCs became more “quiescent-like” with age (Extended Data Fig. 6b,c). However, adhesion and proliferation could also be uncoupled, as some old activated NSCs can exhibit adhesion changes without exhibiting proliferation changes (Extended Data Fig. 6c). Although a subpopulation of old aNSCs/NPCs are more quiescent and could contribute to the changes in cell adhesion and migration, this subpopulation is unlikely to explain all adhesion changes with age, and age-dependent adhesion changes also occur at the single cell level.

### Immunofluorescence staining of young and old primary NSC cultures

NSCs were cultured as described above (see “*Primary NSC cell culture*”) in complete activated media until Passage 2-4 and then passaged with Accutase (STEMCELL Technologies, 07920) and seeded at a density of 5,000 cells per well in Matrigel (Corning, 354230, Lot #0062012, diluted [1:100] in cold DMEM/F12 (Thermo Fisher Scientific, 11320033))-coated coverslips in 24-well plates with complete activated or quiescent media. After 48 hours, adherent aNSCs/NPCs were washed 1x with PBS, then fixed with 4% paraformaldehyde (Electron Microscopy Sciences, 15714) for 15 min at room temperature. Wells were washed 4x with PBS and then stored in PBS, wrapped in parafilm, at 4°C until immunofluorescence staining.

Quiescent media was replaced every other day for 7 days and then adherent qNSCs were fixed with 4% paraformaldehyde (Electron Microscopy Sciences, 15714) for 15 min at room temperature, washed 4x with PBS, and then stored in PBS, wrapped in parafilm, at 4°C until immunofluorescence staining.

For immunostaining, cells were permeabilized with 0.1% Triton X-100 (Fisher Scientific, BP151) for 10 min at room temperature, and then washed 2x with PBS. Blocking was performed for 30 min with 1% BSA (Sigma, A7979) in 1x PBS. Primary staining was conducted for 1-1.5 hour at room temperature with phalloidin (Invitrogen, A12379, 665217, [1:500]), ALCAM/CD166 (Bio-techne, AF1172-SP, [1:40]), paxillin (Abcam, ab32084, [1:200]), or cleaved-caspase3 (Cell Signaling Technology, 9664T, [1:1000]) resuspended in 1% BSA (Sigma, A7979). Wells were then washed 4x with 1x PBS and secondary staining was performed for 1 hour at room temperature with Donkey anti-Rabbit Alexa 568 (Invitrogen, A10042, [1:500]) or Donkey anti-Goat Alexa 647 (Invitrogen, A21447, [1:500]) resuspended in 1% BSA (Sigma, A7979). DAPI (ThermoFisher, 62248 [1:500]) was added during secondary antibody staining. Wells were washed 4x with 1x PBS+0.2% TWEEN 20 (Sigma-Aldrich, P1379-1L), then 4x with 1x PBS. Coverslips were mounted onto glass slides with ProLong Gold Antifade Mountant with DAPI (ThermoFisher, P36931) and visualized with a Nikon Eclipse Ti confocal microscope equipped with a Zyla sCMOS camera (Andor) and NIS- Elements software (AR 4.30.02, 64-bit). Quantification of immunofluorescence staining of ALCAM was done using a custom pipeline in Fiji (v2)^35^. A mask was created using the phalloidin channel to create a cell mask and overlaid onto ALCAM channel. The sum of pixel intensity within the cell mask (RawIntDen) was then used to determine fluorescence intensity for each cell, normalized by cell size. For each experiment, values were normalized by dividing each cell’s fluorescence intensity by the mean fluorescence intensity of all cells in the young condition for aNSCs/NPCs and the old condition for qNSCs and the same threshold settings were used to create the phalloidin cell mask for all images within one experiment. For focal adhesion quantification, paxillin channel was used. Images were converted to 8-bit and background subtracted (rolling=5 sliding), threshold was set (kept the same for all images taken within one experiment, see Source Data for thresholding information), a region of interest (ROI) was drawn around cell periphery, and number of focal adhesions counted (“Analyze Particles”, size=0.5-infinity). For cleaved- caspase3 quantification, CellProfiler (v4.2.1) was used to identify and count nuclei (DAPI) using the IdentifyPrimaryObject function and cleaved-caspase3-positive cells using the IdentifySecondaryObjects function. Staining, imaging, and analysis was performed in a blinded manner. *P*-values were calculated with a two-tailed Mann-Whitney test comparing sample means. For immunofluorescence images (Fig. 2h,k, Extended Data Fig. 7e), brightness and contrast were adjusted in Fiji (v2) to enhance visualization. These adjustments were performed after all data quantification was complete. The same settings were applied to all images shown for each experiment.

We have also tested antibodies for a number of other cell adhesion proteins (ITGB1 (abcam, ab24693), ITGA6 (abcam, ab181551), EFNB2 (Invitrogen, PA5-106933), PLPP3 (Sigma, HPA072751), N-Cadherin (Cell Signaling Technology, 13116T), VE-Cadherin (Santa Cruz, sc- 9989), E-Cadherin (Cell Signaling Technology, 3195T), NRXN1 (abcam, ab222806), ATDC (Santa Cruz, sc-166707), and talin (Sigma-Aldrich, T3287), but the antibodies for did not work well for immunostaining or FACS on cultured NSCs due to non-specific staining or lack of signal. PTPRD (Bioworld, BS71853) showed a trending, but not significant increase with age in qNSCs and aNSCs/NPCs.

We used immunostaining instead of FACS for quantification, because of the potential concern that the proteolytic enzymes used to dissociate cells into single cell suspension for FACS could affect the localization, endocytosis, or cleavage of membrane proteins involved in cell adhesion.

### Detachment assay of cultured qNSCs and aNSCs/NPCs

For the cell detachment assay, aNSCs/NPCs were cultured in complete activated media until Passage 2-6 and then seeded at a density of 30,000 cells per well in PDL-coated 96-well plates (Falcon, 353072) with complete activated media. After 16-24 hours, media on adherent aNSCs/NPCs was replaced with complete activated media containing 10 nM Syto64 (Invitrogen, S11346, Lot #8344573) to visualize the cells and incubated for 1-2 hours. Media was replaced with complete activated media to remove excess Syto64. The plates were immediately brought to the Incucyte S3 system and imaged with the 4x objective for whole-well imaging with the red image channel. Each well was visually inspected for appropriate cell density, no cell clumping, and even distribution of cells. After imaging, media was removed and replaced with 100 μL of Accutase (STEMCELL Technologies, 07920) and incubated at room temperature for 5 min to lift the cells. Accutase (STEMCELL Technologies, 07920) was then removed and wells were washed once with 1x PBS and 100 μL of complete activated media was added to the plates. The plates were immediately imaged with Incucyte S3 4x objective for whole-well imaging with the red image channel. For analysis, well quantification area was restricted by 250 pixels using Incucyte’s built-in analysis software to decrease noise from cell debris accumulating at edges of well. Cell count was calculated using Incucyte’s built-in analysis software and percent cells remaining was calculated as 100*(amount of cells after Accutase/amount of cells before Accutase).

For cell detachment assay on quiescent NSCs, aNSCs/NPCs were first cultured in complete activated media until Passage 2-6 and then seeded at a density of 10,000 cells per well in PDL- coated 96-well plates with complete quiescent media. Complete quiescent media was replaced every other day for 7 days to induce quiescence. Next, qNSCs were incubated with Syto64, imaged, and analyzed as described above for aNSCs/NPCs with the following change: as qNSCs are more adhesive than aNSCs/NPCs, they were treated with Trypsin-EDTA (0.25%) phenol red (Thermo Fisher Scientific 25200114) (instead of Accutase) for 15 min at room temperature to lift the cells. *P-*values were calculated using a two-tailed Mann-Whitney test on the average of 2-4 technical replicates for each biological replicate. Experiments were not performed in a blinded manner; however, quantification of percent cells remaining was done in an automated manner using Incucyte’s analysis software for cell counting.

### NFIC CRISPR-Cas9 gene knockout in young and old cultured aNSCs/NPCs

#### Selection of gene to knockout

To knockout a transcription factor family involved in the program of genes identified to change with age in aNSCs, we used the following criteria: 1) Transcription factor identified in chromatin accessibility to change with age in aNSCs, 2) Transcription factor expressed in aNSC (using single cell RNA-seq), 3) Transcription factor known to be implicated in cell adhesion and migration. The NFI family of transcription factors (NFIA, NFIB, NFIC and NFIX) met these criteria. Among these 4 family members, we focused on NFIC because our pilot studies targeting this isoform were the most promising.

#### Cloning of sgRNAs

To knockout NFIC, we used a CRISPR/Cas9 approach, using aNSCs/NPCs cells microdissected from SVZs of young and old Cas9-expressing mice (Rosa26-Cas9^2^) and delivered guide RNAs to these cells using a lentivirus system. To deliver NFIC or Safe targeting control guide RNAs that target a safe harbor locus, we used the sgRNA-expressing plasmid MCB320 (gift from Mike Bassik Lab) (https://www.addgene.org/89359/) and subcloned sgRNAs of interest. This plasmid

contains a puromycin resistance gene and mCherry reporter for selection. See Supplementary Table 7 for all primers used for cloning and sequencing. The MCB320 plasmid was digested with the Blp1 and BstXI restriction enzymes and the gel-extracted, purified band was used for ligation reaction with a double-stranded oligonucleotide containing the sgRNA sequence. For the forward oligonucleotide of each sgRNA sequence, we added the following cloning adapter sequences: 5’-ttgg and 3’- gtttaagagc. For the reverse complement oligonucleotide of each sgRNA sequence, we used the reverse complement of the sgRNA sequence and added 5’- ttagctcttaaac and 3’ -ccaacaag. Each pair of oligonucleotides (IDT, standard desalting) (1 μM each of forward and reverse) was annealed in 98 μL of nuclease free duplex buffer (IDT, 11-05- 03-01) for 5 min at 95°C and then gradually cooled to room temperature. Then, 1 μL of a 1:20 dilution (in nuclease free duplex buffer (IDT, 11-05-03-01)) and was ligated with 500 ng of the purified digested plasmid band and then transformed in NEB stable competent cells (New England BioLabs, C3050H). Successful ligation of the guide was confirmed by Sanger sequencing using a mU6 sequencing primer (Supplementary Table 7).

#### Lentivirus production

For lentivirus production, human embryonic kidney 293T cells (HEK293T) were cultured in DMEM (Gibco, 11965092) + 10% FBS (Gibco, 10099-141) + 1% penicillin–streptomycin– glutamine (Gibco, 10378-016) (complete DMEM). HEK293T cells were plated at a density of 5 million cells per 10 cm plate (Greiner Bio-One, 664-160) in 10 mL of complete DMEM media.16 hours after plating, media was replaced on HEK293T cells with fresh complete DMEM. 3-4 hours later, HEK293T cells were transfected using polyethylenimine (PEI) (Polysciences, 23966-2, 1 mg/mL) and 3^rd^ generation lentivirus packaging vectors (pMDLg, pRSV, and pVSVG; gift from Mike Bassik lab). Per transfection, 50 μL of PEI was added to 900 μL of antibiotic and FBS free DMEM (Gibco, 11965092) and incubated at room temperature for 10 min. 0.5 μg of each packaging vector (pMDLg, pRSV, and pVSVG) was combined with 10 μg of MCB320 lentiviral plasmid containing guide of interest into a final volume of 50 μL. This DNA mix was then added to the DMEM-PEI mixture and incubated at room temperature for 20 min. PEI-DNA mixture was then added dropwise to HEK293T cells. 24 hours later, media was replaced with plain Neurobasal-A (Gibco, 10888-022) (no growth factors or B-27). Supernatant (10 mL) was collected 24 hours later and the media was replaced for another collection 48 hours after transfection. Supernatants were combined, centrifuged at 3000 rcf for 5 min, and then filtered through a 0.45 μm filter (EMD Millipore, SE1M003M00). Filtered supernatant was then flash frozen into 1.5 mL aliquots and stored at -80°C until ready for use. Multiple guide sequences were evaluated for knockout efficiency. Guides with high knockout efficiency was subsequently used for all experiments.

#### Infection of aNSCs/NPCs

Activated NSCs/NPCs were cultured from young and old mixed-sex Rosa26-Cas9^2^ knock-in mice as described in “*Primary NSC cell Culture*”. For infection, 1 million aNSCs/NPCs per well were plated onto PDL-coated 6-well plates in 1 mL of complete activated media. 24 hours later, media was removed from adherent aNSCs/NPCs. 0.75 mL of filtered viral supernatant was combined with 0.25 mL of complete activated media, supplemented with 4x growth factors and B-27 (final volume of growth factors and B-27 is 1x) and added to adherent aNSCs/NPCs. After 24 hours, media was replaced with 0.75 mL of filtered viral supernatant and 0.25 mL of completed activated media, supplemented with 4x growth factors and B-27 for a second round of infection. After 24 hours, media was changed to complete activated media. Two days later, successful infection was verified by microscopy using the mCherry reporter. Once mCherry

expression was confirmed, 0.5 μg/mL of puromycin (Sigma-Aldrich, P8833) was added to the culture media to select for cells that had been successfully infected. When all uninfected control aNSCs/NPCs died from puromycin selection (approximately 2-3 days later), adherent aNSCs/NPCs were washed with complete activated media to remove puromycin and allowed to expand for 24-48 hours as adherent cultures. aNSCs/NPCs were then passaged and plated onto PDL-coated 96-well plates for detachment assay. All remaining cells (usually 250,000 – 400,000) were plated onto PDL-coated 24-well plates for genomic DNA collection for verification of the knockout (see below). If there were not enough cells to perform the detachment assay and genomic DNA collection, cells were expanded as neurospheres for an additional 2-3 days. If cultures failed to recover from lentiviral transduction (continued slow growth), they were not used for subsequent experiments. 16-24 hours after plating cells, we performed the detachment assay as described above (see “*Detachment assay of cultured qNSCs and aNSCs/NPCs*”) except cells were incubated with Accutase for 3 min instead of 5 min. All experiments were performed in a paired-manner (paired young and old). *P-*values were calculated using a Wilcoxon Matched-Pairs Signed Ranks Tests comparing sample means. To take into account the paired-manner of the experiment, the difference in percent cells remaining (Old aNSC/NPC – Young aNSC/NPC) was also calculated. Difference in percent cells remaining greater than 0 indicates old cells are more adhesive than young cells. *P*-values comparing difference in percent cells remaining were calculated using a two-tailed Mann-Whitney test.

#### Verification of knockout efficacy

For genomic DNA isolation, aNSCs/NPCs were plated at a density of 250,000 – 400,000 cells per well of a PDL-coated 24-well plate at the same time as cells were plated for the detachment assay. After 16-24 hours, genomic DNA was harvested (same day as detachment assay). Cells were washed with 500 μL cold PBS and then lysed with 100 μL DirectPCR Lysis Reagent (Viagen Biotech, 102-T) with 1% Proteinase K (Fisher Scientific, 25-530-049) for 10 min at room temperature. Supernatant was pipetted repeatedly to lift cells and then transferred to PCR tubes and incubated at 65°C for 25 min and then 95°C for 15 min in a thermocycler. PCR was performed to amplify the region targeted by sgRNA and Sanger sequenced (see primers in Supplementary Table 7). Knockout efficiency was evaluated using Synthego ICE analysis tool (https://ice.synthego.com/#/^36^. With this tool, a score of 50 indicates 50% of the population sequenced was unedited (contains wild-type sequence). With this protocol, we achieved editing efficiencies of 58-84 (Safe targeting control guide, one guide sequence that edits a region of the genome that is considered “neutral”^37^) and 56-79 (*NFIC* targeting guide, one guide sequence that edits *NFIC*).

### Migration assay of cultured qNSCs and aNSCs/NPCs using live cell imaging

aNSCs/NPCs were cultured in complete activated media until Passage 2-4 and then seeded at a density of 200,000 cells per well in a PDL-coated 12-well plate with complete activated media treated with vehicle (H_2_O) or 10 μM Y-27632 (dissolved in H_2_O) (Tocris, 1254). After 48 hours, adherent aNSCs/NPCs were passaged with 500 μL of Accutase (STEMCELL Technologies, 07920), and resuspended in in Neurobasal-A (Gibco, 10888-022) supplemented with 2% B27 minus vitamin A (Gibco, 12587-010) and 1% penicillin–streptomycin–glutamine (Gibco, 10378- 016) with propidium iodide (BioLegend, 421301, 1:5000) for live/dead staining. 1000 live (PI-) cells per well were sorted with flow cytometry using the BD PICI onto PDL-coated Incucyte ImageLock 96-well plates (Essen BioScience, 4379) containing 100 μL of complete activated media with 10 nM Syto64 (Invitrogen, S11346, Lot #8344573) and either vehicle (H_2_O) or 10 μM Y-27632 (dissolved in H_2_O) (Tocris, 1254). The ImageLock plate was immediately brought to the Incucyte Zoom or Incucyte S3 live imaging system to image (with Phase and red image channels) at 37°C every 30 min for 20 hours with the 20X objective. After 20 hours, the cell media was washed 2x with PBS and then replaced with 100 μL of complete quiescent media.

Media was changed every other day for 7 days to induce quiescence. The media was then replaced with complete quiescent media supplemented with 10 nM Syto64 (Invitrogen, S11346, Lot #8344573) prior to imaging. Wells were imaged (with Phase and red image channels) using either the Incucyte S3 or Incucyte Zoom system with the 20X objective every hour for 20 hours at 37°C. Three images (from different parts of the well) at each timepoint were taken for each well and tracked cell metrics were summed across these three images. This 20 hour window includes time needed for cells to adhere to the plate (early time points) in addition to migration (later time points). Cell migration tracking was performed in an automated manner using Imaris (v9.3.0). Every image stack was manually inspected to ensure that cell tracking was correctly performed (e.g. that automated cell tracking was not incorrectly labelling debris and that cell tracks accurately follow individual cells). “Track Velocity” was output for visualization with Prism (v8) and *P*-values were calculated with a two-tailed Mann-Whitney test comparing both single cell values as well as sample mean values (where single cell velocities were averaged by animal). Experiments were not performed in a blinded manner; however, quantification of cell migration velocity was performed in an automated manner using Imaris (v9.3.0).

### Matrigel dispersion assay

For Matrigel dispersion assays, aNSCs/NPCs were cultured in complete activated media until Passage 4-6, passaged with Accutase (STEMCELL Technologies, 07920), and resuspended in complete activated media at a concentration of 100,000 cells/μL. Note that this assay integrates cell migration, as well as other factors (e.g. cell proliferation, survival, etc.). Matrigel (Corning, 354230, Lot #8344573 for experiments with young and old aNSCs/NPCs in Fig. 2 and Lot #9028255 for experiments with ROCKi in Fig. 4) was diluted 1:2 in cold F12:DMEM (Gibco, 11-330-057) while on ice to prevent solidification, and 50 μL was immediately added per well to a 96-well plate. We note that there are lot to lot variations in quantify of extracellular matrix proteins present in Matrigel which likely affect activated NSC’s ability to disperse through the material. The 96-well plate was moved to the 37°C incubator for 3 hours prior to spotting 50,000 cells (in a volume of 0.5 μL) directly to the center of the Matrigel-coated well. The plate was returned to the 37°C incubator for 15-30 min to allow for cells to adhere. Next, 150 μL of complete activated media (treated with vehicle (H_2_O) or 10 μM Y-27632 (dissolved in H_2_O) (Tocris, 1254)) was very carefully added to each well (for the ROCKi experiments), and the 96- well plate was placed in either the Incucyte Zoom or Incucyte S3 live imaging system to image every 6 hours for 48 hours using the 4x objective to image the entire well. For each biological replicate, 1-4 technical replicates were quantified to account for difficulties in spotting 0.5 μL volumes, and the distances migrated at each timepoint were averaged among technical replicates. We manually quantified dispersion distance, defined as the maximum distance from the initial spotting perimeter to the outermost cell body, using Fiji (v2)^35^. Neither imaging nor analysis was performed in a blinded manner. *P*-values were calculated with a two-tailed Mann-Whitney test at each 12-hour timepoint. At early time points, cell proliferation/survival are less likely to be strong contributing factors in the Matrigel assay than at later time points.

### *In vitro* EdU pulse assay of cultured aNSCs/NPCs

Cultured aNSCs/NPCs from young and old male C57BL/6 mice were plated onto PDL-coated coverslips in 24-well plates with complete activated media with or without 10 μM ROCK inhibitor (ROCKi) compound Y-27632 (dissolved in H_2_O) (Tocris, 1254). Forty-eight hours after plating, 10 μM of EdU (Thermo Fisher Scientific, C10086) was added to adherent aNSCs/NPCs culture. Two hours later, aNSCs/NPCs were washed once with 1x PBS and then fixed with 4% paraformaldehyde and processed for immunostaining as previously described (see “*Immunofluorescence staining of young and old cultured NSCs*”), except in this case CliCK-IT EdU Alexa Fluor 488 imaging protocol (Thermo Fisher Scientific, C10337) was performed, according to the manufacturer’s instruction, for 30 min at room temperature prior to blocking.

Samples were imaged at 20x with a Nikon Eclipse Ti confocal microscope equipped with a Zyla sCMOS camera (Andor) and NIS- Elements software (AR 4.30.02, 64-bit). CellProfiler (v4.2.1) was used to identify and count nuclei (DAPI) using the IdentifyPrimaryObjects function and EdU-positive nuclei using the IdentifySecondaryObjects function. Staining, imaging, and analysis was performed in a blinded manner. *P*-values were calculated with a two-tailed Mann-Whitney test comparing sample means.

### Immunofluorescence staining of SVZ coronal sections to measure distance of quiescent and activated NSCs relative to the ventricle

For immunofluorescence staining of SVZ coronal sections to measure distance of quiescent and activated NSCs relative to the ventricle, four young (4 months old) and four old (18-26 months old) GFAP-GFP animals of both sexes (2 male and 2 female animals per age condition), combined over two experiments, were subjected to intracardiac perfusion with 4% paraformaldehyde (Electron Microscopy Sciences, 15714) in 1x PBS. We used male and female GFAP-GFP mice for these experiments, similar to what was used for generating ATAC-seq libraries from freshly isolated cells from neurogenic niches (Fig. 1), see “*ATAC-seq library generation from freshly isolated cells*”. Brains were post-fixed overnight in 4% paraformaldehyde (Electron Microscopy Sciences, 15714) and then dehydrated in 30% sucrose (Sigma-Aldrich, S3929) for 72 hours. Brains were embedded in Tissue-Tek optimal cutting temperature (O.C.T.) compound (Electron Microscopy Sciences, 62550), sectioned in 16 μm coronal sections using a cryostat (Leica CM3050S), and then mounted on electrostatic glass slides (Fisher Scientific, 12-550-15). To ensure a similar anatomic region of the lateral ventricle was being immunostained and analyzed in each animal, the corpus callosum was first identified in the coronal sections as an anatomical landmark. For these experiments, 6 serial coronal sections spaced approximately 100 μm apart were selected for immunostaining. These serial coronal sections correspond to images 42 – 47 from the Allen Institute Coronal Brain Atlas (https://mouse.brain-map.org/static/atlas). The 3 more anterior serial coronal sections were stained with antibodies to GFAP, Ki67, Vinculin, and with DAPI and the 3 more posterior serial coronal sections were stained with antibodies to S100a6, Ki67, ARL13b, and with DAPI (see below).

To perform immunofluorescence staining, coronal sections were washed with 1x PBS and then permeabilized with ice-cold methanol + 0.1% Triton X-100 (Fisher Scientific, BP151) for 15 min at room temperature. Slides were washed 3x with 1x PBS for 5 min. Antigen retrieval was performed in 10 mM sodium citrate buffer (pH 6.0) (2.94 g Tri-sodium citrate dihydrate (Sigma- Aldrich, S1804) in 1000 mL milliQ H_2_O adjusted to pH 6.0 with 1N HCl) + 0.05% TWEEN 20 (Sigma-Aldrich, P1379-1L) in a 85°C water bath for 2 hours. Slides were then cooled to room temperature for 20 min (in antigen retrieval buffer), and coronal sections were blocked with 5% Normal Donkey Serum (NDS) (ImmunoReagents Inc., SP-072-VX10) and 1% BSA (Sigma, A7979) in 1x PBS for 30 min at room temperature. Primary antibody staining was performed overnight at 4°C in 5% Normal Donkey Serum and 1% BSA in 1x PBS. Primary antibodies used were the following: Ki67 (Invitrogen, Clone SolA15, 14-5698-82, [1:200]), GFAP (Abcam, ab53554, [1:500]), Vinculin (Abcam, ab129002, [1:200]), S100a6 (Abcam, ab181975, [1:500]),

and ARL13b (Abcam, ab136648, [1:500]). Vinculin (cell adhesion protein) and ARL13b (ciliary marker) were used to demarcate ventricle border, for GFAP staining and S100a6 staining, respectively. Different antibodies were used to mark the ventricular wall because of species compatibility for the staining panel. Coronal sections were then washed 3x with 1x PBS and 0.2% TWEEN 20 for 10 min at room temperature followed by two 1x PBS washes for 15 min.

Secondary antibody staining was performed at room temperature for 2 hours in 5% NDS and 1% BSA in 1x PBS. Secondary antibodies used were the following: Donkey anti-Goat 488 (Sigma Aldrich, SAB460032-250UL [1:1000]), Donkey anti-Rat 594 (Life Technologies, A21209 [1:1000]), Donkey anti-Mouse 488 (Life Technologies, A21202, [1:1000]), Donkey anti-Goat 488 (Invitrogen, A11055, [1:1000]), Donkey anti-Rabbit 568 (Invitrogen, A10042, [1:1000]), and/or Donkey anti-Rat 647 (Invitrogen, A48272, [1:1000]). DAPI (ThermoFisher, 62248 [1:500]) was added during secondary antibody staining. Sections were washed 3x for with 1x PBS and 0.2% TWEEN 20 for 10 min at room temperature followed by 3 1x PBS washes for 5 min. Coronal sections were mounted with ProLong Gold Antifade Mountant with DAPI (ThermoFisher, P36931). Multiple Z-stacks using the 60x objective of a Nikon Eclipse Ti confocal microscope equipped with a Zyla sCMOS camera (Andor) and NIS- Elements software (AR 4.30.02, 64-bit) were captured to image the entire length of the SVZ from one hemisphere per section (3 sections per animal for GFAP staining and 3 sections per animal for S100a6 staining). For immunofluorescence images (Fig. 3c, Extended Data Fig. 9a,b) brightness and contrast were adjusted in Fiji (v2) to enhance visualization. These adjustments were performed after all data quantification was complete and the same settings were applied to all images shown for each experiment.

### Quantification of the location of quiescent and activated NSCs with respect to the ventricle in coronal sections

For quantification of the location of quiescent and activated NSCs with respect to the ventricle in coronal sections, captured Z-stacks (30 slices per stack for GFAP staining and 15 slices per stack for S100a6 staining) were transformed into a Z-projection (sum slices) and quantified using a custom pipeline. Analysis was restricted to the region that is 200 μm from ventricle wall to not include striatal astrocytes (though the vast majority of cells were less than 50 μm away from the ventricle wall) (Fig. 3c, Extended Data Fig. 9a,b).

GFAP and S100a6 were used as NSC markers. For staining with GFAP, GFAP+/Ki67- cells were identified as qNSCs/astrocytes and GFAP+/Ki67+ cells were identified as aNSCs. For staining with S100a6, S100a6+/Ki67- cells were identified as qNSCs and S100a6+/Ki67+ cells were identified as aNSCs. See also below, *“Single cell RNA-seq to detect potential ependymal- repairing SVZ astrocytes and reactive astrocytes”.* NSCs directly lining blood vessels were censored in the analysis because they were considered to be localized to a blood vessel rather than the niche lining the ventricle wall. The center of each NSC nuclei was chosen as a consistent fixed point to assess cell location within the niche. Distances from the ventricle were calculated by measuring Euclidean distance from the center of the nucleus of each cell of interest to the ventricle border in Fiji (v2). Neither imaging nor cell type annotation was performed in a blinded manner; however, distance quantification was performed in an automated manner. *P*- values were calculated with a two-tailed Mann-Whitney test comparing sample means.

### Immunofluorescence staining of SVZ sagittal sections for distance of quiescent and activated NSCs and neuroblasts relative to the ventricle

For immunofluorescence staining of SVZ sagittal sections for distance of quiescent and activated NSCs and neuroblasts relative to the ventricle, 5 young male (2-3 months) and 5 old male (21 months) C57BL/6 mice were used (combined over 2 independent experiments) (note that these animals were the same as those used in the 4 hour time point in “*In vivo EdU-labelling to follow location of cells in subventricular zone, rostral migratory stream, and olfactory bulb*”, see below). Mice were intraperitoneally injected with 5-ethynyl-2’-deoxyuridine (EdU) (Fisher Scientific, A10044) (resuspended in PBS at 5 mg/mL) at a dose of 50 mg/kg 4 hours prior to intracardiac perfusion with 4% paraformaldehyde (PFA) (Electron Microscopy Sciences, 15714) in PBS.

Brains were processed and immunostained in a similar manner as coronal sections (see above “*Immunofluorescence staining of SVZ coronal sections to measure distance of quiescent and activated NSCs relative to the ventricle*”), except they were sectioned in 16 μm sagittal sections rather than coronal. Sagittal sections corresponding approximately to image 14 from the Allen Institute Sagittal Brain Atlas (https://mouse.brain-map.org/static/atlas) were stained and imaged as described in “*Immunofluorescence staining of SVZ coronal sections for distance of NSCs relative to the ventricle*” with the antibodies described below.

For immunostaining for quiescent NSCs/astrocytes and activated NSCs in sagittal sections, primary antibodies used were the following: Ki67 (Invitrogen, Clone SolA15, 14-5698-82, [1:200]), GFAP (Abcam, ab53554, [1:500]), and Vinculin (Abcam, ab129002, [1:200]).

Secondary antibodies used were the following: Donkey anti-Goat 488 (Sigma Aldrich, SAB460032-250UL [1:1000]), Donkey anti-Rabbit 568 (Invitrogen, A10042, [1:1000]), and Donkey anti-Rat 647 (Invitrogen, A48272, [1:1000]). DAPI (ThermoFisher, 62248 [1:500]) was added during secondary antibody staining.

For immunostaining for EdU+ activated NSCs/NPCs and neuroblasts, the CliCK-IT EdU Alexa Fluor 488 imaging protocol (Thermo Fisher Scientific, C10337) was performed for 30 min at room temperature prior to blocking, according to the manufacturer’s instructions. The following primary antibodies were used: Ki67 (Invitrogen, Clone SolA15, 14-5698-82, [1:200]) and DCX (Cell Signaling Technologies, 4604 [1:500]). The following secondary antibodies were used: Donkey anti-Rabbit Alexa 568 (Invitrogen, A10042 [1:500]) and Goat anti-Rat Alexa 647 (Invitrogen, A21247 [1:500]). DAPI (1 mg/mL) (Thermo Fisher, 62248) was included at a concentration of 1:500 with the secondary antibody mix.

Multiple images tiling the entire length of the ventricle from one sagittal section per animal were captured with a 20x objective using a Zeiss LSM 900 confocal microscope equipped with Zeiss ZEN Blue 3.0 software. Images were automatically stitched together using the Zeiss ZEN Blue 3.0 software to create a single image containing the entire SVZ. For immunofluorescence images (Fig. 3f), brightness and contrast were adjusted in Fiji (v2) to enhance visualization. These adjustments were performed after all data quantification was complete and the same settings were applied to all images shown for each experiment.

### Quantification of the location of quiescent and activated NSCs and neuroblasts with respect to the ventricle in sagittal sections

For quantification of the location of quiescent and activated NSCs and EdU+ aNSCs/NPCs and EdU+ neuroblasts with respect to the ventricle in sagittal sections, one section containing the entire rostral migratory stream (RMS) was used for quantification to ensure consistency in the region being analyzed from animal to animal. Analysis of the SVZ was restricted to the region that is 200 μm from ventricle wall to not include striatal astrocytes (though the vast majority of cells were less than 50 μm away from the ventricle wall) (Fig. 3f).

In the staining panel with DAPI, GFAP, Ki67, and Vinculin, GFAP+/Ki67- cells were considered qNSCs/astrocytes and GFAP+/Ki67+ were considered aNSCs. In staining panel with DAPI, EdU, Ki67, and DCX, EdU+ cells were classified as aNSCs/NPCs if they were Ki67+/DCX- and as neuroblasts if they were DCX+ (and either Ki67+ or Ki67-). Quantification and analysis of distance of qNSCs/astrocytes, aNSCs, EdU+ aNSCs/NPCs, and EdU+ neuroblasts to the ventricle was performed as described above (“*Quantification of the location of quiescent and activated NSCs with respect to the ventricle in coronal sections*”).

### Single cell RNA-seq to detect potential ependymal-repairing SVZ astrocytes and reactive astrocytes

#### Ependymal-repairing SVZ astrocytes

SVZ astrocytes that intercalate into the ependymal lining are GFAP+ and proliferative and can take on ependymal markers^38, 39^. To test if old aNSC populations could contain ependymal- repairing astrocytes, we performed UMAP analysis on cells from the SVZ neurogenic niche as previously described^33^ on single cell gene expression data of 21,458 single cell transcriptomes of cells^33^ from the SVZ neurogenic niches of 28 mice, tiling ages from young to old using the Seurat package (v4.0.5)^30^. We examined this dataset first because it contains many ependymal cells. We filtered for cells expressing markers of ependymal-repairing SVZ astrocytes (*Gfap, S100b, Cd24a,* and *Ctnnb1*)^38, 39^ using FeaturePlot(min.cutoff = “q10”, max.cutoff = “q90”) (Seurat v4.0.5). This identified 4 cells that expressed these 4 markers. All 4 of these cells clustered with ependymal cells. The presence of these cells did not appear to be dependent on old age, as these 4 cells were from animals that were, respectively, 4.7, 5.4, 16.83, and 18.87 months old. We also examined the presence of ependymal-repairing astrocytes in our other single cell RNA-seq dataset^11^ (which has very few ependymal cells), and we also only observed 2 cells exhibiting markers of ependymal-repairing SVZ astrocytes (out of 14,685 cells), 1 from young and 1 from old animals. Together, these data suggest that ependymal-repairing SVZ astrocytes are present but not numerous in the SVZ neurogenic niche of old animals.

#### Reactive astrocytes

To test if any cells in our single cell datasets from the SVZ neurogenic niches of 28 mice^33^ or from young and old SVZ neurogenic niches^11^ (described above) expressed markers of reactive astrocytes, we performed similar UMAP analysis. We used markers of A1 reactive astrocytes (produced in response to neuroinflammation)^40^, A2 reactive astrocytes (produced in response to ischemia)^40^, and pan-injury reactive astrocytes (identified from meta-analysis of 15 transcriptomic reactive astrocyte datasets)^41^(see Supplementary Table 8 for full list of gene markers). We did not detect any cell in either dataset (out of 21,458 cells in one dataset^33^ and 14,685 cells in another dataset^11^) that expressed these types of reactive astrocyte markers^40, 41^. In addition, the ependymal-repairing SVZ astrocytes we identified above did not express these markers.

### *In vivo* EdU-labelling to follow location of cells in subventricular zone, rostral migratory stream, and olfactory bulb

For these experiments, 12 young (2-3 months) and 12 old (21 months) male C57BL/6 animals (combined over 2 independent experiments) were intraperitoneally injected with 5-ethynyl-2’- deoxyuridine (EdU) (Fisher Scientific, A10044) (resuspended in PBS at 5 mg/mL) at a dose of 50 mg/kg either 4 hours, 2 days, or 7 days prior to intracardiac perfusion with 4% paraformaldehyde (PFA) (Electron Microscopy Sciences, 15714) in PBS. The 5 young (2-3 months) and 5 old (21 months) male C57BL/6 that were intraperitoneally injected with EdU 4 hours before intracardiac perfusion used here were the same as the animals used above in “*Immunofluorescence staining of SVZ sagittal sections for distance of quiescent and activated NSCs and neuroblasts relative to the ventricle*” and “*Quantification of distance of EdU-positive aNSCs/NPCs and neuroblasts from the ventricle in sagittal sections*”. Brains were processed and immunofluorescence staining of sagittal sections was performed as described above (see “*Immunofluorescence staining of SVZ sagittal sections for distance of quiescent and activated NSCs and neuroblasts relative to the ventricle*”).

Multiple sections per brain, corresponding approximately to image 14 from the Allen Institute Sagittal Brain Atlas (https://mouse.brain-map.org/static/atlas) were stained and imaged. Sagittal sections were selected for quantification based on presence of an intact RMS as visualized by a stream of DCX-positive cells connecting the SVZ to the OB. Very few sections contained a fully intact RMS and one young 7-day post-injection replicate from the second experiment was censored due to lack of high-quality sections at the appropriate depth. Images of sagittal sections were captured using a 5x objective, and 9x3 images (for a total of 27) were automatically stitched together using the Zeiss ZEN Blue 3.0 software. For quantification, EdU-labelled nuclei were counted using Fiji (v2)^35^. Every image was converted to 8-bit, the threshold was adjusted ((90,255) for replicate 1 and (70,255) for replicate 2), watershed was applied to the image and EdU-labelled nuclei were counted in an automated fashion with “Analyze Particles (size (micron^2): (20-infinity) for replicate 1 and 2, and circularity: (0.2-1.00) for replicate 1)”. EdU- labelled nuclei were quantified in three regions (along the entire length of the ventricle for SVZ, the entire length of the RMS, and the entire OB), which were manually defined using a hand-drawn region of interest (ROI) for each section. Neither imaging nor analysis was performed in a blinded manner; however, cell counting was performed in an automated manner with the same thresholding for all images for each replicate. *P*-values were calculated with a two-tailed Mann-Whitney test. For immunofluorescence images (Fig. 3i,j, Extended Data Fig. 9j), brightness and contrast were adjusted in Fiji (v2) to enhance visualization. These adjustments were performed after all data quantification was complete and the same settings were applied to all images shown for each experiment.

### Quantification of efficiency of EdU incorporation

We used the same sagittal section images that were quantified above for EdU localization to SVZ, RMS, and OB (see “*In vivo EdU-labelling to follow location of cells in subventricular zone, rostral migratory stream, and olfactory bulb*”). To calculate EdU labelling efficiency, we analyzed the number of EdU-labeled nuclei in the SVZ 4 hours after EdU injection. Number of EdU-labelled nuclei in the SVZ was quantified as described above using Fiji (v2). The number of Ki67+ cells was quantified in the same region using the same pipeline as counting EdU-labelled nuclei. See Source Data for thresholding information. The same threshold was applied to all images captured the same day. To calculate EdU labeling efficiency 4 hours after EdU injection, the number of EdU+ cells was divided by the total number of Ki67+ in the SVZ in each image.

Neither imaging nor analysis was performed in a blinded manner; however, cell counting was performed in an automated manner with the same thresholding for all images for each replicate. *P*-values were calculated with a two-tailed Mann-Whitney test.

### Ingenuity Pathway Analysis (IPA)

Genes associated with differentially accessible peaks (FDR<0.05) that open in old aNSCs freshly isolated from the SVZ were uploaded to Ingenuity Pathway Analysis (IPA) (v1.16)^42^ (QIAGEN Inc., https://www.qiagenbioinformatics.com/products/ingenuitypathway-analysis) to identify age-related regulatory changes (see Supplementary Table 6). For each peak-associated gene, we uploaded the log fold-change, p-value, and FDR (q-value) and based the IPA analysis on FDR. Statistical enrichment of pathways was reported with *P*-values calculated by IPA using Fisher’s Exact Test.

### RGD molecular tension sensor FRET measurements of cultured NSCs

Sensors containing a minimal RGD sequence derived from fibronectin (TVYAVTGRGDSPASSAA) were expressed, purified, and labeled with AlexaFluor-546 and - 647, and coverslips passivated with maleimide polyethylene glycol (PEG) succinimidyl carboxymethyl ester (JenKem Technology, A5003-1) were prepared as previously described^43, 44^. Flow chambers for imaging were prepared as described^44^ with slight modifications. 8-well flow chambers (Grace BioLabs, #622505, ∼55 µL channel volume) were attached to PEGylated coverslips as previously described^45^. Chambers were incubated with 300 nM double-labeled sensor at room temperature for 45 min. The chambers were then washed with 150 µL of 1x PBS for 1 min to prevent non-specific cell attachment. 60 µL of cell suspensions at a density of 300,000-400,000 cells/mL were added to the channels, and the chambers were incubated for 3 hours at 37°C and 5% CO2 in complete activation media to allow time for cells to spread.

Chambers were then washed with 150 µL of warm media, and brightfield and FRET measurements were made immediately after and acquired with an objective heater (Bioptechs) set to 37°C. For measurements on cells treated with the ROCK inhibitor Y-27632, complete activated media was supplemented with 10 μM Y-27632 (dissolved in H_2_O) (Tocris, 1254)) for cell suspensions and media washes.

We limited our measurements and analysis to individual cells that looked well-spread in brightfield and cell clusters of no more than 5 cells in which cell outlines could be clearly distinguished. Under conditions used, the vast majority of cells fulfilled the criteria of being “well-spread in brightfield with cell clusters of no more than 5 cells”. FRET fluorescence measurements were performed with objective-type total internal reflection fluorescence (TIRF) microscopy on an inverted microscope (Nikon TiE) with an Apo TIRF 100× oil objective lens, numerical aperture 1.49 (Nikon) as described previously^43^ and controlled using Micromanager^46^. Samples were excited with 532-nm (Crystalaser) or 635-nm (Blue Sky Research) lasers. Emitted light passed through a quad-edge laser-flat dichroic with center/bandwidths of 405/60 nm, 488/100 nm, 532/100 nm, and 635/100 nm from Semrock Inc. (Di01-R405/488/532/635-25×36) and corresponding quad-pass filter with center/bandwidths of 446/37 nm, 510/20 nm, 581/70 nm, 703/88 nm band-pass filter (FF01-446/510/581/703-25). Donor and acceptor images were taken through separate additional cubes stacked into the light path (donor: 550 nm long-pass; acceptor: 679/41 nm and 700/75 nm) and recorded on a Hamamatsu Orca Flash 4.0 camera.

Images were prepared in Fiji (v2)^35^ and analyzed using custom MATLAB scripts, in which FRET efficiencies were computed and thresholded to identify adhesions and quantify forces within adhesions. Adhesions were identified based on FRET values measured using the RGD tension sensor. FRET efficiencies were computed for each pixel in each image, and regions exceeding specific FRET efficiencies and area cutoffs were identified as adhesions using a Watershed algorithm. Forces at adhesions were substantially greater than any signal beneath the cell body, which is similar to background, consistent with previous force measurements at focal adhesions^43^ (Extended Data Fig. 10g,h).

### Immunofluorescence staining of cultured NSCs treated with ROCK inhibitor

aNSCs/NPCs were cultured as described above (see *Primary NSC cell culture*) until Passage 2-4 and then passaged with Accutase (STEMCELL Technologies, 07920) and seeded at a density of 5,000 cells per well on Matrigel (Corning, 354230, Lot #0062012, diluted 1:100 in DMEM/F12 (Thermo Fisher Scientific, 11320033))-coated coverslips in 24-well plates with complete activated or quiescent media with or without 10 μM Y-27632 (dissolved in H_2_O) (Tocris, 1254). After 48 hours, adherent aNSCs/NPCs (with or without 10 μM Y-27632 (dissolved in H_2_O) (Tocris, 1254)) were washed 1x with 1x PBS, then fixed with 4% paraformaldehyde (Electron Microscopy Sciences, 15714) for 15 min at room temperature. Wells were washed 4x with 1x PBS and then stored, wrapped in parafilm, at 4°C until immunofluorescence staining. For quiescent NSCs, cells were plated at the density of 5,000 cells per well in a Matrigel (Corning, 354230, Lot #0062012, diluted 1:100 in DMEM/F12 (Thermo Fisher Scientific, 11320033))- coated 24-well plate in quiescent media (with or without 10 μM Y-27632). The quiescent media (with or without 10 μM Y-27632) was replaced every other day for 7 days and qNSCs were then fixed with 4% paraformaldehyde as described above. Staining and quantification for ALCAM, focal adhesions (paxillin), and cleaved-caspase3 was performed in the same manner as described above (see “*Immunofluorescence staining of young and old cultured NSCs*”). Staining for DCX was performed in the same manner as for ALCAM, paxillin, and cleaved-caspase3.

Quantification for DCX+ cells was performed using CellProfiler (v4.2.1). The function IdentifyPrimaryObjects was used to identify and count nuclei (DAPI) which served as a seed for IdentifySecondaryObjects to identify and count cells that were DCX-positive. Antibodies used were phalloidin (Invitrogen, A12379, 665217, [1:500]), ALCAM/CD166 (Bio-techne, AF1172- SP, [1:40]), paxillin (Abcam, ab32084, [1:200]), and DCX (Cell Signaling Technologies, 4604 [1:500]) resuspended in 1% BSA (Sigma, A7979). Staining, imaging, and analysis was performed in a blinded manner. *P*-values were calculated with a two-tailed Mann-Whitney test comparing sample means. For immunofluorescence images (Fig. 4h), brightness and contrast were adjusted in Fiji (v2) to enhance visualization. These adjustments were performed after all data quantification was complete. The same settings were applied to all images shown for each experiment.

### Osmotic pump surgery for *in vivo* intracerebroventricular delivery of ROCK inhibitor in old mice

For delivery of ROCK inhibitor in the lateral ventricle of old mice, Y-27632 (ROCKi) (Tocris, 1254) was diluted in artificial CSF (Tocris, 3525) in a total volume of 200 μL to a final concentration of 100 μM. A concentration of 100 μM was chosen based on a previous study^47^. The diluted Y-27632 or vehicle control (artificial CSF only) were loaded into 200 μL osmotic pumps (ALZET/Durect, model 2002) with a 14-day infusion rate of 0.5 μL per hour and pumps were allowed to equilibrate in PBS in a 37°C water bath overnight. Before surgery, mice were anaesthetized with isofluorane and received post-surgical buprenorphine and saline. Osmotic pumps were connected to a cannula (ALZET, Brain infusion kit III) and inserted by stereotactic surgery at +1 mm lateral, −0.3 mm anterior–posterior, and −3mm deep relative to bregma in order to target the right lateral ventricle of old (20-22 months old) male C57BL/6 mice (8-10 mice per condition (vehicle control or ROCKi), combined over two independent experiments). The pump connected to the cannula was then placed subcutaneously along the back of the mouse. Mice were randomized to either the control or the Y-27632 group (ROCKi) prior to surgery. The day before surgery, mice were singly-housed and cage-mates were separated across experimental conditions. We alternated performing the surgery between control and Y-27632 mice to avoid batch effect.

One week after start of drug delivery, mice were intraperitoneally injected with 5-ethynyl-2’- deoxyuridine (EdU) (Fisher Scientific, A10044) (resuspended in 1x PBS at 5 mg/mL) at a dose of 50 mg/kg either 4 hours or 7 days prior to intracardiac perfusion with 4% paraformaldehyde (PFA) (Electron Microscopy Sciences, 15714) in PBS.

### Quantification of distance of NSCs relative to ventricle and of location of EdU+ cells in different locations in old mice treated with ROCK inhibitor

Quantification of sagittal distance of aNSCs/NPCs was performed as described above in “*Quantification of the location of quiescent and activated NSCs and neuroblasts with respect to the ventricle in sagittal sections*”, except that all Ki67+/DCX- cells were counted as aNSCs/NPCs (rather than filtering first by cells that are EdU-positive) and images were captured with the 60x objective of a Nikon Eclipse Ti confocal microscope equipped with a Zyla sCMOS camera (Andor) and NIS- Elements software (AR 4.30.02, 64-bit). *P*-values were calculated with a two-tailed Mann-Whitney test comparing sample means.

For quantification of number of EdU cells in the entire SVZ (along the entire ventricle), entire length of the RMS, and entire OB, sagittal sections were immunostained, imaged, and analyzed as described above in “*In vivo EdU-labelling to follow location of cells in SVZ, RMS, and OB*” except analysis of EdU-positive cell localization (in SVZ, RMS, or OB) was performed in a blinded manner.

Brain sections adjacent to the ones used for quantification of number of EdU cells in the entire SVZ, entire RMS, and entire OB were immunostained as described above in “*Immunofluorescence staining of SVZ sagittal sections for distance of quiescent and activated NSCs and neuroblasts relative to the ventricle*”. The following primary antibodies were used: DCX (Cell Signaling Technologies, 4604 [1:500]) and NeuN (Millipore, Clone A60, MAB377, 2919670 [1:500]). The following secondary antibodies were used: Donkey anti-Rabbit Alexa 647 (Invitrogen, A31573 [1:500]) and Donkey anti-Mouse Alexa 568 (Invitrogen, A10037 [1:500]). DAPI (1 mg/mL) (Thermo Fisher, 62248) was included at a concentration of 1:500 with the secondary antibody mix. Images of EdU-positive cells in the OB were captured with the 60x objective of a Nikon Eclipse Ti confocal microscope equipped with a Zyla sCMOS camera (Andor) and NIS- Elements software (AR 4.30.02, 64-bit). For immunofluorescence images (Fig. 5e,h,l), brightness and contrast were adjusted in Fiji (v2) to enhance visualization. These adjustments were performed after all data quantification was complete and the same settings were applied to all images shown for each experiment.

### Statistical analyses

For all experiments, young and old conditions were processed in an alternate manner rather than in two large groups, to minimize the group effect. We did not perform power analyses, though we did take into account previous experiments to determine the number of animals needed. To calculate statistical significance for experiments, all tests were two-sided Mann-Whitney tests (with the exception of Fisher’s Exact Tests and Wilcoxon Matched-Pairs Signed Ranks Tests which are noted in Figure Legends). Results from individual experiments and all statistical analyses are included in Source Data.

---

**Extended Data Figure 1:**
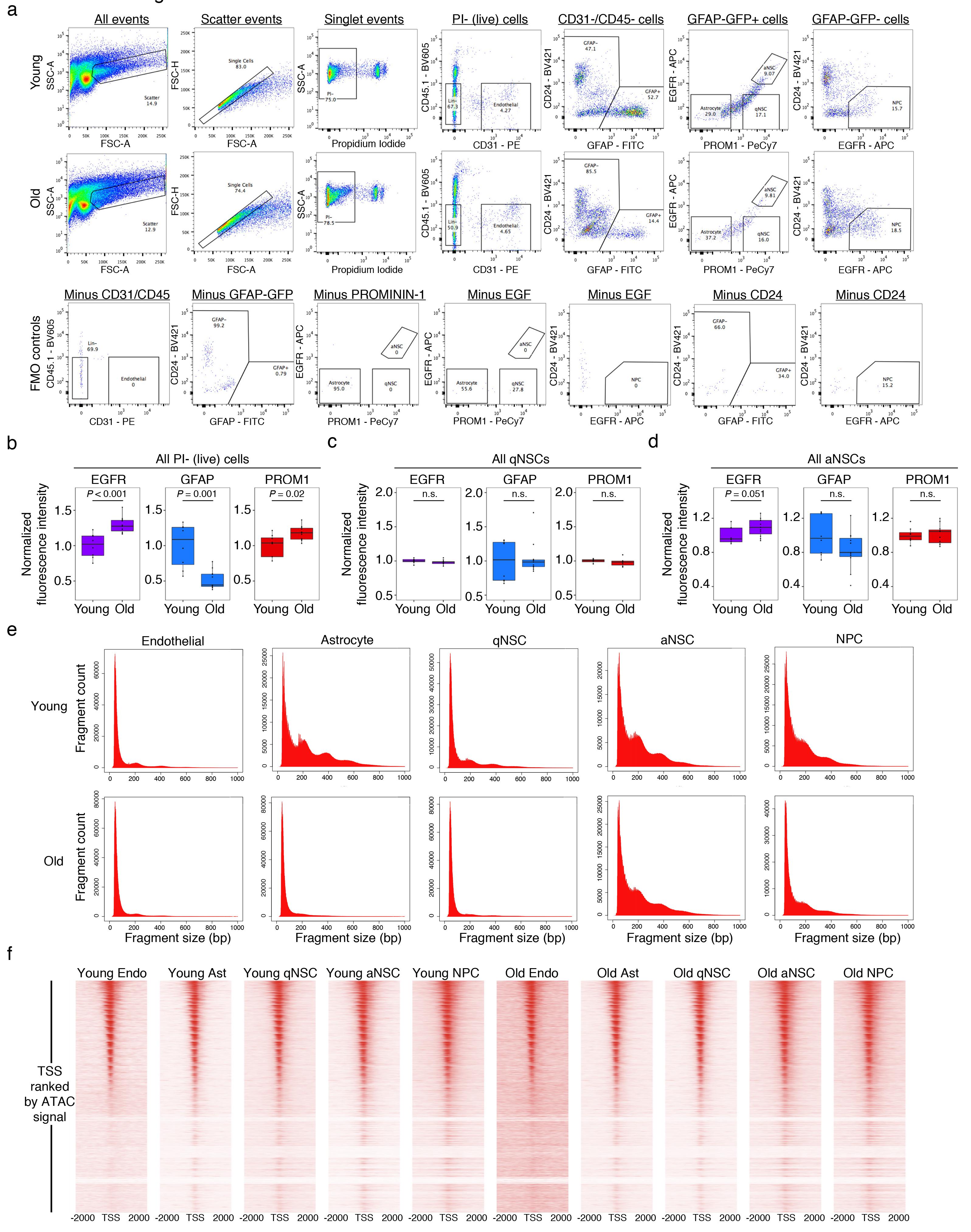
Quality control metrics for FACS and *in vivo* ATAC-seq libraries. a, FACS gating scheme used to isolate 5 cell populations from the SVZ of young and old GFAP-GFP mice. Negative controls (fluorescence minus one (FMO) controls) for each stain are indicated. Negative controls (FMO controls) were stained with all antibodies other than the one(s) for which it was the negative control. b, Changes in FACS markers with age for all live cells. Normalized scaled fluorescence intensity of indicated FACS markers (EGFR, GFAP, and PROM1) of young (4-5 months) and old (20-24 months) PI-negative sorted cells used for generating ATAC-seq libraries from freshly isolated cells from the SVZ neurogenic niche. *n* = 8 young male and female GFAP-GFP mice and *n* = 11 old male and female GFAP-GFP mice combined over 2 experiments. Data normalized to young mean of each individual experiment. Boxplots display median, and lower and upper quartiles. Dots represent mean fluorescence intensity for an individual animal. *P*-values were calculated using a two- tailed Mann-Whitney test. Data from independent experiments are in Source Data. The change in marker expression in all live cells likely reflect changes in cell type composition that occur with age with age. c, Changes in FACS markers with age in quiescent NSCs. Normalized scaled fluorescence intensity of indicated FACS markers (EGFR, GFAP, and PROM1) of young (4-5 months) and old (20- 24 months) FACS-purified quiescent NSCs used for generating ATAC-seq libraries from freshly isolated cells from the SVZ neurogenic niche. *n* = 8 young male and female GFAP-GFP mice and *n* = 11 old male and female GFAP-GFP mice combined over 2 experiments. Data normalized to young mean of each individual experiment. Note that EGFR raw values were very low, as expected, for quiescent NSCs (GFAP+PROM1+EGFR-). Boxplots display median, and lower and upper quartiles. Dots represent mean fluorescence intensity for an individual animal. *P*-values were calculated using a two-tailed Mann-Whitney test. Data from independent experiments are in Source Data. d, Changes in FACS markers with age in activated NSCs. Normalized scaled fluorescence intensity of indicated FACS markers (EGFR, GFAP, and PROM1) of young (4-5 months) and old (20-24 months) FACS- purified activated NSCs used for generating ATAC-seq libraries from freshly isolated cells from the SVZ neurogenic niche. *n* = 8 young male and female GFAP-GFP mice and *n* = 11 old male and female GFAP-GFP mice combined over 2 experiments. Data normalized to young mean of each individual experiment. Boxplots display median, and lower and upper quartiles. Dots represent mean fluorescence intensity for an individual animal. *P*-values were calculated using a two-tailed Mann-Whitney test. Data from independent experiments are in Source Data. e, Insert size distribution histograms of all paired-end reads from a single representative ATAC-seq library for each of the 5 cell types freshly isolated from the SVZ of both young and old animals. f, Heatmap of ATAC-seq enrichment at genome-wide transcription start sites (TSSs) ± 2000 base pairs for all 5 cell types in both young and old conditions.

**Extended Data Figure 2:**
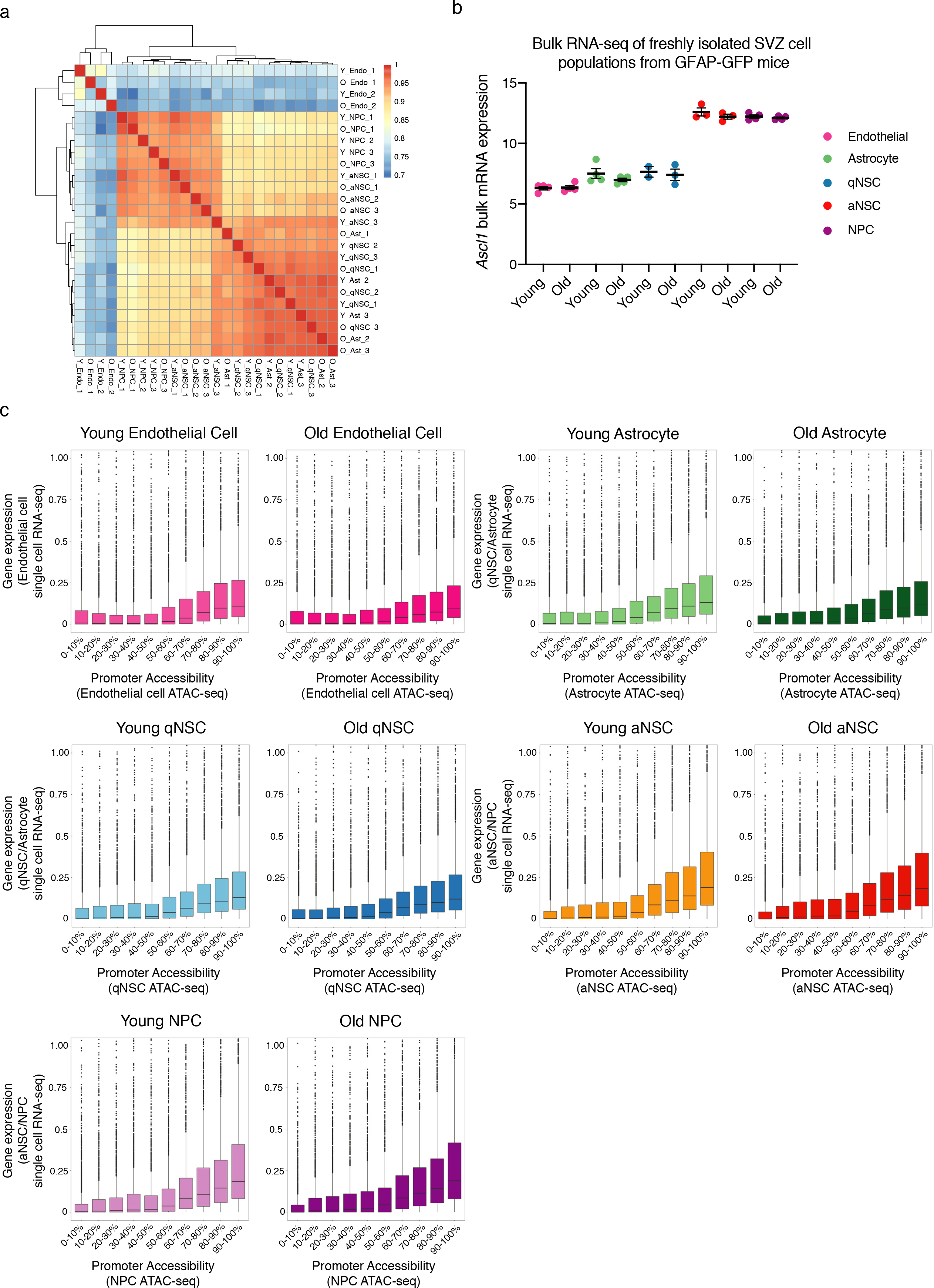
Chromatin accessibility clustering and correlation with gene expression. a, Hierarchical clustering using Pearson’s correlation of raw chromatin accessibility libraries (un- normalized) of all 5 cell types freshly isolated from the SVZ of young and old animals. Biological replicates are noted as numbers following age and cell type. b, mRNA expression of *Ascl1* in all 5 SVZ cell types in young and old conditions from bulk RNA-seq dataset (see Methods). Each dot represents the VST-normalized mRNA expression value from a single RNA-seq library (2-4 libraries per condition). Data are mean ± SEM. c, Decile plot of correlation between promoter accessibility levels (from ATAC-seq data) and mean gene expression values (from single cell RNA-seq data, see Methods). Promoters were binned into deciles based on accessibility level and boxplots were generated to correlate promoter bins with average gene expression for young and old endothelial cells, astrocytes, qNSCs, aNSCs, and NPCs. Each dot represents the gene expression level of an individual promoter binned based on chromatin accessibility signal. Y-axis constrained between 0 and 1 to facilitate boxplot visualization. Boxplot indicates median, and lower and upper quartiles.

**Extended Data Figure 3:**
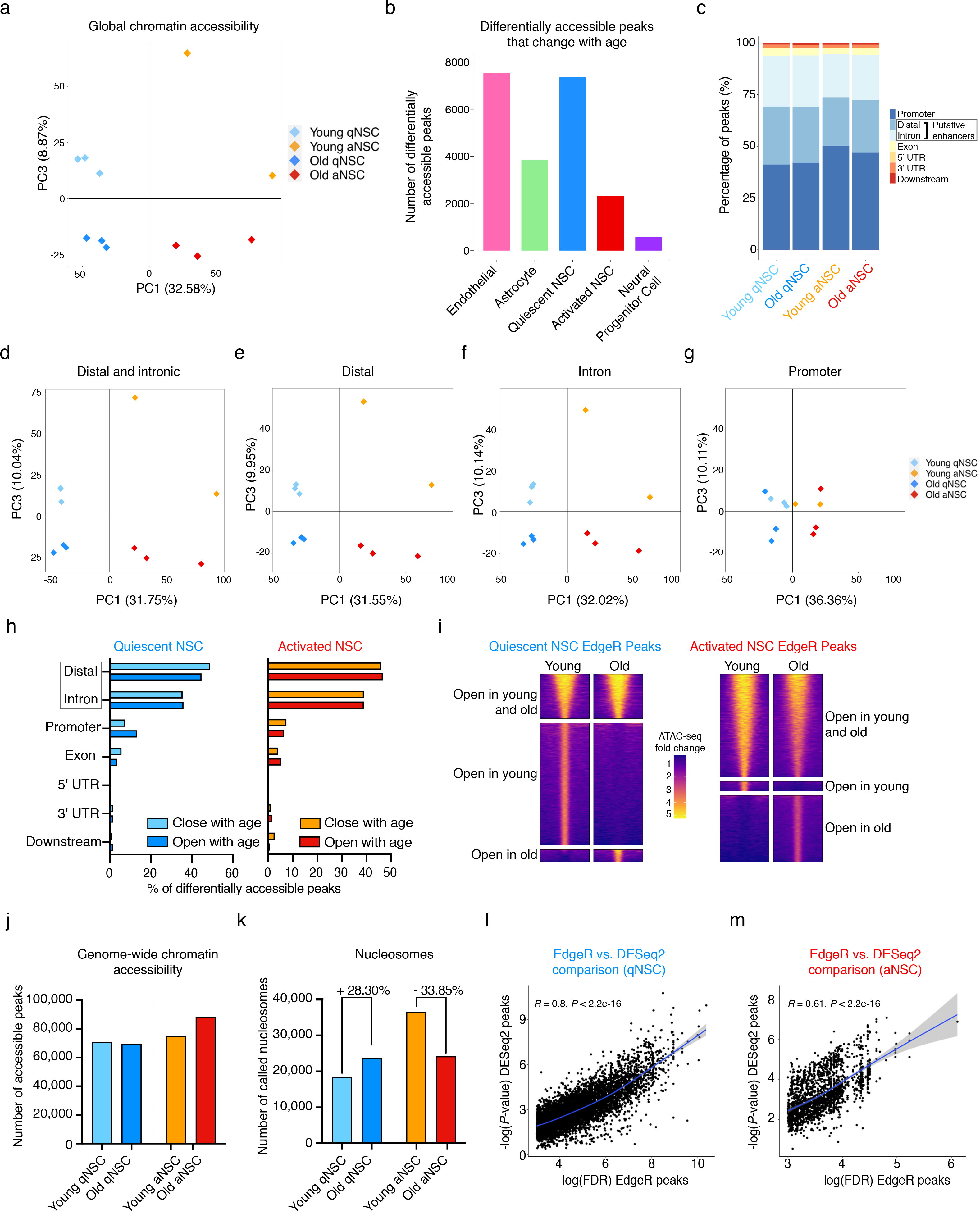
Chromatin accessibility in quiescent and activated NSCs during aging. a, PCA on genome-wide chromatin peaks defining qNSCs and aNSCs freshly isolated from the young and old SVZ where each dot represents a single ATAC-seq library (PC1 vs. PC3). PCA was generated using the variance stabilizing transformation (VST)-normalized NSC consensus count matrix. PC, principal component. b, Barplot denoting the number of differentially accessible peaks that change with age (open and close) in freshly isolated endothelial cells, astrocytes, quiescent NSCs, activated NSCs, and neural progenitor cells. c, Stacked barchart representing the genomic distribution of open chromatin peaks for qNSCs and aNSCs in young and old conditions based on their pooled peaksets. ATAC-seq peaks were annotated with ChIPSeeker (v1.18.0). Grey box indicates distal and intronic peaks that likely contain putative enhancers. d-g, PCA on distal and intronic chromatin peaks (d), exclusively distal (e), exclusively intronic (f), and exclusively promoter (g) chromatin peaks for qNSCs and aNSCs freshly isolated from the SVZ of young and old mice where each dot represents a single ATAC-seq library (PC1 vs. PC3). PCA was generated using the variance stabilizing transformation (VST)-normalized NSC consensus count matrix. PC, principal component. h, Frequency of differentially accessible ATAC-seq peaks that change with age in qNSCs (left) and aNSCs (right) associated with distal regions, introns, promoters, exons, 5’ UTRs, 3’ UTRs, and downstream regions. ATAC-seq peaks were annotated with ChIPSeeker (v1.18.0). Grey box denotes differentially accessible distal and intronic peaks that likely contain putative enhancers. i, Raw signal pileup plots illustrating chromatin accessibility signal of quiescent (left) and activated (right) NSC chromatin peaks separated based on whether they are common to qNSCs and aNSCs (top), open in young (middle), or open in old (bottom) as determined by the differential peak caller EdgeR. j, Barplot denoting the number of genome-wide chromatin peaks (non-differential) in freshly isolated young and old qNSCs and aNSCs. k, Number of nucleosome peaks called by NucleoATAC (v0.2.1) for freshly isolated young and old qNSC and aNSC chromatin landscapes. Nucleosome peaks were called using pooled, sub-sampled (to 30 million unique reads) ATAC-seq reads for each condition. l, Scatterplot illustrating correlation between significance levels of alternative differential peak calling methods: EdgeR FDR values vs DESeq2 P-values for peaks that are differentially open in young quiescent NSCs (called by EdgeR). Regression and *P*-values were respectively calculated using geom_smooth() and state_cor() in R. m, Scatterplot illustrating correlation between significance levels of alternative differential peak calling methods: EdgeR FDR values vs DESeq2 P-values for peaks that are differentially open in old activated NSCs (called by EdgeR). Regression and *P*-values were respectively calculated using geom_smooth() and state_cor() in R.

**Extended Data Figure 4:**
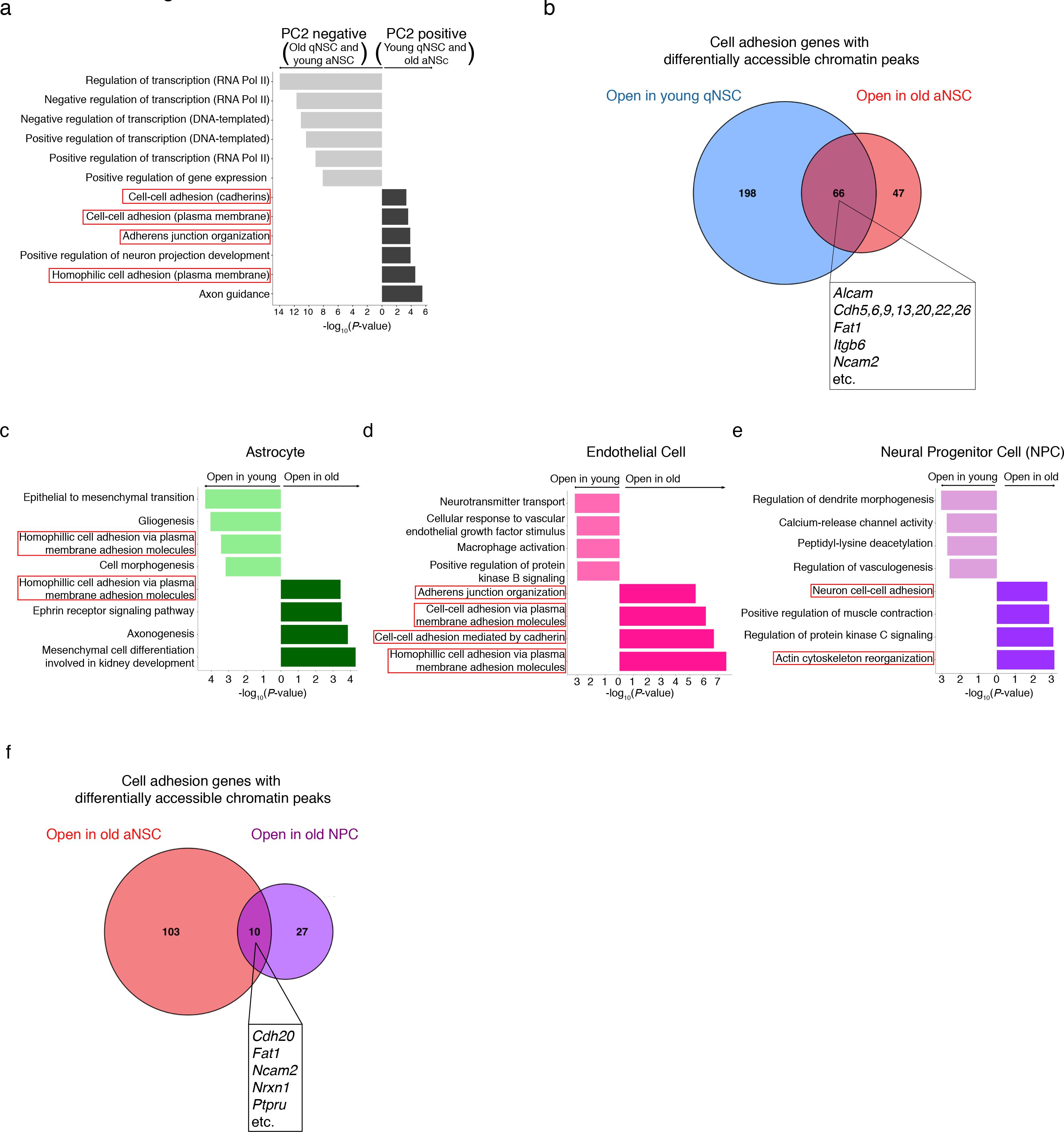
Chromatin accessibility analysis in 5 different cell types. a, Top 6 GO terms associated with the top 1000 ATAC-seq peak-associated genes driving PC2 in the positive (young qNSCs and old aNSCs) and negative (old qNSCs and young aNSCs) direction. Red boxes indicate GO terms associated with cell adhesion. PC, principal component. b, Venn diagram illustrating overlap between genes in “Cell Adhesion” GO category (GO:0007155) with nearby chromatin peaks that are differentially open in young qNSCs (compared to old) and old aNSCs (compared to young). Box in overlap indicates select shared cell adhesion genes. c-e, Selected GO terms associated with differentially accessible ATAC-seq peaks (FDR<0.05) that change with age in freshly isolated astrocytes (c), endothelial cells (d), and neural progenitor cells (NPCs) (e) generated by EnrichR and ranked by *P*-value. ATAC-seq peaks were annotated with their nearest gene using ChIPSeeker (v1.18.0). Red boxes indicate GO terms associated with adhesion and migration. f, Venn diagram illustrating overlap between genes in “Cell Adhesion” GO category (GO:0007155) genes with nearby chromatin peaks that are differentially open in old aNSCs and old NPCs. Box in overlap indicates select shared cell adhesion genes.

**Extended Data Figure 5:**
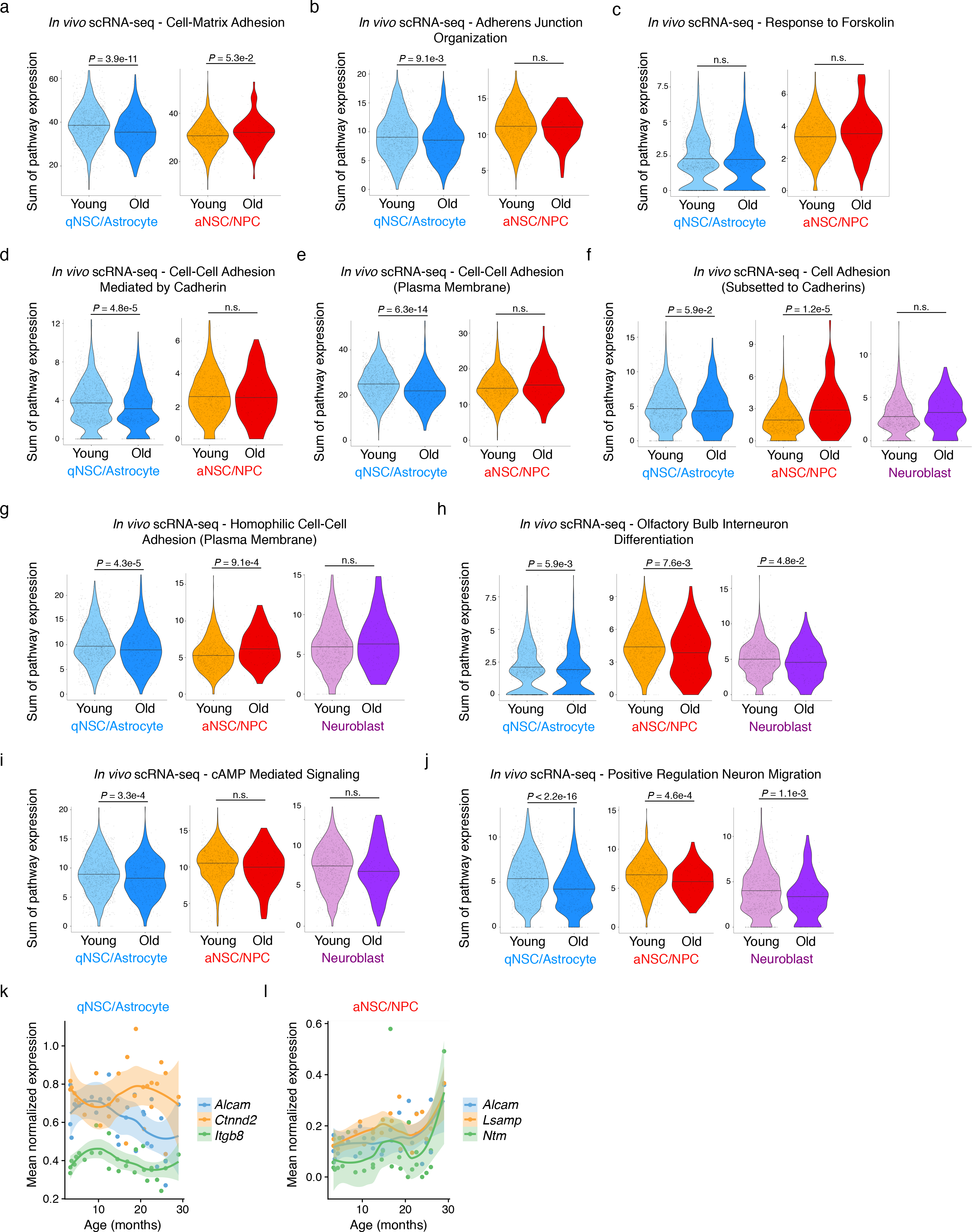
Gene expression of cell adhesion and migration pathways in NSC lineage during aging by single cell RNA-seq. a-e, Violin plots of the distribution of single cell cumulative expression profiles of genes within the “Cell-matrix adhesion” GO category (GO:0007160) (a), “Adherens junction organization” GO category (GO:0034332) (b), “Response to forskolin” GO Category (GO:1904322) (c), “Cell-cell adhesion mediated by cadherin” GO Category (GO:0044331) (d), and “Cell-cell adhesion (plasma membrane)” GO Category (GO:0098742) (e) for young and old qNSCs/Astrocytes (left) and aNSCs/NPCs (right) from single cell RNA-seq data (see Methods). Each overlaid dot represents the cumulative expression of genes within the GO category in a single cell. *P*-values were calculated using a two-tailed Mann-Whitney test. f-j, Violin plots of the distribution of single cell cumulative expression profiles of genes within the “Cell adhesion” GO Category (GO:0007155) restricted to cadherins (f), “Homophilic cell-cell adhesion (plasma membrane)” GO Category (GO:0007156) (g), “Olfactory bulb interneuron differentiation” GO Category (GO:0021889) (h), “cAMP mediated signaling” GO Category (GO:0019933) (i), and “Positive regulation neuron migration” GO Category (GO:2001224) (j) for young and old qNSC/Astrocyte (left), aNSC/NPC (middle), and neuroblasts (right) from single cell RNA-seq data (see Methods). Each overlaid dot represents the cumulative expression of genes within the GO category in a single cell. *P*-values were calculated using a two- tailed Mann-Whitney test. k, Gene expression trajectories as a function of age of select cell adhesion genes (*Alcam, Ctnnd2, Itgb8*) in qNSCs/Astrocytes from single cell RNA-seq dataset (see Methods). l, Gene expression trajectories as a function of age of select cell adhesion genes (*Alcam, Lsamp, Ntm*) in aNSCs/NPCs from single cell RNA-seq dataset (see Methods).

**Extended Data Figure 6:**
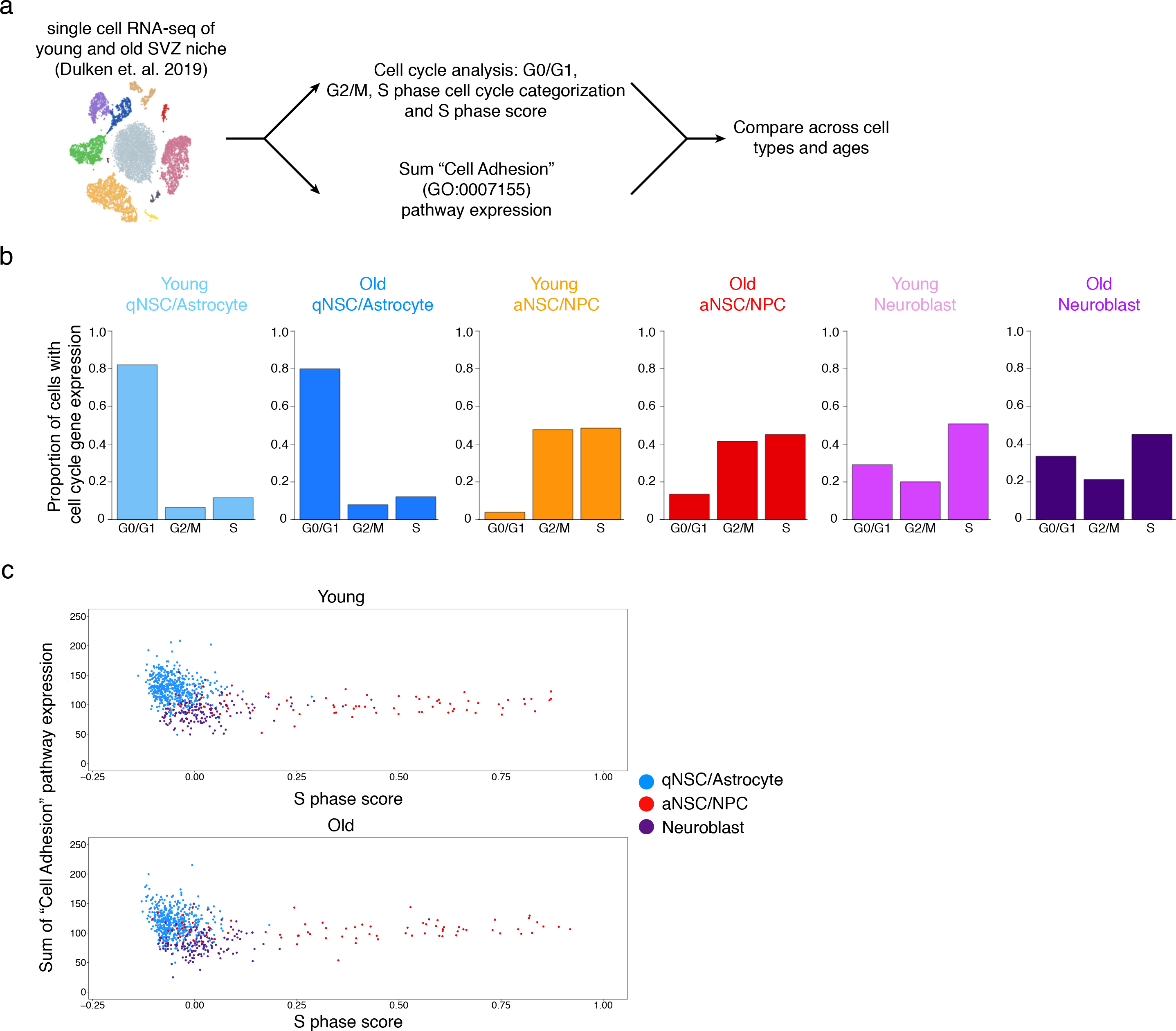
Analysis of cell cycle and cell migration signatures in NSCs during aging by single cell RNA-seq. a, Single cell RNA-seq data was used to assess cell cycle gene expression and Cell Adhesion (GO:0007155) pathway expression for young and old qNSCs/astrocytes, aNSCs/NPCs, and neuroblasts (see Methods). b, Barplots illustrating the proportion of young and old qNSCs/astrocytes, aNSCs/NPCs, and neuroblasts in the GO/G1, G2/M, or S phase of the cell cycle based on gene expression signatures from single cell RNA-seq data (see Methods). c, Scatterplot of qNSCs/astrocytes, aNSCs/NPCs, and neuroblasts from young (top) and old (bottom) single cell RNA- seq data (see Methods) illustrating the relationship between Seurat’s “S phase score” and the cumulative single cell expression levels of the genes in the “Cell Adhesion” GO category (GO:0007155). Young cells were downsampled to visualize the same number of young and old cells per cell type (480 qNSCs/astrocytes, 82 aNSCs/NPCs, and 146 neuroblasts).

**Extended Data Figure 7:**
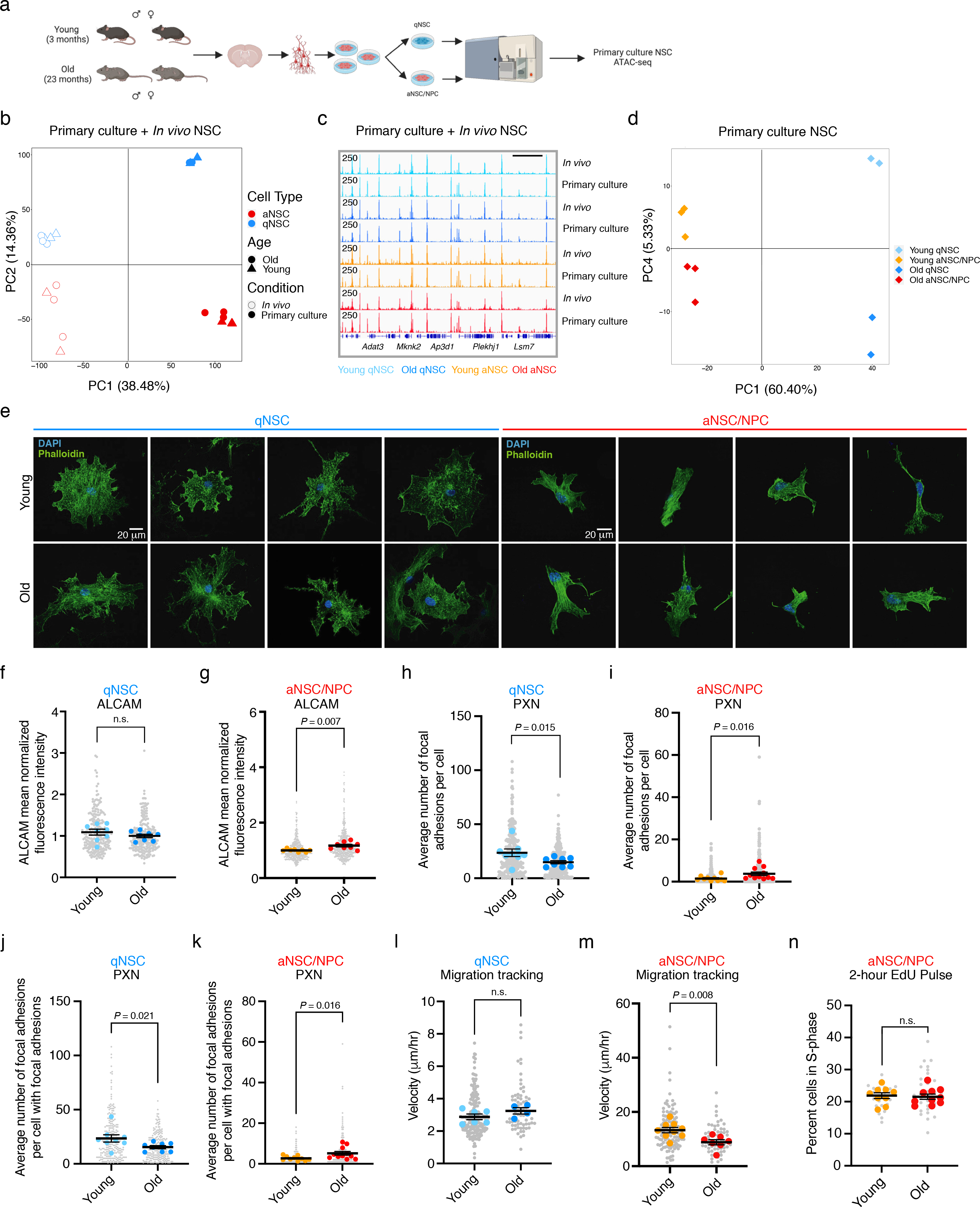
Primary cultures of qNSCs and aNSCs/NPCs for ATAC-seq, immunostaining, and adhesion and migration assays. a, Experimental design for the generation of ATAC-seq libraries from quiescent and activated NSCs cultured from young (3 months) and old (23 months) mixed-sex C57BL/6 mice. For each library, fluorescence-activated cell sorted (FACS) was used to sort 10,000-15,000 live cells from cultures generated from mixed male and female C57BL/6 SVZs (see Methods). b, PCA on chromatin peaks defining cultured and *in vivo* qNSC and aNSC landscapes where each dot represents a single ATAC- seq library (PC1 vs. PC2). PCA was generated using the variance stabilizing transformation (VST)- normalized global consensus count matrix of all cultured and *in vivo* NSC peaks. PC, principal component. c, Genome browser (IGV) view of chromatin accessibility signal tracks on chromosome 10 of freshly isolated and cultured qNSC and aNSC from the young and old SVZ. *Adat3*, Adenosine Deaminase TRNA Specific 3. *Mknk2*, MAPK Interacting Serine/Threonine Kinase 2. *Ap3d1*, Adaptor Related Protein Complex 3 Subunit Delta 1. *Plekhj1*, Pleckstrin Homology Domain Containing J1. *Lsm7*, LSM7 Homology U6 Small Nuclear RNA and mRNA Degradation Associated. Scale bar, 50kb. d, PCA on chromatin peaks defining qNSC and aNSC/NPC cultured from young and old SVZs where each dot represents a single ATAC-seq library (PC1 vs. PC4). PCA was generated using the variance stabilizing transformation (VST)-normalized global consensus count matrix. PC, principal component. e, Representative immunofluorescent images of young (2.5-3 months) (top) and old (21-22.1 months) (bottom) of aNSC/NPC (left) and qNSC (right) highlighting heterogeneity in cell size and shape. Green, phalloidin (F-actin); Blue, DAPI (nuclei). Scale bar, 20 μm. f, g, ALCAM fluorescence intensity of young (2.5-3 months) and old (21-22.1 months) qNSC (f) and aNSC/NPC (g) on Matrigel. Each grey dot represents mean normalized fluorescent intensity per cell. Each colored dot represents mean normalized fluorescence intensity of 30 fields (each containing 1-3 cells) in a primary culture derived from an individual mouse, normalized by experiment and cell size. *n* = 8 young male mice, and *n* = 8 old male mice, combined over 2 (qNSC) or 3 (aNSC/NPC) independent experiments. Data are mean ± SEM. *P*-values were calculated using a two-tailed Mann-Whitney test comparing sample means. Data from independent experiments are in Source Data. h,i, Quantification of paxillin (PXN) immunostaining of young (2.5-3 months) and old (22.1-22.1 months) qNSCs (h) and aNSCs/NPCs (i) cultured on Matrigel. Each grey dot represents number of focal adhesions per cell. Each colored dot represents average number of focal adhesions per cell (28-32 cells per dot) in a primary culture derived from an individual mouse. *n* = 8 young male mice, and *n* = 8 old male mice (qNSCs), *n* = 10 young male mice, and *n* = 11 old male mice (aNSCs/NPCs), combined over 2 (qNSCs) or 3 (aNSCs/NPCs) independent experiments. Data are mean ± SEM. *P*-values were calculated using a two-tailed Mann- Whitney test comparing sample means. Data from independent experiments are in Source Data. j,k, Quantification of subpopulation of cells with at least one active focal adhesion as assessed with paxillin (PXN) immunostaining of young (2.5-3 months) and old (22.1-22.1 months) qNSCs (j) and aNSCs/NPCs (k) cultured on Matrigel. Each grey dot represents number of focal adhesions per cell. Each colored dot represents average number of focal adhesions per cell of cells that have at least once focal adhesion (4-23 cells per dot) from an individual mouse. *n* = 8 young male mice, and *n* = 8 old male mice (qNSCs), *n* = 10 young male mice and *n* = 11 old male mice (aNSCs/NPCs), combined over 2 (qNSCs) or 3 (aNSCs/NPCs) independent experiments. Same experiment as in (h) and (i). Data are mean ± SEM. *P*-values were calculated using a two-tailed Mann-Whitney test comparing sample means. Data from independent experiments are in Source Data. l, Migration speed of young (3-4 months) and old (23-24 months) qNSCs on Poly-D-Lysine. Each grey dot represents the average velocity of a single cell over a 20-hour period. Each dot represents the average velocity over a 20-hour period of 5-42 cells in a primary culture derived from an individual mouse. *n* = 6 young male mice and *n* = 4 old male mice, combined over 2 independent experiments. Data are mean ± SEM. *P*-values were calculated using a two-tailed Mann-Whitney test comparing sample means. Data from independent experiments are in Source Data. m, Migration speed of young (3-4 months) and old (21-24 months) aNSCs/NPCs migrating on Poly-D-Lysine. Each grey dot represents the average velocity of a single cell over a 20-hour period. Each colored dot represents the average velocity per mouse over a 20-hour period of 6-28 cells. *n* = 9 young male mice and *n* = 7 old male mice, combined over 3 independent experiments. Data are mean ± SEM. *P*-values were calculated using a two-tailed Mann-Whitney test comparing sample means. Data from independent experiments are in Source Data. n, Percent cells that are EdU-positive (S-phase) after a 2-hour pulse for young (2.5-4 months) and old (21-23 months) aNSCs/NPCs on Poly-D-Lysine. Each grey dot represents percent of cells in given field that are EdU- positive. Each colored dot represents average percent cells EdU-positive for an individual mouse (average of 5-7 fields containing at least 100 cells each). *n =* 9 young male mice, and *n =* 10 old male mice, combined over 3 experiments. Data are mean ± SEM. *P*-values were calculated using a two-tailed Mann-Whitney test comparing sample means. Data from independent experiments are in Source Data.

**Extended Data Figure 8:**
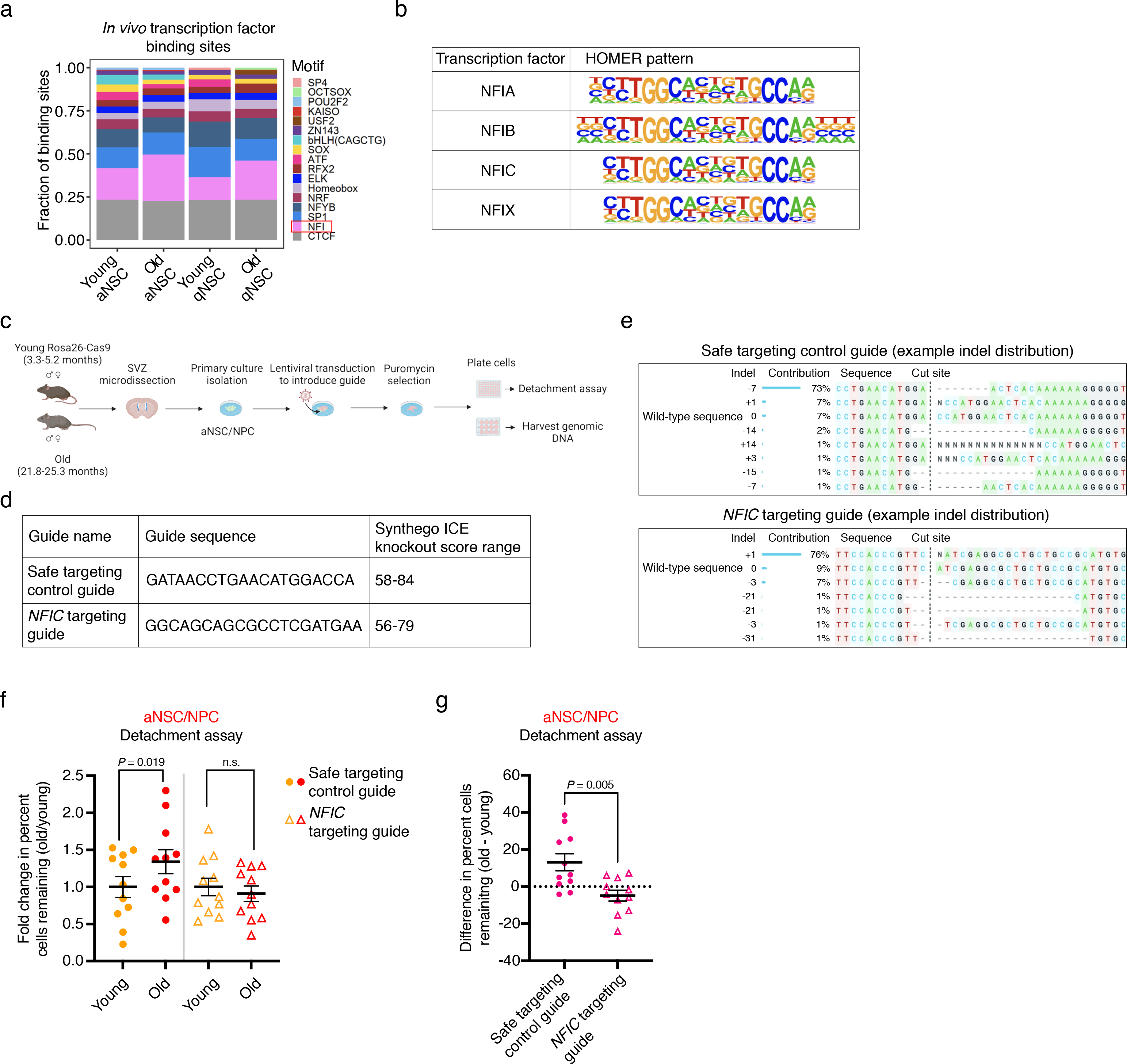
NFI family of transcription factors during aging in activated NSCs. a, Distribution of transcription factor binding sites in accessible chromatin regions from freshly isolated young and old aNSCs and qNSCs identified using a deep learning model (see Methods). b, NFI family transcription factor binding site motifs for each NFI isoform from HOMER. c, Experimental pipeline for knocking out *NFIC* in primary cultures of young and old aNSCs/NPCs. Primary cultures derived from young and old Rosa26-Cas9 mixed male and female mice were infected with lentivirus to deliver guides of interest, puromycin selected, and then used for detachment assay and for harvesting genomic DNA to assess knockout efficiency (see Methods). d, Guide sequences used for safe targeting control guide which targets a safe harbor locus (gene that can be modified or disrupted with CRISPR-Cas9 without any adverse effect on the cell) and *NFIC* targeting guide from Bassik Lab mouse CRISPR Knockout library (see Methods) as well as Synthego ICE knockout score range achieved with each guide. With this tool, a score of 50 indicates 50% of the population sequenced was unedited (contains wild-type sequence). e, Sample traces for safe targeting control guide and *NFIC* targeting guide from Synthego ICE analysis tool indicating indel distribution and cut site. f, Detachment assay upon *NFIC* knockout in young and old aNSC/NPC using *NFIC* targeting guide. Quantification of percent of aNSCs/NPCs remaining after 3 minute Accutase incubation and wash with 1x PBS, normalized to young average from paired experiment. Each data point is average of 2-3 technical replicates per primary culture derived from an individual mouse. Safe targeting control guide samples (solid dot) were normalized to average of all young safe targeting control samples. *NFIC* targeting guide samples (open triangle) were normalized to average of all young *NFIC* targeting guide samples. Cells imaged and analyzed using Incucyte S3 system. Young (3.33-5.2 months) and old (21.8-25.3 months) aNSCs/NPCs were infected with lentivirus containing safe targeting control guide control or *NFIC* targeting guide. *n* = 11 young male and female mice, and *n* = 11 old male and female mice, combined over 7 experiments. Data are mean ± SEM. *P*-values were calculated using a Wilcoxon Matched-Pairs Signed Ranks Test, since experiments were performed in a paired manner (paired young and old). g, Detachment assay upon *NFIC* knockout in young and old aNSCs/NPCs using *NFIC* targeting guide. Same experiment as in (f). Quantification of difference of percent of aNSCs/NPCs remaining (old – young) after 3 minute Accutase incubation and wash with 1x PBS for safe targeting control guide samples (solid dot) and *NFIC* targeting guide samples (open triangle). Cells imaged and analyzed using Incucyte S3 system. Young (3.33-5.2 months) and old (21.8-25.3 months) aNSCs/NPCs were infected with lentivirus containing safe targeting control guide control or *NFIC* targeting guide. *n* = 11 young male and female mice, and *n* = 11 old male and female mice, combined over 7 experiments. Data are mean ± SEM. *P*-value was calculated using a two-tailed Mann-Whitney test.

**Extended Data Figure 9:**
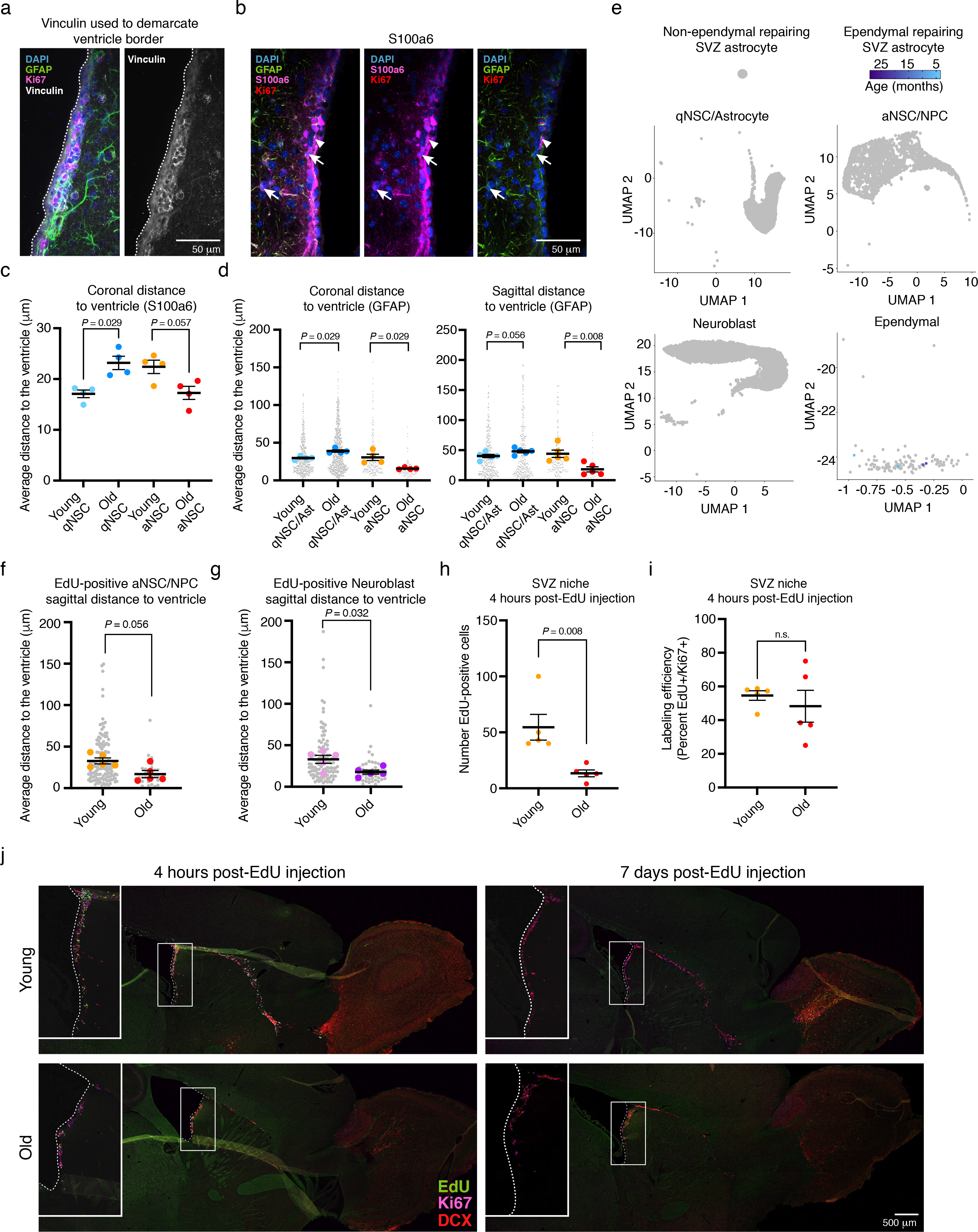
*In vivo* location of EdU-labelled cells in the SVZ neurogenic niche. a, Representative image of vinculin immunofluorescence staining of old coronal section from GFAP- GFP mouse (Fig. 3c) indicating how vinculin (white) was used to demarcate ventricle border. The ventricular lining is indicated by a dotted white line. Green, GFAP (astrocyte/NSC marker); Pink, Ki67 (proliferation marker); Blue, DAPI (nuclei). Scale bar, 50 μm. b, Representative image of S100a6 immunofluorescent staining of old coronal section from GFAP-GFP mouse. White arrows: qNSC (S100a6+/Ki67- and GFAP+/Ki67-); white arrowheads aNSC (S100a6+/Ki67+ and GFAP+/Ki67+). Green, GFAP (astrocyte/NSC marker); Pink, S100a6 (NSC marker in the SVZ neurogenic niche); Red, Ki67 (proliferation marker); Blue, DAPI (nuclei). Scale bar, 50 μm. c, NSC distance to the ventricle was calculated for qNSCs (S100a6+/Ki67-) and aNSCs (S100a6+/Ki67+) in coronal sections of young (4 months) and old (18-26 months) SVZs. Each dot represents the mean distance from the ventricle of 140 – 574 cells in 6-11 fields per section (3 sections per dot) for an individual mouse. *n* = 4 mixed-sex young mice, and *n* = 4 mixed-sex old mice, combined over 4 independent experiments. Data are mean ± SEM. *P*-values were calculated using a two-tailed Mann-Whitney test comparing sample means. Data from independent experiments are in Source Data. d, NSC distance to the ventricle was calculated for qNSCs/astrocytes (GFAP+/Ki67-) and aNSCs (GFAP+/Ki67+) in coronal sections (left) of young (4 months) and old (18-26 months) SVZs from mixed sex GFAP-GFP mice and sagittal sections (right) of young (2-3 months) and old (21 months) SVZs from male C57BL/6 mice. Each grey dot represents the distance of a single cell from the ventricular lining. Each colored dot represents the mean distance from the ventricle of 19 - 231 cells in 3-7 fields per section (3 sections for coronal sections or 1 section for sagittal section) per individual mouse. Coronal sections: *n* = 4 young, and *n* = 4 old mixed-sex GFAP-GFP mice, combined over 4 independent experiments. Sagittal sections: *n* = 5 young male mice, and *n* = 5 old male C57BL/6 mice combined over 2 independent experiments. Data are mean ± SEM. *P*-values were calculated using a two-tailed Mann-Whitney test comparing sample means. Data from independent experiments are in Source Data. e, Single cell RNA-seq analysis of 28 C57BL/6 mice ranging in age from 3.3 months to 29 months (see Methods). Out of 21,458 cells, there are 4 cells expressing all markers of ependymal-repairing SVZ astrocytes (*S100b, Gfap, CD24a, Ctnnb1*) (see Methods). These cells are colored by age (4.7 months, 5.4 months, 16.83 months, and 18.87 months). Ependymal-repairing SVZ astrocytes are absent from qNSC/astrocyte, aNSC/NPC, and neuroblast clusters but are present in ependymal cell cluster. f, Distance to the ventricle was calculated for EdU+ aNSC/NPC (Ki67+/DCX-) in sagittal sections of young (2-3 months) and old (21 months) SVZs 4 hours after EdU injection. Each grey dot represents the distance of a single aNSC/NPC from the ventricle. Each colored dot represents the mean aNSC/NPC distance from the ventricle of 2-35 cells from 1 section per individual mouse. *n* = 5 young male mice, and *n* = 5 old male mice, combined over 2 experiments. Data are mean ± SEM. *P*-values were calculated using a two-tailed Mann-Whitney test comparing sample means. Data from independent experiments are in Source Data. g, Distance to the ventricle was calculated for EdU+ neuroblasts (DCX+) in sagittal sections of young (2-3 months) and old (21 months) SVZs 4 hours after EdU injection. Each grey dot represents the distance of a single neuroblast from the ventricle. Each colored dot represents the mean neuroblast distance from the ventricle of 3-25 cells from 1 section per individual mouse. *n* = 5 young male mice, and *n* = 5 old male mice, combined over 2 experiments. Data are mean ± SEM. *P*-values were calculated using a two-tailed Mann-Whitney test comparing sample means. Data from independent experiments are in Source Data. h, Quantification of number of EdU+ cells counted in the SVZ (along the entire length of the ventricle) from young (2-3 months) and old (21 months) sagittal sections 4 hours post-injection of EdU. *n* = 5 young male mice, and *n* = 5 old male mice, combined over 2 experiments. Data are mean ± SEM. *P*-values were calculated using a two-tailed Mann-Whitney test. Data from independent experiments are in Source Data. i, Quantification of percent of Ki67+ cells that are EdU+ 4 hours post- injection of EdU in the SVZ (counting cells along the entire length of the ventricle). *n* = 5 young male mice, and *n* = 5 old male mice, combined over 2 experiments. Data are mean ± SEM. *P*-values were calculated using a two-tailed Mann-Whitney test. Data from independent experiments are in Source Data. j, Representative images of immunofluorescence staining of sagittal sections encompassing the SVZ, RMS, and OB regions from a young (2-3 months) (top) or old (21 months) (bottom) male C57BL/6 mouse 4hrs (left) or 7 days (right) after intraperitoneal EdU injection. White inset denotes SVZ. Dotted white line indicates ventricle lining. Green, EdU (proliferation marker); pink, Ki67 (aNSC/NPC/neuroblast marker); red, DCX (neuroblast marker). Scale bar, 500 μm.

**Extended Data Figure 10:**
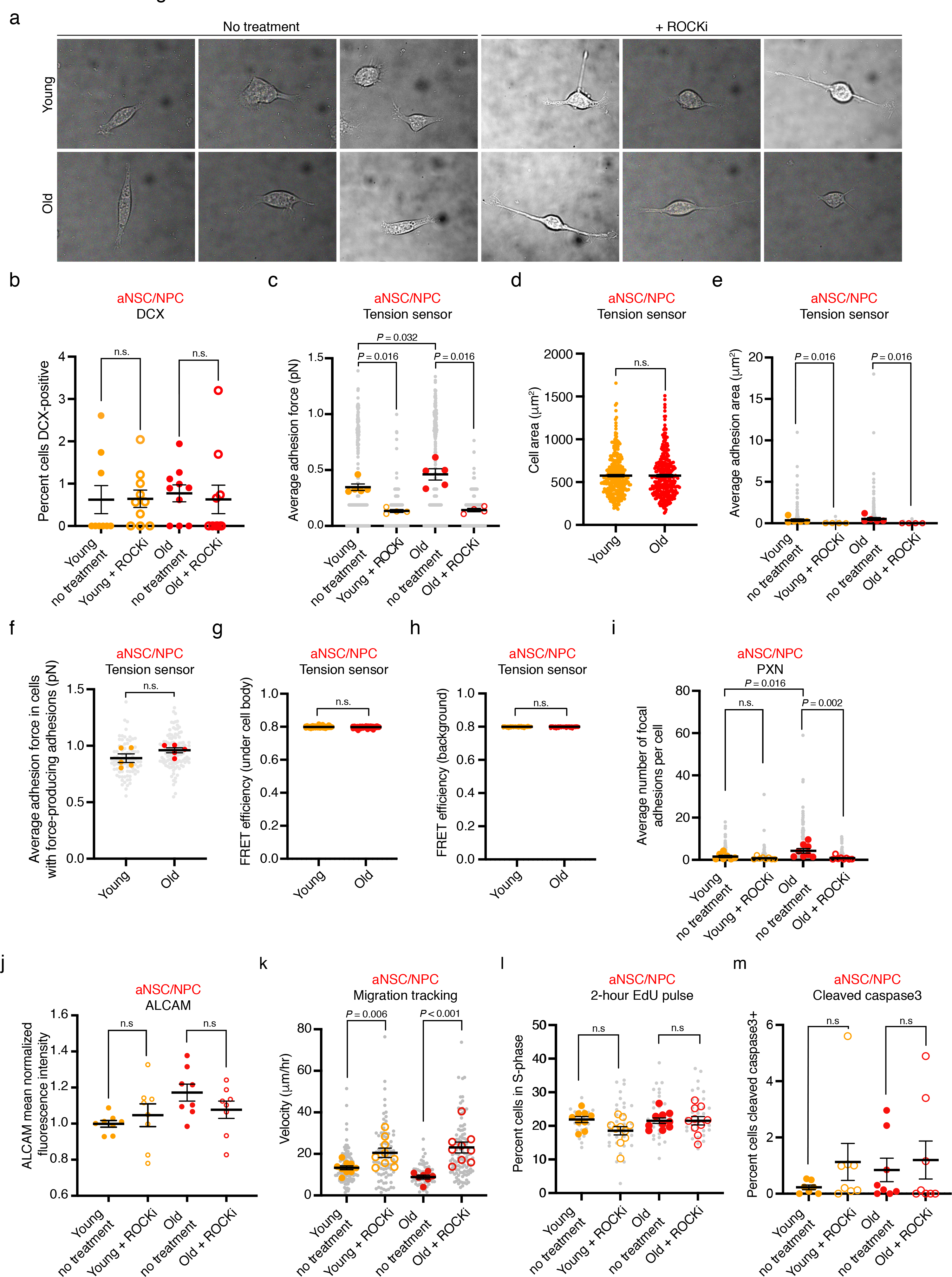
Effect of ROCK inhibitor *in vitro* and *in vivo*. a, Representative brightfield images of young (2.4-3 months) (top) and old (20.5-21 months) (bottom) aNSCs/NPCs on RGD molecular tension sensors with H2O vehicle (left) or 10 μM ROCKi (right). b, Percent cells that are positive for DCX staining in young (4 months) and old (22 months) aNSCs/NPCs treated with H2O vehicle (no treatment, solid dots) or 10 μM ROCKi (ROCKi treatment, open circles) on Poly-D-Lysine. Each dot represents percent cells DCX-positive in a given field (containing 38-349 cells) for an individual animal (4-5 fields per animal). *n =* 2 young male mice and *n* = 2 old male mice (no treatment), and *n =* 2 young male mice and *n* = 2 old male mice (+ROCKi), combined over 1 experiment. Data are mean ± SEM. *P*-values were calculated using a two-tailed Mann-Whitney. Data from independent experiments are in Source Data. c, Quantification of average adhesion force (pN) exhibited by young (2.4-3 months) and old (20.5-21 months) cultured aNSCs/NPCs treated with H2O vehicle (no treatment, solid dots) or 10 μM ROCKi (ROCKi treatment, open circles) determined using RGD molecular tension sensors. Each grey dot represents adhesion force produced by one cell. Each colored dot represents the average force produced by one cell (15– 89 cells per dot) in a primary culture derived from an individual mouse. *n* = 5 young male mice and *n* = 5 old male mice (no treatment), *n =* 4 young male mice and *n* = 4 old male mice (ROCKi treatment), combined over 5 independent experiments. Data are mean ± SEM. *P*-values were calculated using a two-tailed Mann- Whitney test comparing sample means. Data from independent experiments are in Source Data. d, Cell areas of young (2.4-3 months) and old (20.5-21 months) cultured aNSCs/NPCs determined using bright field images taken on RGD molecular tension sensors. *n* = 306 young aNSCs/NPCs, and *n* = 298 old aNSCs/NPCs, combined over 5 experiments. Data are mean ± SEM. *P*-values were calculated using a two-tailed Mann-Whitney test. Data from independent experiments are in Source Data. e, Quantification of average adhesion area under force-producing adhesions (μm^2^) of young (2.4-3 months) and old (20.5-21 months) cultured aNSCs/NPCs treated with H2O vehicle (no treatment, solid dots) or 10 μM ROCKi (ROCKi treatment, open circles) determined using RGD molecular tension sensors. Each grey dot represents adhesion area of force-producing adhesion from a single cell. Each colored dot represents the average adhesion area of force-producing adhesions from a single cell (15 – 89 cells per dot) in a primary culture derived from an individual mouse. Same experiment as in (c). Data are mean ± SEM. *P*-values were calculated using a two-tailed Mann-Whitney test comparing sample means. Data from independent experiments are in Source Data. f, Quantification of average adhesion force (pN) exhibited by young (2.4-3 months) and old (20.5-21 months) cultured aNSCs/NPCs that have at least one force-producing adhesion determined using RGD molecular tension sensors. Same experiment as in (c). Each grey dot represents adhesion force produced by one cell. Each colored dot represents the average force produced by one cell in cells that have at least one force-producing adhesion (8-29 cells per dot) in a primary culture derived from an individual mouse. *N* = 5 young male mice and *n* = 5 old male mice, combined over 5 independent experiments. Data are mean ± SEM. *P*-values were calculated using a two-tailed Mann-Whitney test comparing sample means. Data from independent experiments are in Source Data. g, FRET efficiency values underneath cell bodies of young (2.4-3 months) and old (20.5-21 months) cultured aNSC/NPC on RGD molecular tension sensors. *n* = 306 young aNSCs/NPCs, and *n* = 298 old aNSCs/NPCs, combined over 5 experiments. Same experiment as in (c). Data are mean ± SEM. *P*-values were calculated using a two- tailed Mann-Whitney test. Data from independent experiments are in Source Data. h, Background FRET efficiency values of young (2.4-3 months) and old (20.5-21 months) cultured aNSCs/NPCs on RGD molecular tension sensors. *n* = 306 young aNSCs/NPCs, and *n* = 298 old aNSCs/NPCs, combined over 5 experiments. Same experiment as in (c). Data are mean ± SEM. *P*-values were calculated using a two-tailed Mann-Whitney test. Data from independent experiments are in Source Data. i, Quantification of average number of focal adhesions, determined using paxillin (PXN) staining, exhibited by young (2.5 months) and old (22.1 months) cultured aNSCs/NPCs treated with H2O vehicle (no treatment, solid dots) or 10 μM ROCKi (ROCKi treatment, open circles). Each grey dot represents number of focal adhesions per cell. Each colored dot represents the average number of focal adhesions per cell from a primary culture (30 cells per dot) derived from an individual mouse. *N* = 7 young male mice and *n* = 8 old male mice (no treatment) and *n* = 8 young male mice and *n =* 8 old male mice (ROCKi treatment), combined over 2 experiments. Data are mean ± SEM. *P*-values were calculated using a two-tailed Mann-Whitney test comparing sample means. Data from independent experiments are in Source Data. j, Quantification of ALCAM fluorescence intensity of young (2.5-3 months) and old (21-22.1 months) aNSCs/NPCs on Matrigel treated with H2O vehicle (no treatment, solid dots) or 10 μM ROCKi (ROCKi treatment, open circles). Each dot represents mean-normalized fluorescence intensity of 30 fields (each containing 1-3 cells) in a primary culture derived from an individual mouse, normalized by experiment and cell size. *n* = 8 young male mice and *n* = 8 old mice (no treatment) and *n =*8 young male mice and *n* = 8 old male mice (ROCKi treatment), combined over 2 experiments. Data are mean ± SEM. *P*-values were calculated using a two-tailed Mann-Whitney test comparing sample means. Data from independent experiments are in Source Data. k, Quantification of migration speed of young (3-4 months) and old (21-24 months) aNSC/NPC cultured on Poly-D-Lysine ± treatment with 10 μM ROCKi. Each grey dots represents average velocity of a single cell over a 20- hour period. Each colored dot represents the average velocity over a 20-hour period of cells (2-28) from one culture derived from one individual mouse. *n* = 9 young male mice and *n* = 7 old male mice (no treatment, solid dots) and *n* = 9 young male mice and *n* = 9 old male mice (ROCKi treatment, open circles), combined over 3 experiments. Data are mean ± SEM. *P*-values were calculated using a two- tailed Mann-Whitney test comparing sample means. Data from independent experiments are in Source Data. l, Percent cells that are EdU-positive (S-phase) after a 2 hour pulse for young (2.5-4 months) and old (21-23 months) aNSCs/NPCs treated with H2O vehicle (no treatment, solid dots) or 10 μM ROCKi (ROCKi treatment, open circles) on Poly-D-Lysine. Each grey dot represents percent of cells in given field that are EdU-positive. Each colored dot represents average percent cells EdU-positive for an individual mouse (average of 5-7 fields containing at least 100 cells each). *n =* 9 young male mice and *n* = 10 old male mice (no treatment), and *n* = 10 young male mice and *n* = 10 old male mice (ROCKi treatment) combined over 3 experiments. Data are mean ± SEM. *P*-values were calculated using a two-tailed Mann-Whitney test comparing sample means. Data from independent experiments are in Source Data. m, Percent cells that are positive for cleaved-caspase 3 staining in young (3-4 months) and old (21-23 months) aNSCs/NPCs treated with H2O vehicle (no treatment, solid dots) or 10 μM ROCKi (ROCKi treatment, open circles) on Poly-D-Lysine. Each dot represents average percent cells cleaved caspase3-positive for an individual mouse (average of 5 fields containing at least 100 cells). *n =* 7 young male mice and *n* = 8 old male mice (no treatment), and *n =* 8 young male mice and *n* = 8 old male mice (+ROCKi), combined over 2 experiments. Data are mean ± SEM. *P*-values were calculated using a two-tailed Mann-Whitney test comparing sample means. Data from independent experiments are in Source Data.

## References

1. Bond, A.M., Ming, G.L. & Song, H. Adult Mammalian Neural Stem Cells and Neurogenesis: Five Decades Later. Cell Stem Cell 17, 385–95 (2015).

2. Navarro Negredo, P., Yeo, R.W. & Brunet, A. Aging and Rejuvenation of Neural Stem Cells and Their Niches. Cell Stem Cell 27, 202–223 (2020).

3. Gage, F.H. & Temple, S. Neural stem cells: generating and regenerating the brain. Neuron 80, 588–601 (2013).

4. Silva-Vargas, V., Crouch, E.E. & Doetsch, F. Adult neural stem cells and their niche: a dynamic duo during homeostasis, regeneration, and aging. Curr Opin Neurobiol 23, 935–42 (2013).

5. Denoth-Lippuner, A. & Jessberger, S. Formation and integration of new neurons in the adult hippocampus. Nat Rev Neurosci 22, 223–236 (2021).

6. Amano, M., Nakayama, M. & Kaibuchi, K. Rho-kinase/ROCK: A key regulator of the cytoskeleton and cell polarity. Cytoskeleton (Hoboken) 67, 545–54 (2010).

7. Ishizaki, T. et al. Pharmacological properties of Y-27632, a specific inhibitor of rho- associated kinases. Mol Pharmacol 57, 976–83 (2000).

8. Hattiangady, B., Rao, M.S. & Shetty, A.K. Plasticity of hippocampal stem/progenitor cells to enhance neurogenesis in response to kainate-induced injury is lost by middle age. Aging Cell 7, 207–24 (2008).

9. Conover, J.C. & Shook, B.A. Aging of the subventricular zone neural stem cell niche. Aging Dis 2, 49–63 (2011).

10. Nicaise, A.M., Willis, C.M., Crocker, S.J. & Pluchino, S. Stem Cells of the Aging Brain. Front Aging Neurosci 12, 247 (2020).

11. Mirzadeh, Z., Merkle, F.T., Soriano-Navarro, M., Garcia-Verdugo, J.M. & Alvarez- Buylla, A. Neural stem cells confer unique pinwheel architecture to the ventricular surface in neurogenic regions of the adult brain. Cell Stem Cell 3, 265–78 (2008).

12. Shen, Q. et al. Adult SVZ stem cells lie in a vascular niche: a quantitative analysis of niche cell-cell interactions. Cell Stem Cell 3, 289–300 (2008).

13. Tavazoie, M. et al. A specialized vascular niche for adult neural stem cells. Cell Stem Cell 3, 279–88 (2008).

14. Doetsch, F., Garcia-Verdugo, J.M. & Alvarez-Buylla, A. Cellular composition and three- dimensional organization of the subventricular germinal zone in the adult mammalian brain. J Neurosci 17, 5046–61 (1997).

15. Kernie, S.G. & Parent, J.M. Forebrain neurogenesis after focal Ischemic and traumatic brain injury. Neurobiol Dis 37, 267–74 (2010).

16. Faiz, M. et al. Adult Neural Stem Cells from the Subventricular Zone Give Rise to Reactive Astrocytes in the Cortex after Stroke. Cell Stem Cell 17, 624–34 (2015).

17. Capilla-Gonzalez, V., Cebrian-Silla, A., Guerrero-Cazares, H., Garcia-Verdugo, J.M. & Quinones-Hinojosa, A. Age-related changes in astrocytic and ependymal cells of the subventricular zone. Glia 62, 790–803 (2014).

18. Enwere, E. et al. Aging results in reduced epidermal growth factor receptor signaling, diminished olfactory neurogenesis, and deficits in fine olfactory discrimination. J Neurosci 24, 8354–65 (2004).

19. Luo, J., Daniels, S.B., Lennington, J.B., Notti, R.Q. & Conover, J.C. The aging neurogenic subventricular zone. Aging Cell 5, 139–52 (2006).

20. Tropepe, V., Craig, C.G., Morshead, C.M. & van der Kooy, D. Transforming growth factor-alpha null and senescent mice show decreased neural progenitor cell proliferation in the forebrain subependyma. J Neurosci 17, 7850–9 (1997).

21. Molofsky, A.V. et al. Increasing p16INK4a expression decreases forebrain progenitors and neurogenesis during ageing. Nature 443, 448–52 (2006).

22. Shi, Z. et al. Single-cell transcriptomics reveals gene signatures and alterations associated with aging in distinct neural stem/progenitor cell subpopulations. Protein Cell 9, 351–364 (2018).

23. Artegiani, B. et al. A Single-Cell RNA Sequencing Study Reveals Cellular and Molecular Dynamics of the Hippocampal Neurogenic Niche. Cell Rep 21, 3271–3284 (2017).

24. Dulken, B.W. et al. Single-cell analysis reveals T cell infiltration in old neurogenic niches. Nature 571, 205–210 (2019).

25. Kalamakis, G. et al. Quiescence Modulates Stem Cell Maintenance and Regenerative Capacity in the Aging Brain. Cell 176, 1407–1419 e14 (2019).

26. Klemm, S.L., Shipony, Z. & Greenleaf, W.J. Chromatin accessibility and the regulatory epigenome. Nat Rev Genet 20, 207–220 (2019).

27. Kane, A.E. & Sinclair, D.A. Epigenetic changes during aging and their reprogramming potential. Crit Rev Biochem Mol Biol 54, 61–83 (2019).

28. Gontier, G. et al. Tet2 Rescues Age-Related Regenerative Decline and Enhances Cognitive Function in the Adult Mouse Brain. Cell Rep 22, 1974–1981 (2018).

29. Benayoun, B.A. et al. Remodeling of epigenome and transcriptome landscapes with aging in mice reveals widespread induction of inflammatory responses. Genome Res 29, 697–709 (2019).

30. Berg, D.A. et al. A Common Embryonic Origin of Stem Cells Drives Developmental and Adult Neurogenesis. Cell 177, 654–668 e15 (2019).

31. Martynoga, B. et al. Epigenomic enhancer annotation reveals a key role for NFIX in neural stem cell quiescence. Genes Dev 27, 1769–86 (2013).

32. Lupo, G. et al. Molecular profiling of aged neural progenitors identifies Dbx2 as a candidate regulator of age-associated neurogenic decline. Aging Cell 17, e12745 (2018).

33. Maybury-Lewis, S.Y. et al. Changing and stable chromatin accessibility supports transcriptional overhaul during neural stem cell activation and is altered with age. Aging Cell 20, e13499 (2021).

34. Zhuo, L. et al. Live astrocytes visualized by green fluorescent protein in transgenic mice. Dev Biol 187, 36–42 (1997).

35. Codega, P. et al. Prospective identification and purification of quiescent adult neural stem cells from their in vivo niche. Neuron 82, 545–59 (2014).

36. Leeman, D.S. et al. Lysosome activation clears aggregates and enhances quiescent neural stem cell activation during aging. Science 359, 1277–1283 (2018).

37. Buenrostro, J.D., Giresi, P.G., Zaba, L.C., Chang, H.Y. & Greenleaf, W.J. Transposition of native chromatin for fast and sensitive epigenomic profiling of open chromatin, DNA- binding proteins and nucleosome position. Nat Methods 10, 1213–8 (2013).

38. Ucar, D. et al. The chromatin accessibility signature of human immune aging stems from CD8(+) T cells. J Exp Med 214, 3123–3144 (2017).

39. Shcherbina, A. et al. Dissecting Murine Muscle Stem Cell Aging through Regeneration Using Integrative Genomic Analysis. Cell Rep 32, 107964 (2020).

40. Koohy, H. et al. Genome organization and chromatin analysis identify transcriptional downregulation of insulin-like growth factor signaling as a hallmark of aging in developing B cells. Genome Biol 19, 126 (2018).

41. Ge, Y. et al. The aging skin microenvironment dictates stem cell behavior. Proc Natl Acad Sci U S A 117, 5339–5350 (2020).

42. Guillemot, F. & Hassan, B.A. Beyond proneural: emerging functions and regulations of proneural proteins. Curr Opin Neurobiol 42, 93–101 (2017).

43. Buckley, M.T. et al. Cell type-specific aging clocks to quantify aging and rejuvenation in regenerative regions of the brain. bioRxiv (2022).

44. Mira, H. et al. Signaling through BMPR-IA regulates quiescence and long-term activity of neural stem cells in the adult hippocampus. Cell Stem Cell 7, 78–89 (2010).

45. Jones, K.M. et al. CHD7 maintains neural stem cell quiescence and prevents premature stem cell depletion in the adult hippocampus. Stem Cells 33, 196–210 (2015).

46. Knobloch, M. et al. A Fatty Acid Oxidation-Dependent Metabolic Shift Regulates Adult Neural Stem Cell Activity. Cell Rep 20, 2144–2155 (2017).

47. Swart, G.W. Activated leukocyte cell adhesion molecule (CD166/ALCAM): developmental and mechanistic aspects of cell clustering and cell migration. Eur J Cell Biol 81, 313–21 (2002).

48. Masedunskas, A. et al. Activated leukocyte cell adhesion molecule is a component of the endothelial junction involved in transendothelial monocyte migration. FEBS Lett 580, 2637–45 (2006).

49. Lunter, P.C. et al. Activated leukocyte cell adhesion molecule (ALCAM/CD166/MEMD), a novel actor in invasive growth, controls matrix metalloproteinase activity. Cancer Research 65, 8801–8 (2005).

50. Schaller, M.D. Paxillin: a focal adhesion-associated adaptor protein. Oncogene 20, 6459–72 (2001).

51. Hu, Y.L. et al. FAK and paxillin dynamics at focal adhesions in the protrusions of migrating cells. Scientific Reports 4, 6024 (2014).

52. Brown, M.A. et al. The use of mild trypsinization conditions in the detachment of endothelial cells to promote subsequent endothelialization on synthetic surfaces. Biomaterials 28, 3928–35 (2007).

53. Loffek, S. et al. Transmembrane collagen XVII modulates integrin dependent keratinocyte migration via PI3K/Rac1 signaling. PLoS One 9, e87263 (2014).

54. Mizrahi, A., Lu, J., Irving, R., Feng, G. & Katz, L.C. In vivo imaging of juxtaglomerular neuron turnover in the mouse olfactory bulb. Proc Natl Acad Sci U S A 103, 1912–7 (2006).

55. Mobley, A.S. et al. Age-dependent regional changes in the rostral migratory stream. Neurobiol Aging 34, 1873–81 (2013).

56. Shuboni-Mulligan, D.D. et al. In vivo serial MRI of age-dependent neural progenitor cell migration in the rat brain. Neuroimage 199, 153–159 (2019).

57. Capilla-Gonzalez, V., Cebrian-Silla, A., Guerrero-Cazares, H., Garcia-Verdugo, J.M. & Quinones-Hinojosa, A. The generation of oligodendroglial cells is preserved in the rostral migratory stream during aging. Front Cell Neurosci 7, 147 (2013).

58. Fritze, J. et al. Loss of Cxcr5 alters neuroblast proliferation and migration in the aged brain. Stem Cells 38, 1175–1187 (2020).

59. Zhao, X. et al. 4D imaging analysis of the aging mouse neural stem cell niche reveals a dramatic loss of progenitor cell dynamism regulated by the RHO-ROCK pathway. Stem Cell Reports 17, 245–258 (2022).

60. Morante-Redolat, J.M. & Porlan, E. Neural Stem Cell Regulation by Adhesion Molecules Within the Subependymal Niche. Front Cell Dev Biol 7, 102 (2019).

61. Kazanis, I. et al. Quiescence and activation of stem and precursor cell populations in the subependymal zone of the mammalian brain are associated with distinct cellular and extracellular matrix signals. J Neurosci 30, 9771–81 (2010).

62. Kjell, J. et al. Defining the Adult Neural Stem Cell Niche Proteome Identifies Key Regulators of Adult Neurogenesis. Cell Stem Cell 26, 277–293 e8 (2020).

63. Luo, J., Shook, B.A., Daniels, S.B. & Conover, J.C. Subventricular zone-mediated ependyma repair in the adult mammalian brain. J Neurosci 28, 3804–13 (2008).

64. Boom, A. et al. Astrocytic calcium/zinc binding protein S100A6 over expression in Alzheimer’s disease and in PS1/APP transgenic mice models. Biochim Biophys Acta 1742, 161–8 (2004).

65. Hoyaux, D. et al. S100A6 overexpression within astrocytes associated with impaired axons from both ALS mouse model and human patients. J Neuropathol Exp Neurol 61, 736–44 (2002).

66. Gotz, M., Sirko, S., Beckers, J. & Irmler, M. Reactive astrocytes as neural stem or progenitor cells: In vivo lineage, In vitro potential, and Genome-wide expression analysis. Glia 63, 1452–68 (2015).

67. Liddelow, S.A. & Barres, B.A. Reactive Astrocytes: Production, Function, and Therapeutic Potential. Immunity 46, 957–967 (2017).

68. Buffo, A. et al. Origin and progeny of reactive gliosis: A source of multipotent cells in the injured brain. Proc Natl Acad Sci U S A 105, 3581–6 (2008).

69. Liddelow, S.A. et al. Neurotoxic reactive astrocytes are induced by activated microglia. Nature 541, 481–487 (2017).

70. Das, S., Li, Z., Noori, A., Hyman, B.T. & Serrano-Pozo, A. Meta-analysis of mouse transcriptomic studies supports a context-dependent astrocyte reaction in acute CNS injury versus neurodegeneration. J Neuroinflammation 17, 227 (2020).

71. Ahlenius, H., Visan, V., Kokaia, M., Lindvall, O. & Kokaia, Z. Neural stem and progenitor cells retain their potential for proliferation and differentiation into functional neurons despite lower number in aged brain. J Neurosci 29, 4408–19 (2009).

72. Morimatsu, M., Mekhdjian, A.H., Adhikari, A.S. & Dunn, A.R. Molecular tension sensors report forces generated by single integrin molecules in living cells. Nano Lett 13, 3985–9 (2013).

73. Chang, A.C. et al. Single Molecule Force Measurements in Living Cells Reveal a Minimally Tensioned Integrin State. ACS Nano 10, 10745–10752 (2016).

74. Kramer, A., Green, J., Pollard, J., Jr. & Tugendreich, S. Causal analysis approaches in Ingenuity Pathway Analysis. Bioinformatics 30, 523–30 (2014).

75. Suzuki, N., Hajicek, N. & Kozasa, T. Regulation and physiological functions of G12/13- mediated signaling pathways. Neurosignals 17, 55–70 (2009).

76. Christie, K.J., Turbic, A. & Turnley, A.M. Adult hippocampal neurogenesis, Rho kinase inhibition and enhancement of neuronal survival. Neuroscience 247, 75–83 (2013).

77. Emre, N. et al. The ROCK inhibitor Y-27632 improves recovery of human embryonic stem cells after fluorescence-activated cell sorting with multiple cell surface markers. PLoS One 5, e12148 (2010).

78. Uehata, M. et al. Calcium sensitization of smooth muscle mediated by a Rho-associated protein kinase in hypertension. Nature 389, 990–4 (1997).

79. Kim, J.E., Ryu, H.J., Kim, M.J. & Kang, T.C. LIM kinase-2 induces programmed necrotic neuronal death via dysfunction of DRP1-mediated mitochondrial fission. Cell Death Differ 21, 1036–49 (2014).

80. Leong, S.Y., Faux, C.H., Turbic, A., Dixon, K.J. & Turnley, A.M. The Rho kinase pathway regulates mouse adult neural precursor cell migration. Stem Cells 29, 332–43 (2011).

81. Peh, G.S. et al. The effects of Rho-associated kinase inhibitor Y-27632 on primary human corneal endothelial cells propagated using a dual media approach. Sci Rep 5, 9167 (2015).

82. Narumiya, S., Ishizaki, T. & Uehata, M. Use and properties of ROCK-specific inhibitor Y-27632. Methods Enzymol 325, 273–84 (2000).

83. Koester, J. et al. Niche stiffening compromises hair follicle stem cell potential during ageing by reducing bivalent promoter accessibility. Nat Cell Biol 23, 771–781 (2021).

84. Tumpel, S. & Rudolph, K.L. Quiescence: Good and Bad of Stem Cell Aging. Trends Cell Biol 29, 672–685 (2019).

85. Liu, L. et al. Chromatin modifications as determinants of muscle stem cell quiescence and chronological aging. Cell Rep 4, 189–204 (2013).

86. Schworer, S. et al. Epigenetic stress responses induce muscle stem-cell ageing by Hoxa9 developmental signals. Nature 540, 428–432 (2016).

87. Sun, D. et al. Epigenomic profiling of young and aged HSCs reveals concerted changes during aging that reinforce self-renewal. Cell Stem Cell 14, 673–88 (2014).

88. Mauffrey, P. et al. Progenitors from the central nervous system drive neurogenesis in cancer. Nature 569, 672–678 (2019).

89. Nascimento, M.A., Sorokin, L. & Coelho-Sampaio, T. Fractone Bulbs Derive from Ependymal Cells and Their Laminin Composition Influence the Stem Cell Niche in the Subventricular Zone. J Neurosci 38, 3880–3889 (2018).

90. Sato, Y. et al. Ventricular-subventricular zone fractones are speckled basement membranes that function as a neural stem cell niche. Mol Biol Cell 30, 56–68 (2019).

91. Porlan, E. et al. MT5-MMP regulates adult neural stem cell functional quiescence through the cleavage of N-cadherin. Nat Cell Biol 16, 629–38 (2014).

92. Kokovay, E. et al. VCAM1 is essential to maintain the structure of the SVZ niche and acts as an environmental sensor to regulate SVZ lineage progression. Cell Stem Cell 11, 220–30 (2012).

93. Segel, M. et al. Niche stiffness underlies the ageing of central nervous system progenitor cells. Nature 573, 130–134 (2019).

94. Li, C.X. et al. MicroRNA-21 preserves the fibrotic mechanical memory of mesenchymal stem cells. Nat Mater 16, 379–389 (2017).

95. Yang, C., Tibbitt, M.W., Basta, L. & Anseth, K.S. Mechanical memory and dosing influence stem cell fate. Nat Mater 13, 645–52 (2014).

96. Nasrollahi, S. et al. Past matrix stiffness primes epithelial cells and regulates their future collective migration through a mechanical memory. Biomaterials 146, 146–155 (2017).

97. Miroshnikova, Y.A., Nava, M.M. & Wickstrom, S.A. Emerging roles of mechanical forces in chromatin regulation. J Cell Sci 130, 2243–2250 (2017).

98. Nava, M.M. et al. Heterochromatin-Driven Nuclear Softening Protects the Genome against Mechanical Stress-Induced Damage. Cell 181, 800–817 e22 (2020).

99. Jones, D.L. et al. ZNF416 is a pivotal transcriptional regulator of fibroblast mechanoactivation. J Cell Biol 220(2021).

100. Goetsch, K.P., Snyman, C., Myburgh, K.H. & Niesler, C.U. ROCK-2 is associated with focal adhesion maturation during myoblast migration. J Cell Biochem 115, 1299–307 (2014).

101. Salhia, B. et al. Inhibition of Rho-kinase affects astrocytoma morphology, motility, and invasion through activation of Rac1. Cancer Res 65, 8792–800 (2005).

102. Chen, Y., Chou, W.C., Ding, Y.M. & Wu, Y.C. Caffeine inhibits migration in glioma cells through the ROCK-FAK pathway. Cell Physiol Biochem 33, 1888–98 (2014).

103. Fu, P.C. et al. The Rho-associated kinase inhibitors Y27632 and fasudil promote microglial migration in the spinal cord via the ERK signaling pathway. Neural Regen Res 13, 677–683 (2018).

104. Piltti, J., Varjosalo, M., Qu, C., Hayrinen, J. & Lammi, M.J. Rho-kinase inhibitor Y- 27632 increases cellular proliferation and migration in human foreskin fibroblast cells. Proteomics 15, 2953–65 (2015).

105. Rudolph, J. et al. The JAK inhibitor ruxolitinib impairs dendritic cell migration via off- target inhibition of ROCK. Leukemia 30, 2119–2123 (2016).

106. Srinivasan, S. et al. Blockade of ROCK inhibits migration of human primary keratinocytes and malignant epithelial skin cells by regulating actomyosin contractility. Sci Rep 9, 19930 (2019).

107. Dyberg, C. et al. Inhibition of Rho-Associated Kinase Suppresses Medulloblastoma Growth. Cancers (Basel) 12(2019).

108. Rolando, C. et al. Distinct roles of Nogo-a and Nogo receptor 1 in the homeostatic regulation of adult neural stem cell function and neuroblast migration. J Neurosci 32, 17788–99 (2012).

109. Feng, J.F. et al. Guided migration of neural stem cells derived from human embryonic stem cells by an electric field. Stem Cells 30, 349–55 (2012).

110. Galindo, L.T. et al. Chondroitin Sulfate Impairs Neural Stem Cell Migration Through ROCK Activation. Mol Neurobiol 55, 3185–3195 (2018).

111. Gengatharan, A. et al. Adult neural stem cell activation in mice is regulated by the day/night cycle and intracellular calcium dynamics. Cell 184, 709–722 e13 (2021).

112. Koch, J.C. et al. ROCK inhibition in models of neurodegeneration and its potential for clinical translation. Pharmacol Ther 189, 1–21 (2018).

113. Rikitake, Y. et al. Inhibition of Rho kinase (ROCK) leads to increased cerebral blood flow and stroke protection. Stroke 36, 2251–7 (2005).

114. Sladojevic, N., Yu, B. & Liao, J.K. ROCK as a therapeutic target for ischemic stroke. Expert Rev Neurother 17, 1167–1177 (2017).

## References

1. Zhuo, L. et al. Live astrocytes visualized by green fluorescent protein in transgenic mice. Dev Biol 187, 36–42 (1997).

2. Platt, R.J. et al. CRISPR-Cas9 knockin mice for genome editing and cancer modeling. Cell 159, 440–55 (2014).

3. Leeman, D.S. et al. Lysosome activation clears aggregates and enhances quiescent neural stem cell activation during aging. Science 359, 1277–1283 (2018).

4. Codega, P. et al. Prospective identification and purification of quiescent adult neural stem cells from their in vivo niche. Neuron 82, 545–59 (2014).

5. Dulken, B.W., Leeman, D.S., Boutet, S.C., Hebestreit, K. & Brunet, A. Single-Cell Transcriptomic Analysis Defines Heterogeneity and Transcriptional Dynamics in the Adult Neural Stem Cell Lineage. Cell Rep 18, 777–790 (2017).

6. Beckervordersandforth, R. et al. In vivo fate mapping and expression analysis reveals molecular hallmarks of prospectively isolated adult neural stem cells. Cell Stem Cell 7, 744–58 (2010).

7. Buenrostro, J.D., Giresi, P.G., Zaba, L.C., Chang, H.Y. & Greenleaf, W.J. Transposition of native chromatin for fast and sensitive epigenomic profiling of open chromatin, DNA- binding proteins and nucleosome position. Nat Methods 10, 1213–8 (2013).

8. Conover, J.C. & Shook, B.A. Aging of the subventricular zone neural stem cell niche. Aging Dis 2, 49–63 (2011).

9. Maslov, A.Y., Barone, T.A., Plunkett, R.J. & Pruitt, S.C. Neural stem cell detection, characterization, and age-related changes in the subventricular zone of mice. J Neurosci 24, 1726–33 (2004).

10. Kalamakis, G. et al. Quiescence Modulates Stem Cell Maintenance and Regenerative Capacity in the Aging Brain. Cell 176, 1407–1419 e14 (2019).

11. Dulken, B.W. et al. Single-cell analysis reveals T cell infiltration in old neurogenic niches. Nature 571, 205–210 (2019).

12. Martynoga, B. et al. Epigenomic enhancer annotation reveals a key role for NFIX in neural stem cell quiescence. Genes Dev 27, 1769–86 (2013).

13. Shen, L., Shao, N., Liu, X. & Nestler, E. ngs.plot: Quick mining and visualization of next-generation sequencing data by integrating genomic databases. BMC Genomics 15, 284 (2014).

14. Stark, R.B., G.D. DiffBind: differential binding analysis of ChIP-seq peak data. Bioconductor (2011).

15. Ross-Innes, C.S. et al. Differential oestrogen receptor binding is associated with clinical outcome in breast cancer. Nature 481, 389–93 (2012).

16. Yu, G., Wang, L.G. & He, Q.Y. ChIPseeker: an R/Bioconductor package for ChIP peak annotation, comparison and visualization. Bioinformatics 31, 2382–3 (2015).

17. Love, M.I., Huber, W. & Anders, S. Moderated estimation of fold change and dispersion for RNA-seq data with DESeq2. Genome Biol 15, 550 (2014).

18. Robinson, M.D., McCarthy, D.J. & Smyth, G.K. edgeR: a Bioconductor package for differential expression analysis of digital gene expression data. Bioinformatics 26, 139–40 (2010).

19. McCarthy, D.J., Chen, Y. & Smyth, G.K. Differential expression analysis of multifactor RNA-Seq experiments with respect to biological variation. Nucleic Acids Res 40, 4288–97 (2012).

20. Schep, A.N. et al. Structured nucleosome fingerprints enable high-resolution mapping of chromatin architecture within regulatory regions. Genome Res 25, 1757–70 (2015).

21. Chen, E.Y. et al. Enrichr: interactive and collaborative HTML5 gene list enrichment analysis tool. BMC Bioinformatics 14, 128 (2013).

22. Kuleshov, M.V. et al. Enrichr: a comprehensive gene set enrichment analysis web server 2016 update. Nucleic Acids Res 44, W90–7 (2016).

23. Avsec, Z. et al. Base-resolution models of transcription-factor binding reveal soft motif syntax. Nat Genet 53, 354–366 (2021).

24. Trevino, A.E. et al. Chromatin and gene-regulatory dynamics of the developing human cerebral cortex at single-cell resolution. Cell 184, 5053–5069 e23 (2021).

25. Lundberg, S. & Lee, S. A Unified Approach to Interpreting Model Predictions. *arxiv*, arXiv:1705.07874 (2017).

26. Shrikumar, A., Greenside, P. & Kundaje, A. Learning Important Features Through Propagating Activation Differences. in Proceedings of the 34th International Conference on Machine Learning Vol. 70 (eds Doina, P. & Yee Whye, T.) 3145–3153 (PMLR, Proceedings of Machine Learning Research, 2017).

27. Shrikumar, A. et al. Technical Note on Transcription Factor Motif Discovery from Importance Scores (TF-MoDISco) version 0.5.6.5. arXiv:1811.00416 (2018).

28. Kulakovskiy, I.V. et al. HOCOMOCO: towards a complete collection of transcription factor binding models for human and mouse via large-scale ChIP-Seq analysis. Nucleic Acids Res 46, D252–D259 (2018).

29. Gupta, S., Stamatoyannopoulos, J.A., Bailey, T.L. & Noble, W.S. Quantifying similarity between motifs. Genome Biol 8, R24 (2007).

30. Stuart, T. et al. Comprehensive Integration of Single-Cell Data. Cell 177, 1888–1902 e21 (2019).

31. Dominguez Conde, C. et al. Cross-tissue immune cell analysis reveals tissue-specific features in humans. Science 376, eabl5197 (2022).

32. Tabula Muris, C. A single-cell transcriptomic atlas characterizes ageing tissues in the mouse. Nature 583, 590–595 (2020).

33. Buckley, M.T. et al. Cell type-specific aging clocks to quantify aging and rejuvenation in regenerative regions of the brain. bioRxiv (2022).

34. Liu, L. et al. Exercise reprograms the inflammatory landscape of multiple stem cell compartments during mammalian aging. bioRxiv (2022).

35. Schindelin, J. et al. Fiji: an open-source platform for biological-image analysis. Nat Methods 9, 676–82 (2012).

36. Conant, D. et al. Inference of CRISPR Edits from Sanger Trace Data. CRISPR J 5, 123–130 (2022).

37. Morgens, D.W. et al. Genome-scale measurement of off-target activity using Cas9 toxicity in high-throughput screens. Nat Commun 8, 15178 (2017).

38. Luo, J., Shook, B.A., Daniels, S.B. & Conover, J.C. Subventricular zone-mediated ependyma repair in the adult mammalian brain. J Neurosci 28, 3804–13 (2008).

39. Luo, J., Daniels, S.B., Lennington, J.B., Notti, R.Q. & Conover, J.C. The aging neurogenic subventricular zone. Aging Cell 5, 139–52 (2006).

40. Liddelow, S.A. & Barres, B.A. Reactive Astrocytes: Production, Function, and Therapeutic Potential. Immunity 46, 957–967 (2017).

41. Das, S., Li, Z., Noori, A., Hyman, B.T. & Serrano-Pozo, A. Meta-analysis of mouse transcriptomic studies supports a context-dependent astrocyte reaction in acute CNS injury versus neurodegeneration. J Neuroinflammation 17, 227 (2020).

42. Kramer, A., Green, J., Pollard, J., Jr. & Tugendreich, S. Causal analysis approaches in Ingenuity Pathway Analysis. Bioinformatics 30, 523–30 (2014).

43. Chang, A.C. et al. Single Molecule Force Measurements in Living Cells Reveal a Minimally Tensioned Integrin State. ACS Nano 10, 10745–10752 (2016).

44. Tan, S.J. et al. Regulation and dynamics of force transmission at individual cell-matrix adhesion bonds. Sci Adv 6, eaax0317 (2020).

45. Morimatsu, M., Mekhdjian, A.H., Chang, A.C., Tan, S.J. & Dunn, A.R. Visualizing the interior architecture of focal adhesions with high-resolution traction maps. Nano Lett 15, 2220–8 (2015).

46. Edelstein, A.D. et al. Advanced methods of microscope control using muManager software. J Biol Methods 1(2014).

47. Leong, S.Y., Faux, C.H., Turbic, A., Dixon, K.J. & Turnley, A.M. The Rho kinase pathway regulates mouse adult neural precursor cell migration. Stem Cells 29, 332–43 (2011).

